# Mapping *Drosophila* insulin receptor structure to the regulation of aging through analysis of amino acid substitutions

**DOI:** 10.1101/2020.06.30.180505

**Authors:** Rochele Yamamoto, Michael Palmer, Helen Koski, Noelle Curtis-Joseph, Marc Tatar

**Affiliations:** Department of Ecology and Evolutionary Biology, Brown University, Providence, Rhode Island, USA; C4 Therapeutics, Watertown, MA, USA

**Author notes:** Correspondence to: Marc Tatar.

**Keywords:** insulin receptor, longevity, reproduction, *Drosophila*, transphosphorylation, kinase insert domain

## Abstract

Genetic manipulations of the *Drosophila* insulin/IGF signaling system slow aging, but it remains unknown how the insulin/IGF receptor acts to modulate lifespan or differentiate this control from that of growth, reproduction and metabolism. With homologous recombination we produced an allelic series of single amino acid substitutions in the fly insulin receptor (InR). Based on emerging biochemical and structural data, we map amino acid substitutions to receptor function to longevity and fecundity. We propose *InR* mutants generate bias in the process of asymmetric transphosphorylation when the receptor is activated. This induces specific kinase subdomains that modulate lifespan by additive processes, one involving survival costs of reproduction and the other involving reproduction-independent systems of longevity assurance. We identify a mutant in the kinase insert domain that robustly extends lifespan without affecting growth or reproduction, suggesting this element controls aging through unique mechanisms of longevity assurance.

## INTRODUCTION

Analysis of insulin-like signaling provides a powerful approach to study aging. Early work extended lifespan through mutations of the insulin-like receptors *C. elegans daf-2* and Drosophila *InR* (1-3). The homology of these receptors to human insulin receptor (IR) and insulin growth factor-1 receptor (IGF1R) encouraged we could address human aging if we understood how they extend lifespan (4, 5). Progress has been simultaneously impressive and frustrated. Studies identified many downstream outputs of insulin/IGF receptors to mediate aging, including transduction pathways, transcription factors, interactions with TOR, and post-translational regulation of cellular metabolism (6-12). These outputs modulate cellular mechanisms of homeostasis including autophagy, mitochondrial uncoupling, reactive oxygen species, genomic stability, collagen remodeling, lipogenesis and AMP kinase (13-24). Work with invertebrates uncovered how insulin/IGF signaling controls aging within cells and tissues as well as non-autonomously across tissues (25-28). Altered insulin/IGF likewise effects aging in mammals. With some variation in outcome, knockdown of IR, IGF1R and insulin receptor substrates (IRS1, IRS2) is associated with extended mouse lifespan and reduced functional aging (29-33). In humans, polymorphisms of IGF1R and the transcription factor FOXO3A are associated with successful aging and exceptional lifespan (34-36). Overall, insulin/IGF signaling integrates cells, tissue and physiology to control adult survival and lifespan.

The breadth of insulin/IGF function, however, also challenges our ability to understand how it affects aging (37-40). Insulin/IGF signaling simultaneously regulates many traits including growth, metabolism, cell proliferation, differentiation, Dauer/diapause, and reproduction. Furthermore, these traits are mediated by a single insulin-like receptor in *C elegans* and *Drosophila* while these invertebrates produce many insulin-like ligands (41-43). The insulin-like receptor is a molecular switch-board taking many incoming calls and routing each to distinct signaling destinations. Broadly, understanding such pleiotropy is a longstanding problem in receptor tyrosine kinase biology (44, 45). For instance, how does mammalian IR and IGFR differentially control metabolism relative to growth (46, 47), how does the fibroblast growth factor receptor (FGFR) modulate mitogenesis relative to glucose homeostasis (48), or how does the c-Kit receptor differentially activate progenitors of hematopoietic cells relative to mast cells (49)?

One approach to this problem studies insulin/IGF receptor amino acid variants in cultured cells, in polymorphic or mutant animals, and even in humans with inherited insulin and IGF resistance (50-54). Insight is developed by relating the amino acid substitutions to knowledge of receptor structure. This strategy was illustrated by work with *C. elegans daf-2* (55, 56). About two dozen *daf-2* variants extend lifespan, and variously also effect Dauer, fertility, and growth.

Classes of these traits were defined by where substitutions fell in the ectodomain relative to the kinase domain, but understanding how the substitutions control outcomes was limited by the known structural data of the time.

Structural analysis has advanced in recent years and we can now productively apply this approach with *Drosophila*. The *Drosophila* insulin-like receptor was initially identified in the lab of Rosen (57). Subsequently, EMS mutants screened over a deficiency produced alleles that were viable as trans-heterozygotes (58-60). Sequencing of the cytoplasmic portion of several alleles identified substitutions in the kinase domain (42). These genotypes variously reduced cell growth, fecundity and receptor kinase activity (42, 60). In early work with this receptor, aging was slowed by one EMS-generated mutant (*InR*^E19^) when heterozygous over a P-element insertion allele (5). Here, we characterize how other, archival single amino acid *InR* EMS mutants effect aging using the method of Quantitative Complementation Testing (61, 62). We validate and extend these results by independently generating the putative substitutions of the *InR* EMS alleles through homologous recombination gene replacement and evaluate lifespan, growth, development rate and fecundity. Finally, recent advances provide remarkable insights into the structure of activated human insulin receptors (63, 64). We combine these data with our genetic analysis to generate new ways to understand how the insulin-like receptor modulates aging.

We will propose four interpretations. 1) *Drosophila* InR mediates aging in two independent ways. One effects reproduction that carries associated adult survival costs, the other effects mechanisms of longevity assurance independent of reproduction. These outcomes may explain why a variety of longevity-extending manipulations reduce reproduction, but others do not: they impact different programs regulated by insulin/IGF signaling. 2) Reducing the *quantity* of insulin receptors has little effect on aging. Longevity mutants of the insulin receptor simultaneously change the *intensity and quality* of signaling. 3) The genetic effects of *InR* alleles may be understood through the model of tyrosine kinase receptor activation by asymmetric transphosphorylation (45, 64). In wildtype homodimeric receptors the direction of asymmetry is random. We propose particular *InR* mutants bias asymmetry whereby only the protomer from one allele of the InR heterodimer can be activated. In mutant heterozygotes, aging is slowed when the activated protomer carries an appropriate mutation. 4) Some mutants slow aging by altering the receptor’s specificity for substrate adaptor proteins. We identify a dominant substitution in the kinase insert domain that robustly increases lifespan yet paradoxically produces normal growth and robust fecundity in the lab setting. We hypothesize the kinase insert domain modulates a unique SH2 binding site that regulates mechanisms of longevity assurance that are independent of reproduction.

## RESULTS

### Demographic quantitative complementation

We used Quantitative Complementation Testing (QTC) to screen *InR* EMS-alleles (Frasch series) for their ability to slow aging. QTC measures allelic effects as the phenotypic difference of a tested allele complemented to a standard hypomorph or deficiency relative to when complemented to a standard wildtype allele (61, 62, 65-67). Figure 1 illustrates mortality for several *InR* alleles when complemented to *InR*^E19^ relative to TM3, *InR*^+^ *Sb* (Fig. 1, Table S1). Mortality rate increases exponentially for all cohorts after the ‘left-hand boundary’ (68). The natural wildtype *InR*^NC442^ is tested in each block *i* of demographic analysis to control for period effects (Table S2). Figure 1G, H (Block 3) illustrates modestly reduced mortality (*β* < 0) for *InR*^NC442^/ *InR*^E19^ relative to *InR*^NC442^/TM3, *InR*^+^ *Sb* – a pattern observed in all blocks. In contrast, mortality coefficients for EMS alleles ranged from -1.80 to 1.64 (block adjusted 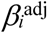) (Table S1). To infer when an allele slows aging, we asked when does an EMS allele effect mortality more than expected relative to segregating variation among a sample of wildtype alleles? Accordingly, we applied QCT to a collection of 18 wildtype *InR* alleles extracted from a natural population (NC series) and evaluated each EMS mutant relative to this distribution (*z*-test) (Table S1). Mortality coefficients for five EMS *InR* mutants did not significantly differ from wildtype alleles. One EMS mutant (*InR*^327^) increased mortally. *InR*^*74*^ and *InR*^*211*^ significantly reduced mortality in males and females. The *InR*^*353*^ allele reduced female mortality but this difference did not reach significance relative to the wildtype distribution. The transformation 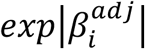 estimates fold change in mortality: *InR*^74^ and *InR*^211^ alleles reduced male and female mortality 2.6-to 6-fold, while *InR*^353^ reduced female mortality about 2-fold (Fig 2A, B).

**Figure 1.**
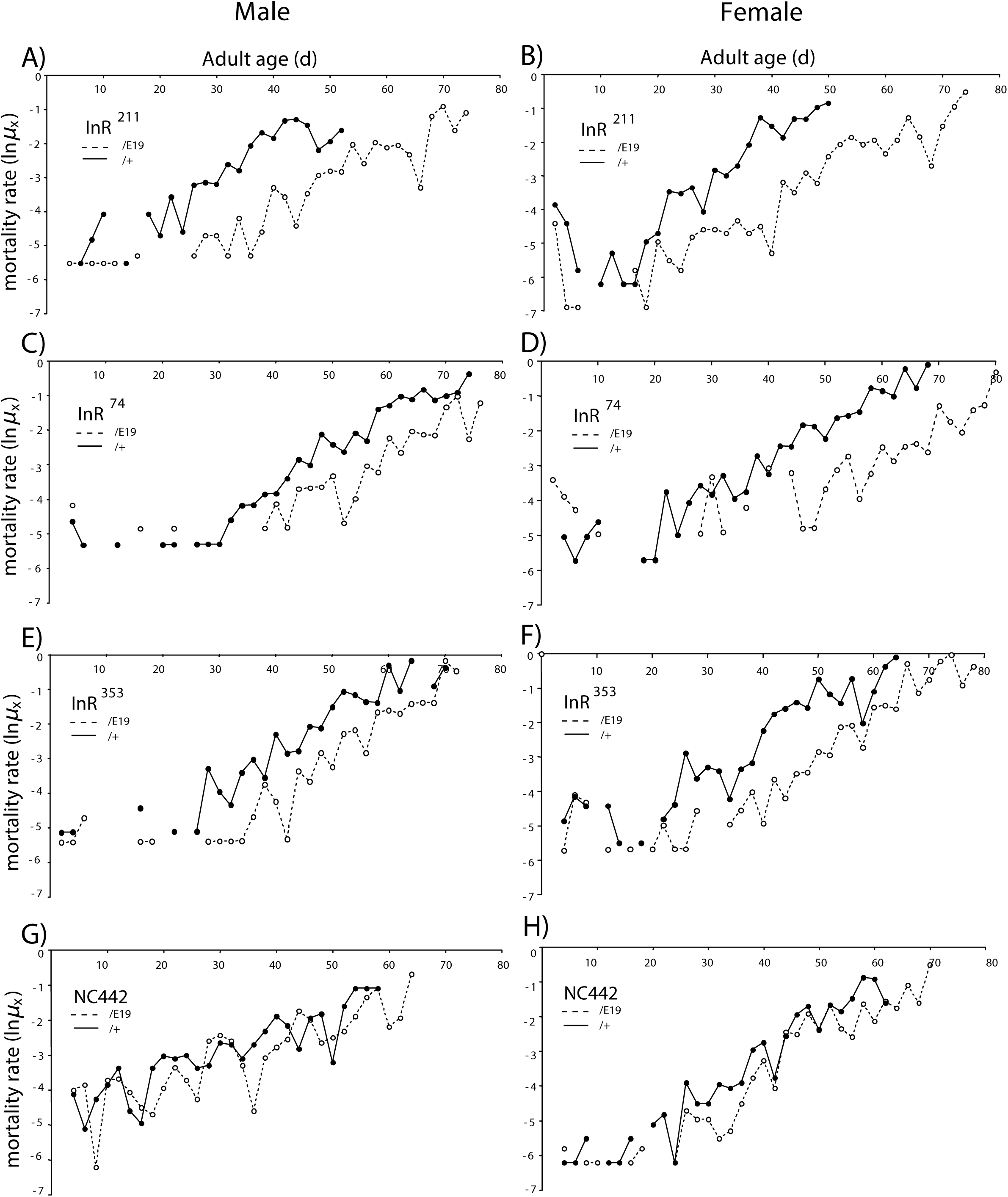
Mortality of EMS-generated and wildtype *InR* alleles in Quantitative Complementation Test. Log morality rate derived from observed age-specific mortality (*q*_*x*_), ln*μ*_*x*_ = ln(-ln(1-*q*_*x*_)). Block 3 data; *InR*^allele^/*InR*^E19^ (solid line), *InR*^allele^ /TM3, *InR*^+^ *Sb* (dashed line). Initial cohort size (N_0_) of genotypes combined. **A, B)** *InR*^211^: males (N_0_ = 484, *β* = -1.65, s.e. = 0.11); females (N_0_ = 712, *β* = -1.82, s.e. = 0.10). **C, D)** *InR*^74^: males (N_0_ = 337, *β* = -0.98, s.e. = 0.12); females (N_0_ = 442, *β* = -1.68, s.e. = 0.13). **E, F)** *InR*^353^: males (N_0_ = 394, *β* = -0.99, s.e. = 0.11); females (N_0_ = 560, *β* = -1.42, s.e. = 0.099). **G, H)** NC442 third chromosome (wildtype *InR*): males (N_0_= 443, *β* = -0.504, s.e. = 0.099); females (N_0_ = 571, *β* = -0.97, s.e. = 0.11). Cox proportional hazard analysis used to estimate mortality coefficient *β* for each allele as *InR*^allele^ /*InR*^E19^ relative to *allele*/TM3 *InR*^+^ *Sb. β* < 1.0 indicates *InR*^allele^/*InR*^E19^ has reduced mortality, p < 0.0001 in A-H (survival statistics in Table S1).

**Figure 2.**
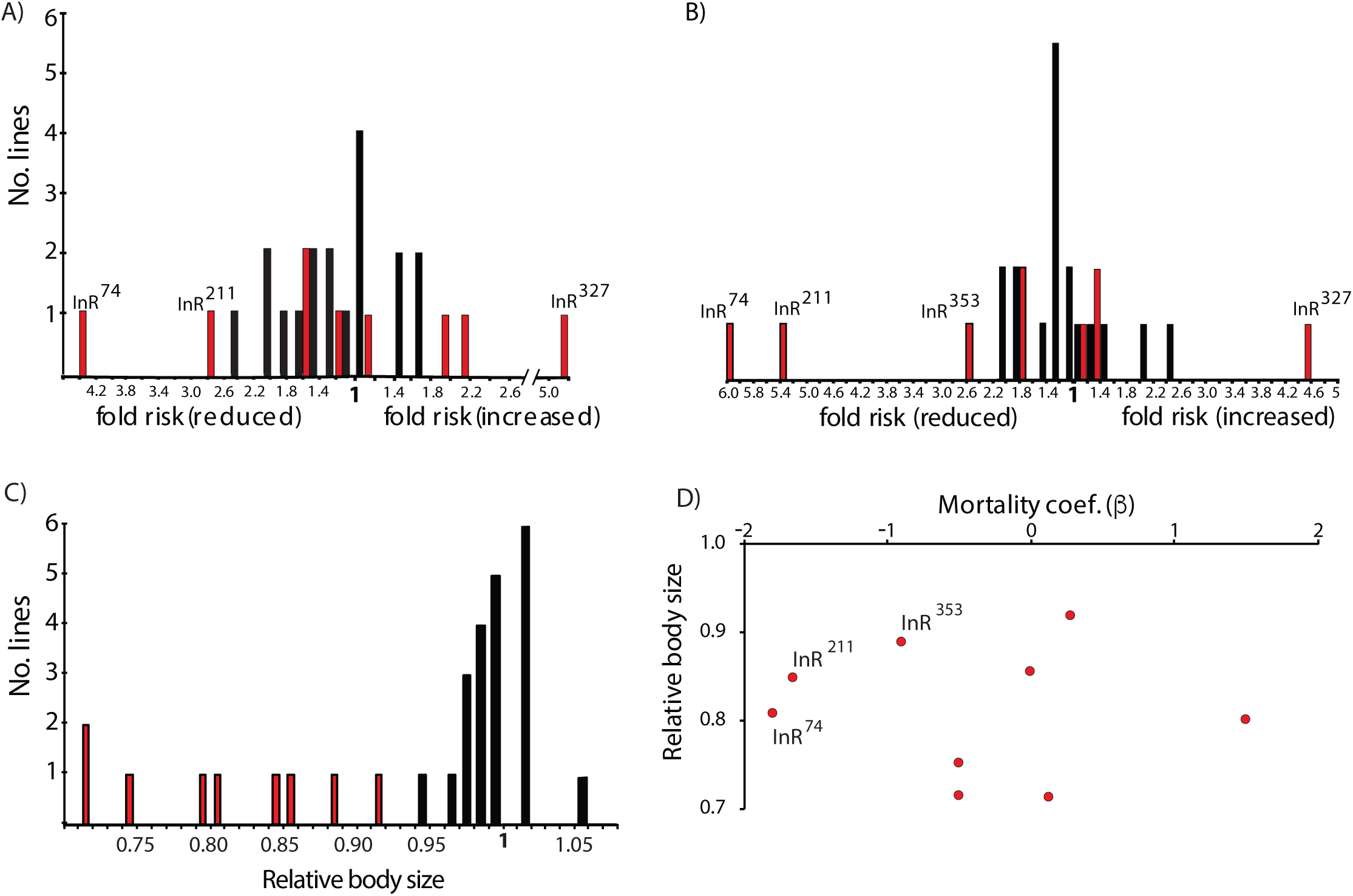
Traits from Quantitative Complementation Test of *InR* among EMS mutant and wildtype accessions. **A)** Males, **B)** Females; distribution of proportional hazard (fold risk of mortality) for nine EMS mutants (red bars) and 18 wildtype (NC series) accessions (black bars); estimated as exp(|β|). Labeled *InR* alleles have difference relative to distribution of wildtype hazard (Table S1). **C)** Distribution of relative body size (head capsule width) among mutant (red) and wildtype (black) females. All mutants smaller than the expected distribution of wildtype, *z*-test each allele, p < 0.0007. **D)** Absence of correlation between relative female mortality (*β*) and relative body size among *InR* alleles assessed in QCT (r^2^ = 0.08, p = 0.85).

### Body size quantitative complementation

To describe the allelic effects on body size, we quantified the ratio of each EMS allele complemented to *InR*^*E19*^ relative to TM3, *InR*^*+*^ *Sb*. All EMS *InR* alleles significantly reduced body size (female), with relative effects ranging from 0.9 to less than 0.75. (Fig 2C). Among the EMS *InR* alleles, relative body size does not associate with proportional change in mortality (Fig 2D).

### Homologous recombination alleles

Quantitative Complementation Testing has limitations: it only measures recessive allelic effects; it confounds epistatic interactions from cosegregating second site mutations; alleles are not derived from the same wildtype *InR* or background and may contain unrecognized polymorphisms; and allelic effects are relative rather than absolute and in each case these relative measures include deleterious effects of the TM3 haplotype.

To resolve these issues we generated new, single amino acid substitutions from a common *InR* in a universal background, focusing on alleles that potentially slowed aging in QTC, or in one case an allele that did not reduce mortality but produced small adults (*InR*^*246*^). We used published and unpublished data on the putative substitution sites (Fig 3, S1) to produce new alleles with single amino acid substitutions by ends out homologous recombination, designated as *InR*^*E19*(HR)^, *InR*^*74*(HR)^, *InR*^*211*(HR)^, *InR*^*246*(HR)^, and *InR*^*353*(HR)^, along with a wildtype control (*InR*^*+*(HR)^), a double mutant allele (*InR*^*E19;74*(HR)^), and a null *InR* allele (*InR*^*null*(HR)^). Except for the allele *InR*^*E19*(HR)^ which resides in the Fibronectin III-1 ectodomain, the remaining substitutions are located in the intracellular kinase domain (Fig 3).

**Figure 3.**
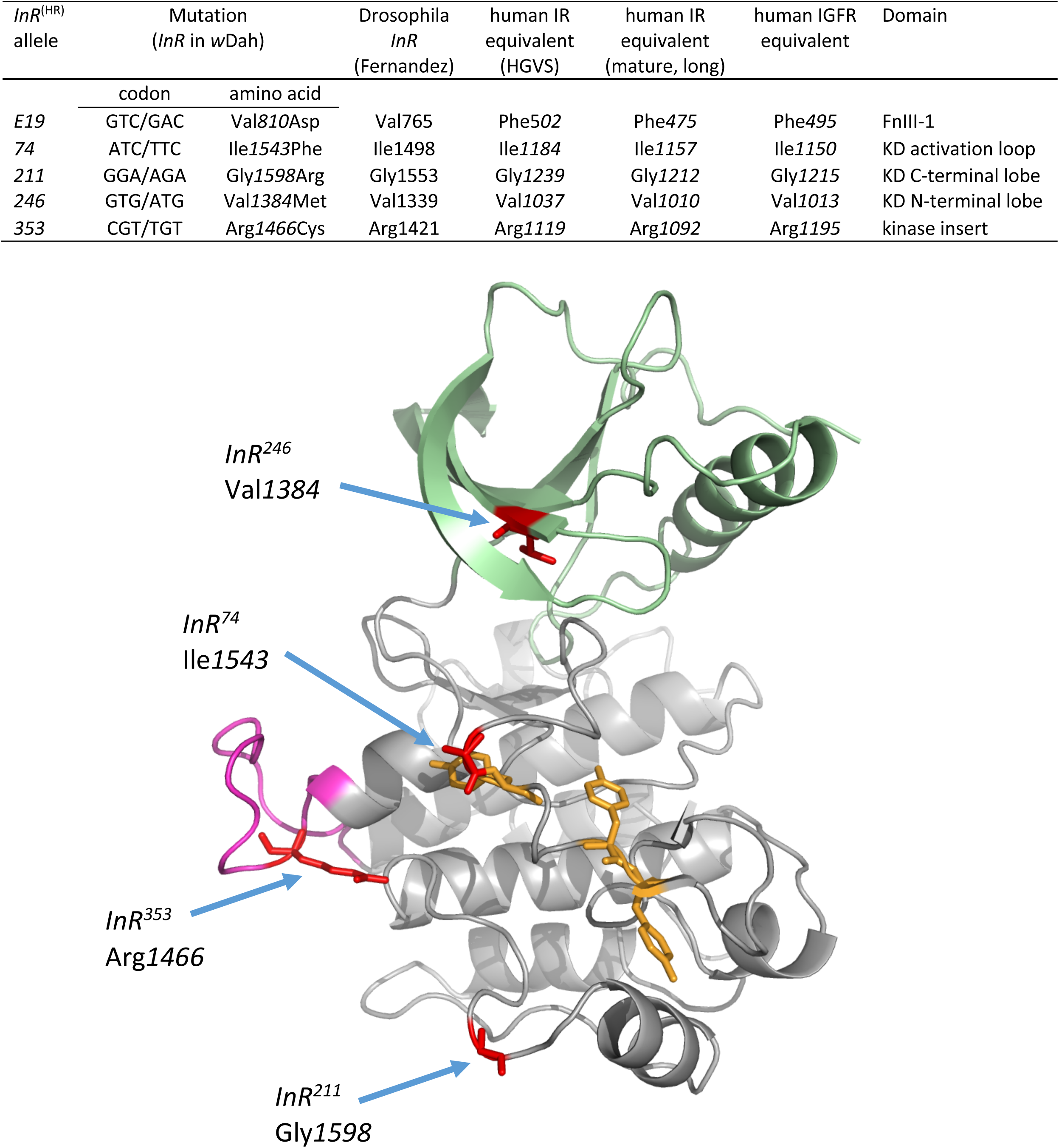
Sites, nomenclature and structure site of InR alleles generated by homologous recombination. Amino acids numbered from the translation initiation site of *w*Dah progenitor *InR* wildtype used in homologous recombination (GenBank accession MT_563159; Supplemental Figs S1-3). This TIS is 45 additional N-terminal amino acids from the TIS reported in the Fernandez (59). Human insulin receptor sequence is numbered following nomenclature of the Human Genome Variation Society (HGVS; http://www.hgvs.org/rec.html) and based on the mature, long-form type IR cDNA (105). The human IGF1R amino acid sequence is based on NCBI Reference Sequence: NP_000866.1. KD: kinase domain. FnIII: extra-cellular fibronectin domain III. Ribbon model of kinase domain (based on hIR), indicating conserved sites of amino acid substitutions (red). N-terminal lobe: green; C-terminal lobe: grey; kinase insert domain: magenta; Tyrosine residues of the activation loop: yellow.

We evaluated viability, eclosion time, and adult size for all allele combinations (Fig 4A, B, Fig S2). *InR*^*+*(HR)^ complemented every mutant to produce normal phenotypes, while all mutants were homozygous lethal except for some escapes in *InR*^*E19*(HR)^/*InR*^*E19*(HR)^. All kinase domain mutants complemented *InR*^*E19*(HR)^, albeit with some delay in development. Several kinase domain mutants were viable as transheterozygotes: *InR*^*74*(HR)^/*InR*^*353*(HR)^ and *InR*^*74*(HR)^/*InR*^*211*(HR)^. Adult size was reduced in all viable mutant genotypes, except *InR*^*+*(HR)^/*InR*^*353*(HR)^ which were as large as wildtype.

**Figure 4.**
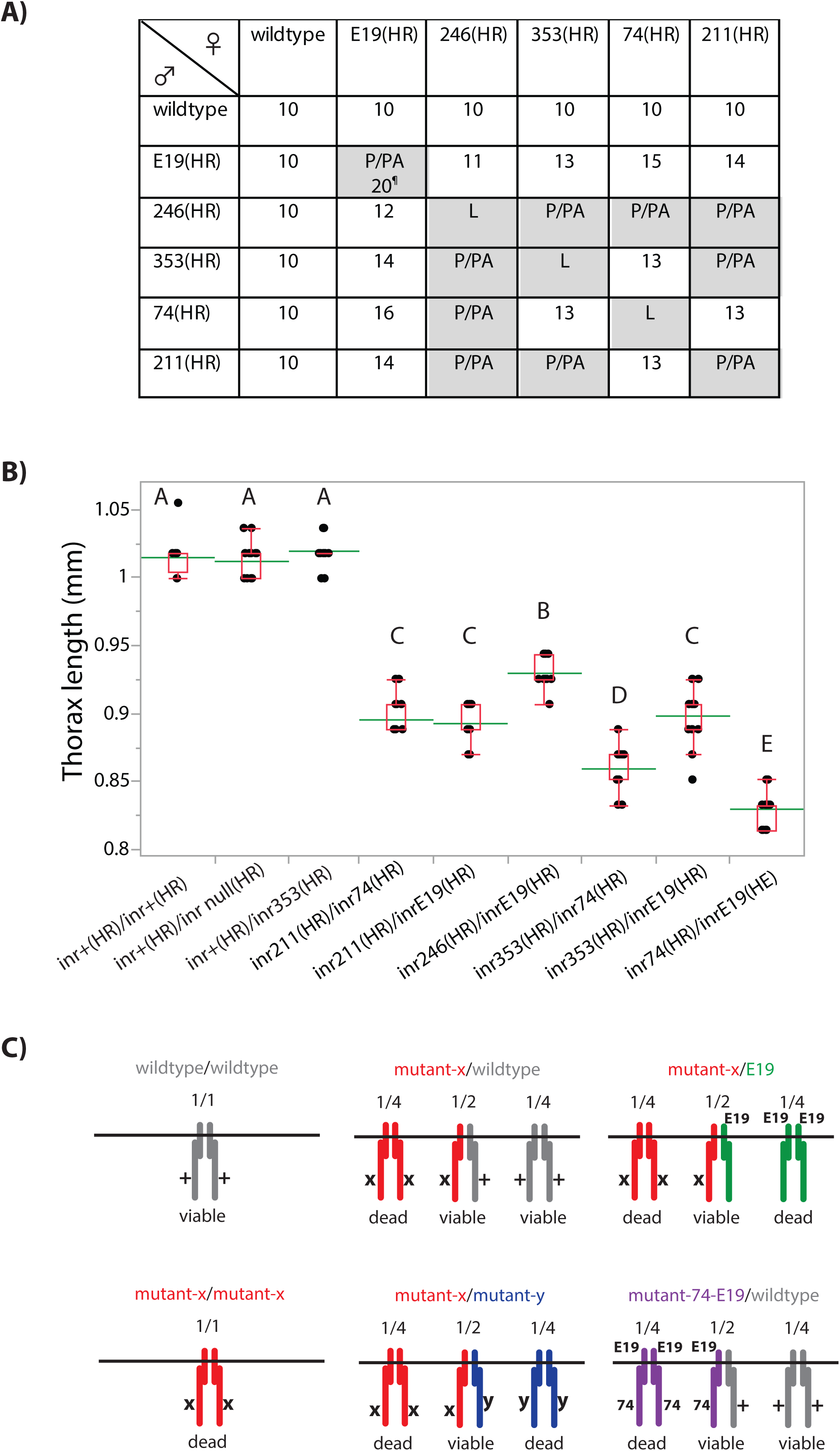
Characteristics of homologous recombination *InR*^(HR)^ alleles. **A)** Development time (days to first eclosion), or stage of lethality for all allele combinations, reciprocal crosses. Lethality at first or second stage larvae (L), late pupae (P) or pharate adult (PA). ¶: *InR*^E19(HR)^/*InR*^E19(HR)^ produce few minute adults at 20 days, but otherwise pupal/pharate lethal. **B)** Adult size (thorax length, mm) of *InR*^(HR)^ alleles across genotypes. Means, std. dev. and range shown, N=20 females each genotype. Means not connected by same letter are significantly different (one-way ANOVA, Tukey-Kramer HSD comparison, p < 0.05). **C)** *InR* genotypes produce combinations of constitutive dimers within expressing cells, with indicated ratios if protomers pair randomly. Wildtype homozygotes produce a full complement of functional dimers, while homozygote mutants generate (largely) nonfunctional (dead) dimers. In mutant/wildtype heterozygotes, homodimers are nonfunctional but their heterodimers are inferred to be functional, along with their wildtype homodimers. In mutant/*InR*^E19(HR)^ heterozygotes, only heterodimeric receptors are viable. Viable trans-heterozygotes (mutant-x/mutant-y) must produce viable heterodimers. The double mutant *InR*^cis-74(HR),E19(HR)^ when heterozygote with wildtype is inferred to produce functional heterodimers along with normal wildtype homodimers.

### Demography of hemizygote InR^+(HR)^/InR^null(HR)^

Insulin receptors are preformed dimers assembled with protomers from both alleles (Fig 4C). Homodimeric receptors of *InR*^(HR)^ mutants appear to be nonfunctional because homozygotes of each mutant are inviable. Heterozygotes between two different mutants must produce functional heterodimers because these genotypes are viable, but these heterozygotes will also generate fewer functional dimers per cell. This raises the possibility that *InR* mutant heterozygotes extend longevity because their cells contain fewer functional receptors. To address this issue, we measured the survival of the hemizygote *InR*^*+*(HR)^*/InR*^*null*(HR)^. Hemizygotes produced 26% less *InR* mRNA and increased life expectancy between 2 to 4 days when assessed across replicate trials, but only significantly so in Trial 2 (Fig 5, Table S3). We use this 2 to 4-day difference as a benchmark to infer when an *InR* genotype extends longevity more than expected from quantitative loss of functional InR receptor dimers.

**Figure 5.**
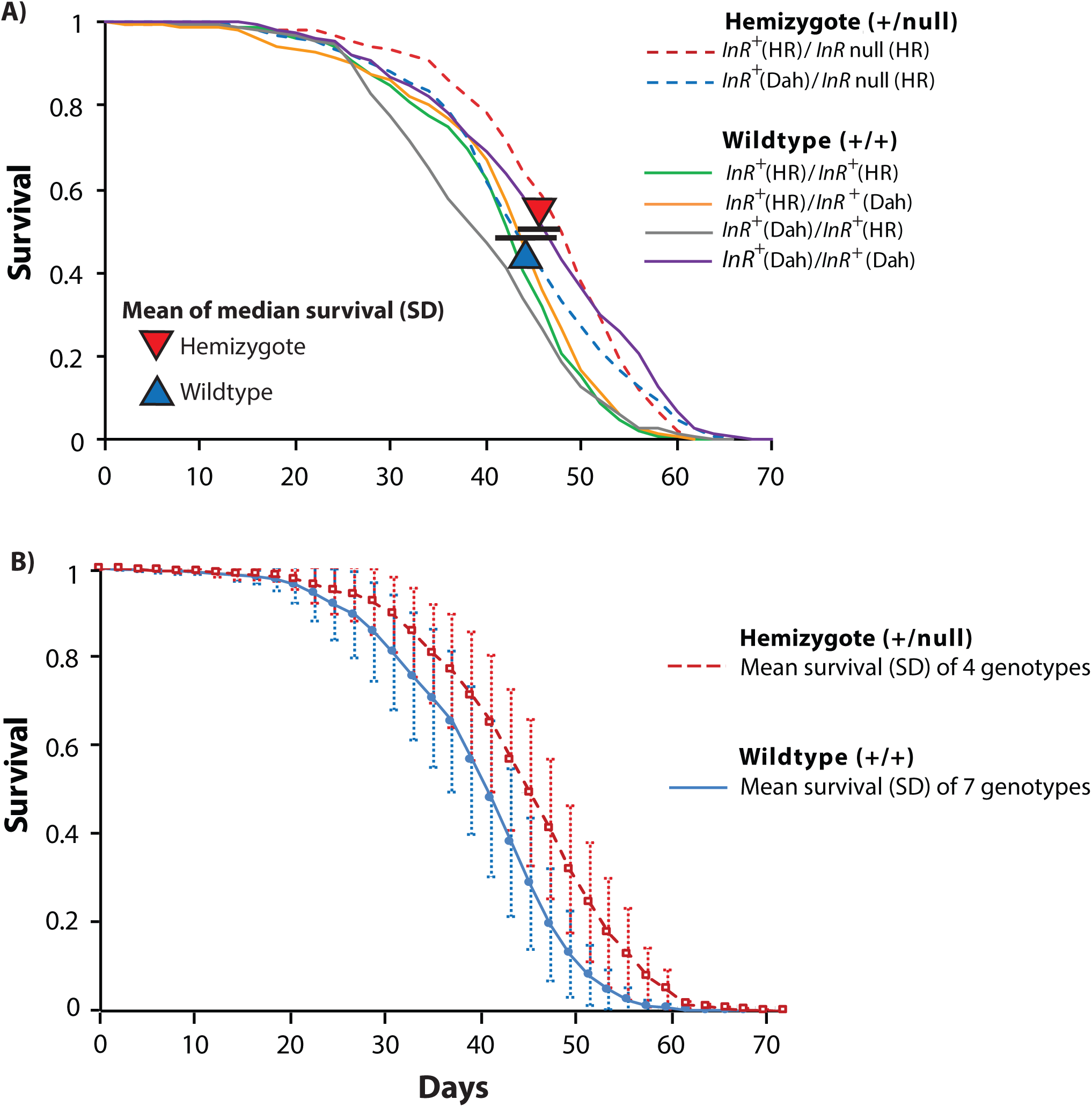
Impact of *InR* hemizygosity upon adult survival. **A)** Trial 1. Two hemizygote genotypes using one *InR*^null(HR)^, and wildtypes *InR*^+(HR)^ and *w*Dah (*InR*^+Dah^), compared to four homozygous wildtype genotypes generated from reciprocal, pairwise combinations of wildtype alleles. Triangles mark among cohort average of median lifespan within genotypic class, with standard deviation (bars). Hemizygotes are 2 days longer lived (among genotype-class median survival); derives from lower proportional hazard mortality of hemizygote (*β* = -0.14, likelihood ratio genotype class *χ* ^2^ = 36.4, p < 0.0001, DF = 1, N_0_ = 2126). **B)** Trial 2. Four hemizygote genotypes (combinations of two null and three wildtype) compared to seven homozygous wildtype (combinations of three wildtypes). Plotted: genotype-class survival with standard deviation. Hemizygotes are 4 days longer lived (among genotype-class median survival); derives from lower proportional hazard mortality of hemizygote (*β* = -0.25, likelihood ratio genotype class *χ*^2^ = 275.1, p < 0.0001, DF = 1, N_0_ = 5002).

### *Demography of mutant InR^(HR)^* alleles

The heterozygote *InR*^*E19(HR)*^ /*InR*^*+(HR)*^ does not extend lifespan (Fig 6A): average median life expectancy among replicate accessions was 43.3 d, compared to 44 d for wildtype (Table S4). Likewise, the intracellular kinase domain alleles *InR*^*74*(HR)^, *InR*^*211*(HR)^ and *InR*^*246*(HR)^ when complemented by *InR*^*+(HR)*^ did not reduce mortality any more than *InR* hemizygotes (Fig 6B, 6C, Table S4). In contrast, in replicated trials with independent accessions, *InR*^+^/ *InR*^*353*(HR)^ increased life expectancy by 10 to 16d by decreasing mortality ∼4-fold (Fig 6C, D) (Table S4).

**Figure 6.**
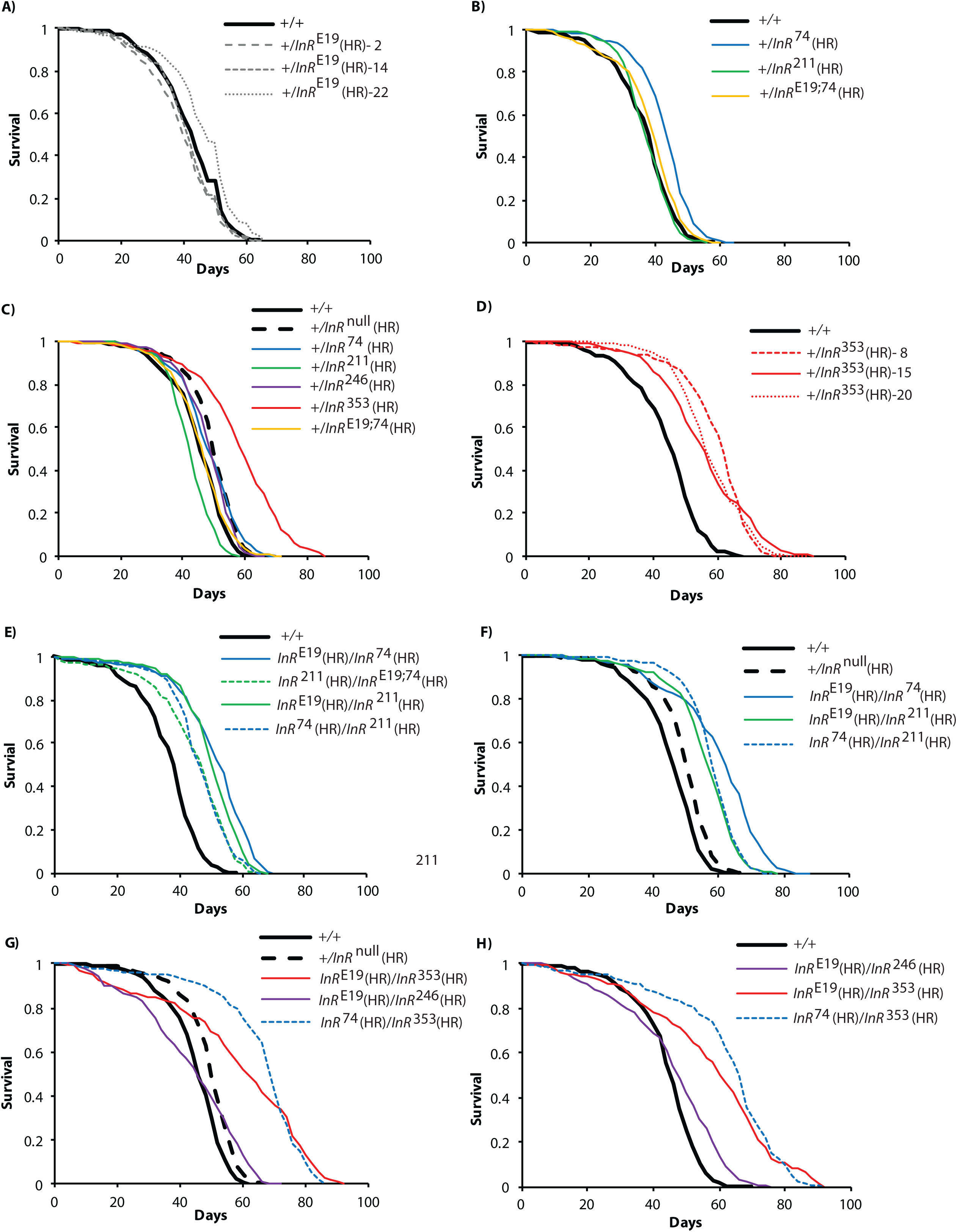
Survival of homologous recombination InR alleles. Females. Plots represent independent trials except for contemporaneous data in 6B and 6E, and in 6C, 6F and 6G. Wildtype and null in all trials were homologous recombinant *InR*^+(HR)29B^ and *InR*^null(HR)29B^ respectively. **A)** Wildtype heterozygotes with three accessions of ectodomain *InR*^E19(HR)^. **B)** Wildtype heterozygotes with mutants of the kinase domain A-loop and C-terminal lobe. **C)** Wildtype heterozygote and hemizygote with mutants of kinase domain N-terminal lobe, A-loop, KID and C-terminal lobe. **D)** Wildtype heterozygotes with the *InR*^353(HR)^ mutant of the kinase domain KID, three accessions. **E)** Heterozygote mutant genotypes with ectodomain *InR*^E19(HR)^ and kinase domain mutants *InR*^74(HR)^ and *InR*^211(HR)^. **F)** Heterozygote mutant genotypes with ectodomain *InR*^E19(HR)^ and kinase domain mutants *InR*^74(HR)^ and *InR*^211(HR)^ (independent trial). **G)** Heterozygote mutant genotypes with ectodomain *InR*^E19(HR)^, A-loop *InR*^74(HR)^, N-terminal lobe *InR*^246(HR)^ and KID *InR*^353(HR)^. **H)** Heterozygote mutant genotypes with ectodomain *InR*^E19(HR)^, A-loop *InR*^74(HR)^, N-terminal lobe *InR*^246(HR)^ and KID *InR*^353(HR)^ (independent trial). Lifetable summaries and proportional hazard statistics in Supplemental Table S4.

In QTC, kinase domain alleles reduced mortality when complemented to *InR*^*E19*^ relative to when complemented to TM3, *InR*^+^ *Sb*. Here we estimate the absolute allelic effect of each substitution using the *InR* homologous recombination alleles. Across replicate trials, the genotypes *InR*^*74*(HR)^/*InR*^*E19*(HR)^ and *InR*^*211*(HR)^/*InR*^*E19*(HR)^ extend lifespan 6-14 days (Fig 6E, F, Table S4). In a result not available from QTC, the heterozygote of kinase domain mutations *InR*^*74*(HR)^/*InR*^*211*(HR)^ also extended lifespan (Fig 6E, F). In every case, longevity benefits exceeded that of wildtype hemizygotes (Table S4).

Because the *trans*-heterozygote *InR*^*E19*(HR)^ and *InR*^*74*(HR)^ slows aging, we asked if these alleles could extend lifespan as a *cis*-double mutant, where both substitutions are on the same protomer. They do not: *InR*^*E19,74*(HR)^ had little effect on survival when heterozygous with wildtype *InR*^*+(HR)*^ (Fig 6B). We likewise tested *InR*^*E19,74*(HR)^ when *tran*s-heterozygous with *InR*^*211*(HR)^, and here adults were long-lived but no more so than *InR*^*74*(HR)^/*InR*^*211*(HR)^ (Fig 6E, Table S4). Among these mutant alleles, longevity is only extended when substitutions occur on opposing protomers.

In QCT, the *InR*^*246*^ allele did not slow aging more than expected relative to a sample of wildtype alleles, although it did reduce body size. We confirm these observations with *InR*^*246*(HR)^/*InR*^*E19*(HR)^: adults are small (Fig 4) but not long-lived (Fig 6G, H). We note, however, the survival curve of *InR*^*246*(HR)^/*InR*^*E19*(HR)^ crosses over that of wildtype, suggesting the mutant cohort has high age-independent mortality potentially coupled with reduced demographic aging.

As noted, *InR*^*+*(HR)^/*InR*^*353*(HR)^ extends lifespan relative to wildtype. Remarkably, *InR*^*353*(HR)^ further slows aging when complemented with *InR*^*E19*(HR)^ (14 to 15 days) and even more so when complemented with *InR*^*74*(HR)^ (21 to 22 days from 6-fold decrease in mortality risk) (Fig 6G, H, Table S4).

Overall, the homologous recombination alleles validate inferences from QCT but now measured as absolute effects. We also describe a new heteroallelic genotype that extend lifespan, and document an exceptional longevity benefit conferred by *InR*^*353*(HR)^ that was obscured in QCT because this allele is dominant.

### Fecundity

Drosophila insulin signaling modulates fecundity, and survival is a well-known cost of reproduction (69). We therefore determined how the *InR* alleles effect egg production (Fig 7A, B). Wildtype, hemizygotes and *InR*^*E19*(HR)^/*InR*^*246*(HR)^ have similar egg production. Mutant heterozygotes reduce fecundity. In contrast, *InR*^*+*(HR)^/*InR*^*353*(HR)^ females produce more eggs than wildtype. This exceptional fecundity could arise because these females have more ovarioles or produce more eggs per ovariole. Accordingly, we measured the number of ovarioles for all genotypes and regressed this trait against daily egg production. Fecundity associates with ovariole number, but ovarioles from genotypes with high fecundity also produces more eggs per day because the observed slope < 1.0 implies fecundity increases faster than the number of ovarioles (Fig 7C). We therefore regressed lifespan against ovariole egg production (eggs/day/ovariole) across genotypes, treating the *InR*^*353*(HR)^ allele as a covariate (Fig 7D). A striking pattern emerges: egg production per ovariole is negatively associated with life expectancy (although, N=8, p = 0.06), but independent of egg production, genotypes with one *InR*^*353*(HR)^ allele have ∼10 to 13 days greater life expectancy (p = 0.013). The *InR*^*353*(HR)^ allele confers an additive effect upon longevity assurance that is independent of reproductive trade-offs.

**Figure 7.**
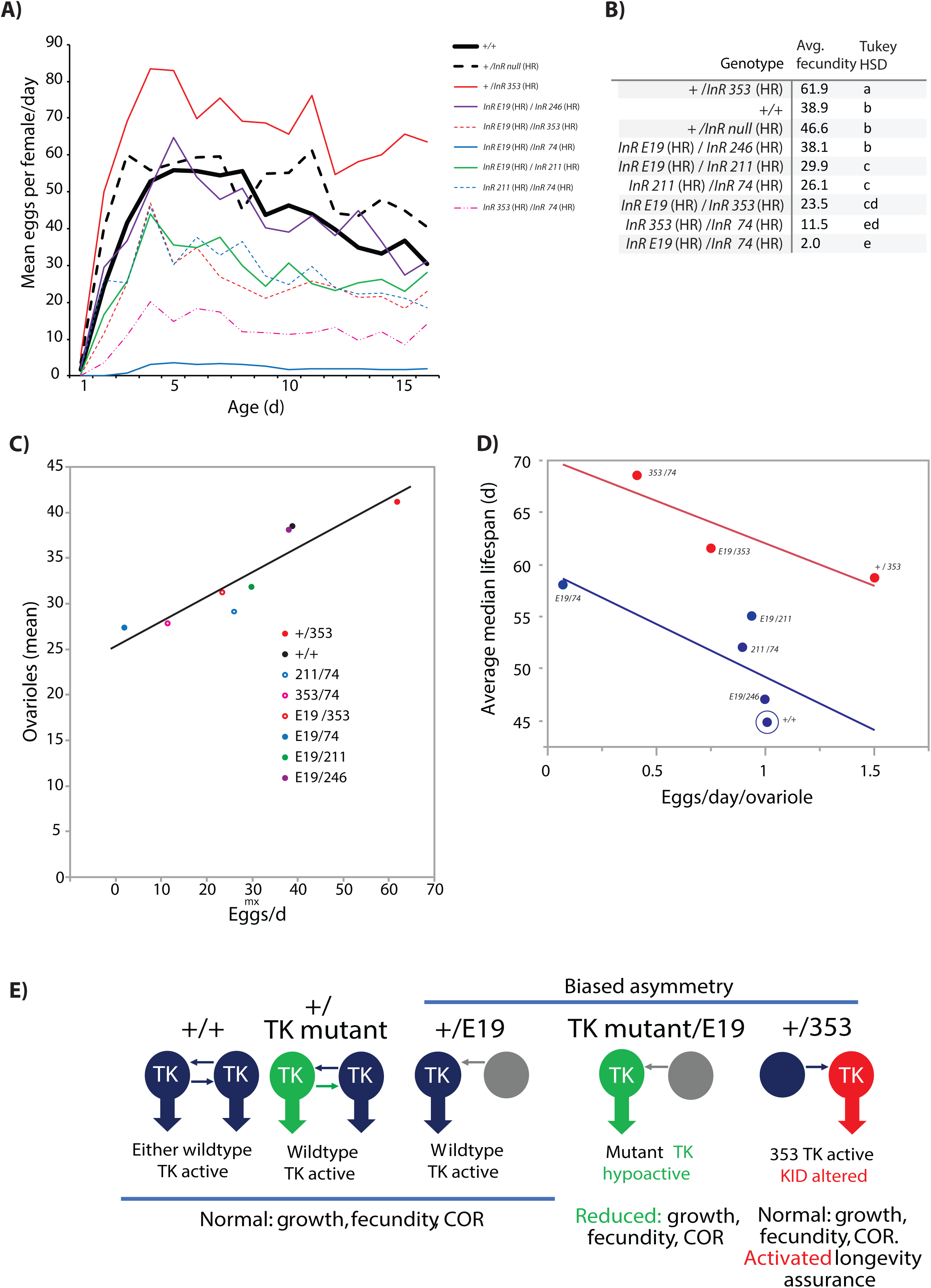
Fecundity, ovarioles and visual model. **A)** Among female mean eggs laid from eclosion through age 15d for mutant heterozygotes, wildtype and hemizygote. **B)** Average daily fecundity. Eggs per day averaged over 16 days. ANOVA with Tukey HSD post hoc analysis: genotypes with the same letter show no significant difference in average egg production. **C)** Regression of mean ovariole number (total from both ovarioles) upon average fecundity among genotypes. Least squares linear regression: R^2^= 0.85, *m* = 0.27< 1.0 *F*= 34.1, *p* < 0.001. **D)** Relationship between lifespan and rate of ovariole egg production among genotypes treating InR353 as a covariate. Average median lifespan is the average of median lifespans observed for each genotype among all its independent cohorts and accessions. Eggs/day/ovariole is estimated from average daily fecundity/mean ovariole number of each genotype. Parameters by ANCOVA: *β*_(fecundity/ovariole)_ = -9.15, *t* = -2.58, *p* = 0.061; *β* _(InR353)_ = 6.23, *t* = 4.25, p = 0.013; *β* _(fecundity/ovariole x InR353)_ = 1.01, *t* = 0.28, *p* = 0.79). **E)** Model for biased asymmetry modulates aging through InR domain specific function. Wildtype protomers (blue) each have the potential to asymmetrically act as either enzyme or substrate, activating catalysis by either tyrosine kinase (TK), producing wildtype phenotypes through the activation of just one TK. Protomers with mutation in tyrosine kinase activity (green) can act as substrate or enzyme protomers; when heterozygous with a wildtype protomer, enough dimers produce wildtype TK activation for normal phenotypes. The *InR*^E19(HR)^ protomer (grey) can only act as enzyme and transphosphorylate its partner protomer (biased asymmetry). When this is a wildtype protomer, the TK is active and phenotypes are normal. When the partner protomer has a hypomorphic mutation in the TK, this mutant TK reduces growth and reproduction, relieving survival costs of reproduction (COR) to extend lifespan. The 353 protomer only acts as substrate (biased asymmetry) but retains normal TK catalytic activity when transphosphorylated, and thus normal growth and reproduction. The protomer alters its kinase insert domain (KID) function, which induces a program of longevity assurance independent of reproduction.

## DISCUSSION

The impact of InR on longevity was originally studied with EMS-generated alleles heterozygous with a p-element *InR* mutant (3, 59). In particular, *InR*^*E19*^/ P{PZ}*InR*^*05545*^ extended lifespan and this genotype reduced InR tyrosine kinase activity (3). *InR* mutants are generally said to slow aging because they diminish insulin signaling, but this explanation is incomplete because many *InR* mutants reduce growth and reproduction without benefits to survival (3). Rather, InR may regulate aging through signals that are distinct from how it effects other traits. Here we explore this idea based on how *Drosophila InR* alleles impact aging, growth and reproduction.

### Alleles in the quantitative complementation test

We used quantitative complementation testing (QCT) to identify EMS *InR* mutants that reduced adult mortality. Among alleles complemented to *InR*^*E19*^ relative to TM3, *InR*^+^ *Sb*, three alleles reduced relative mortality more than expected from a sample of naturally occurring wildtype alleles. We followed the same approach to quantify how mutant alleles effect growth. Every allele significantly reduced adult size, and there was no association between relative size and relative mortality among genotypes.

QCT has caveats (62, 67). Our archival *InR* mutants were not generated on a common third chromosome. Unknown second site mutations could affect traits we attribute to *InR* and we cannot rule out the impact of epistatic loci that segregate with *InR*^*+*^ upon the TM3*Sb* balancer chromosome. We only estimate recessive effects of the mutants as they complement *InR*^*E19*^ relative to when complement TM3, *InR*^*+*^ *Sb*. We cannot evaluate other mutant heterozygote combinations. The causal substitutions within *InR* were not actually known, and unknown polymorphisms within each *InR* locus might exist in the stocks. Finally, QCT measures relative rather than absolute allelic effects, and the balancer haplotype itself appears to be somewhat deleterious.

To address these issues, we reconstituted the putative *InR* substitutions using ends-out homologous recombination. These *de novo* alleles provide new, well-defined substitutions derived from a common wildtype progenitor in a controlled genetic background. We produced four homologous recombination (HR) alleles of the Frasch series (*InR74*^(HR)^, *InR*^*211*(HR)^, *InR*^*246*(HR)^, *InR*^*353*(HR)^), an *InR*^*E19*(HR)^, an HR-derived wildtype allele, a null allele, and double mutant with the *E19* and *74* substitutions upon the same receptor protomer (*InR*^*E19,74*(HR)^). Our first question with these new alleles was, how does the *quantity* of InR affect aging?

### InR gene dosage in longevity assurance

InR, like human IR and IGFR, is a symmetric dimeric transmembrane receptor. Flies with two wildtype alleles generate two normal protomers and all dimeric receptors are potentially functional. Heterozygotes, on the other hand, may produce fewer viable receptors within each cell – mutant-mutant and mutant-wildtype dimers may not be functional. In insulin resistant humans with one mutant IR allele, mutant-wildtype and mutant-mutant dimers are reported to lack tyrosine kinase activity in cell-based assays; patients survive with limited wildtype-wildtype dimers (70). Accordingly, we tested whether *Drosophila* aging is slowed in *InR* heterozygotes because cells have less functional receptors. This does not seem to be the case for two reasons.

First, we find compound heterozygotes (*InR*^74(HR)^/*InR*^211(HR)^ and *InR*^74(HR)^/*InR*^353(HR)^) that are long-lived, despite the fact that each allele is homozygous lethal because their homodimer receptors are presumed to be nonfunctional. Second, the hemizygote *InR*^+(HR)^/*InR*^null(HR)^ slows aging by only 2-4 days. To the extent hemizygotes have fewer wildtype functional InR within cells, loss of receptor quantity has little effect on longevity. These data provide a benchmark: genotypes that extend longevity more than a few days may alter the *quality* of receptor signaling, not just the number of wildtype-dimer receptors. We seek potential qualitative mechanisms by relating the structure of human IR to single amino acid substitutions engineered into *Drosophila* InR.

### InR^E19^: extracellular fibronectin domain

*InR*^*+*^*/InR*^*E19*(HR)^ produces normal phenotypes while *InR*^*E19*(HR)^*/InR*^*E19*(HR)^ is lethal. *InR*^*E19*(HR)^ complemented with the alleles *InR*^74(HR)^, *InR*^353(HR)^, and *InR*^*211*(HR)^ is long-lived, yet *InR*^*E19*(HR)^ is not required to slow aging because some genotypes without this allele are also long-lived. How does *InR*^*E19*(HR)^ slow aging?

The *InR*^*E19*^ substitution Val*810*Asp occurs in a linker sequence between the L2 and FnIII-1 ectodomains, corresponding to Phe*475* of human IR (Ebina) and to Phe*495* of human IGF-1R (Figure 3, Suppl fig 3). Recent cryo-EM analyses show insulin induces a hinge motion in this linker to swing the fibronectin domains inward, bringing together the intracellular domains of each protomer (63, 71, 72). This proximity permits asymmetric kinase transphosphorylation, a non-reciprocal process whereby the kinase domain of one protomer acts as enzyme to phosphorylate the opposing (substrate) activation loop (A-loop) (45, 64, 73). Phosphorylation of the substrate A-loop releases the catalytic potential of its corresponding kinase and the dimer has full function derived from activation of this one kinase domain.

We propose the *Drosophila* InR Val*810*Asp substitution disrupts L2-FnIII linker movement and destabilizes of the protomer’s intracellular position. This event biases the direction of asymmetric transphosphorylation. Thus, *InR*^*+*^*/InR*^*E19*(HR)^ has wildtype traits because while the Val*810*Asp protomer loses its own capacity to be transactivated, it can still transphosphorylate the opposing, normal kinase domain and this is sufficient for full receptor function. *InR*^*E19*(HR)^*/InR*^*E19*(HR)^ is lethal because no dimers are transactivated, although some viability might be restored at lower temperatures (3).

### InR^74(HR)^: kinase domain activation loop

The activation loop (A-loop) of the *Drosophila* insulin/IGF-like receptor is nearly identical to that of human IR and IGF1R (74). This highly conserved loop begins with a DFG motif, followed by five amino acids (MTRDI), a seven amino acid sequence containing three tyrosines that participate in transphosphorylation (Ebina human IR: Tyr*1158*, Try*1162*, Tyr*1163*), and a final 13 residues (75). In unliganded receptors, the A-loop blocks the kinase catalytic cleft and the Mg-ATP binding site of its own protomer. Ligand binding induces A-loop transphosphorylation to alleviate this autoinhibition.

Drosophila *InR*^*74*(HR)^ is an Ile*1543*Phe substitution of the MTRDI motif, plus-one residue from Tyr*1544* (human Tyr*1158*, Ebina). The MTRDI segment becomes structurally organized when the A-loop is transphosphorylated to provide a platform for substrate binding, and this structure could be destabilized by the Ile*1543*Phe substitution (76-78). Alternatively, Ile*1543*Phe may reduce the ability for Tyr*1544* to be phosphorylated and thus diminish the domain’s catalytic activity (75, 79, 80).

Because *InR*^+^/*InR*^*74*(HR)^ generates normal phenotypes, we predict the protomer with the Ile*1543*Phe substitution is still able to transactivate its complementing wildtype protomer. In heteroallelic *InR*^*E19*(HR)^/*InR*^*74*(HR)^, the Val*810*Asp protomer from *InR*^*E19*(HR)^ cannot be activated, yet it transphosphorylates the protomer containing Ile*1543*Phe; the asymmetry of transphosphorylation is biased. Aging is slowed by the hypomorphic function of the sole activated protomer, all with Ile*1543*Phe kinase domains. To test this explanation, we produced a double mutant where Val*810*Asp and Ile*1543*Phe are in same protomer (*InR*^*E*^19, 74^(HR)^). *InR*^*E*^19, 74^(HR)^ is homozygous lethal, but *InR*^+^/*InR*^*E*^19, 74^(HR)^ has normal phenotypes. These substitutions extend longevity only when they *trans*-complement, supporting our model whereby aging is slowed when Ile*1543*Phe resides in the transphosphorylated kinase domain.

### InR^246(HR)^: kinase domain N-terminal lobe

*InR*^*246*(HR)^/*InR*^*E19*(HR)^ delays development and produces small adults that otherwise have normal fecundity and life expectancy (although with a survival crossover). The *InR*^*246*(HR)^ allele is a Val*1384*Met substitution at the invariant ATP binding loop of the N-terminal lobe (IR Val*1037*, Ebina; IGFR Val*1013*, Fig S3) (77). Val*1384*Met will reduce the catalytic rate of the activated protomer (81, 82). The heterozygote *InR*^*+*^/*InR*^*246*(HR)^ is normal because the wildtype protomer kinase is sufficient when activated, and the *InR*^*246*(HR)^/*InR*^*246*(HR)^ is lethal because neither protomer provides enough catalytic activity. *InR*^*E19*(HR)^/*InR*^*246*(HR)^ is viable but small, suggesting either that the protomer from *InR*^*E19*(HR)^ might act as a weak substrate, or the Val*1384*Met protomer produces enough catalytic activity to permit growth although with high age-independent adult mortality.

### InR^211(HR)^: kinase domain C-terminal lobe

The *InR*^*211*(HR)^ allele is normal when heterozygous with wildtype, lethal as a homozygote and extends lifespan when heterozygous with *InR*^*E19*^. Unexpectedly, *InR*^*211*(HR)^ also complements *InR*^*74*(HR)^ to produce small, long-lived adults with low fecundity; their heterodimeric receptors must slow aging by altering the quality of signaling rather by reducing the number of normal receptors.

The Gly*1598*Arg substitution of *InR*^*211*(HR)^ is in the C-terminal lobe of the kinase domain, at a conserved site with human IR (Gly*1212* Ebina) and IGF1R (Gly*1215*) These sites are three conserved residues from the site of the canonical *C. elegans* longevity allele *daf-2*(e1370), Pro*1466*Ser. We can gain insights on this segment from human the fibroblast growth factor receptor FGFR3 where the homolgous C-terminal region stabilizes the asymmetric dimer interface required for transphosphorylation (64). *C. elegans* Pro*1466* corresponds to P*694* of FGFR3, and substitution to serine will disrupt a stabilizing hydrogen bond in this interface. The Drosophila G*1598*R substitution corresponds to G*697* of FGFR3 and this change is likely to disrupt an interface hydrophobic pocket. Based on the FGFR3 structure, we propose that loss of stability at the dimer interface reduces the ability of the protomer with Gly*1598*Arg to act as the enzyme in A-loop transphosphorylation, but maintains its capacity to be transactivated. Like *InR*^*E19*(HR)^, the biased asymmetry of *InR*^*211*(HR)^ may explain why the allele is normal over wildtype while it complements mutations on opposing protomers to extend lifespan.

### InR^353(HR)^: kinase insert domain

The *InR*^*353*(HR)^ allele is exceptional. It is homozygous lethal, yet *InR*^+^/*InR*^*353*(HR)^ extends lifespan with normal growth and elevated fecundity. *InR*^*353*(HR)^ combined with *InR*^*E19*(HR)^ and *InR*^*74*(HR)^ also produces long-lived, small adults with reduced fecundity.

Remarkably, the site of *InR*^*353*(HR)^, Arg*1466*, is described in human disease (42). Drosophila Arg*1466* corresponds to IR Arg*1092* (Ebina). Humans with one Arg*1092*Glu allele have mild insulin resistance, while homozygotes have retarded growth, low viability and severe insulin resistance (83). Chen *et al*. (64) recently described Arg*1092* function in human insulin receptors. Arg*1092* sits in the *α*D helix of the kinase insert domain (KID) (107). When the A-loop of the substrate protomer is transphosphorylated, Arg*1092* on the enzyme-protomer forms a salt bridge with the substrate-protomer to stabilize the dimer in its active state. We therefore expect the disease substitution Arg*1092*Glu disrupts the salt bridge and inhibits substrate protomer kinase activation. However, the Arg*1092*Glu protomer itself can be transphosphorylated and serve as the substrate kinase. We surmise human heterozygotes are effectively haplo-insufficient while homozygotes produce no function IR.

The *Drosophila* kinase insert domain is identical to that of human IR at the first three residues (R_[*1466*/*1092*]_PE), and at the distal residues PPT (76, 84). The *InR353*(HR) allele replaces the conserved arginine at *1466* with cysteine (Arg*1466*Cys). We propose this disrupts the dimer interface salt-bridge and produces biased asymmetry: the Arg*1466*Cys protomer becomes the default substrate kinase of InR heterodimers. Because *InR*^+^/ *InR*^*353*(HR)^ has normal growth and fecundity, we infer the Arg*1466*Cys protomer still retains kinase catalytic activity when transphosphorylated. Why then is *InR*^+^/*InR*^*353*(HR)^ long-lived? Aside from biased asymmetry, we hypothesize Arg*1466*Cys also alters the function of the kinase insert domain so as to favor longevity assurance.

Relative to the kinase insert of human IR, the *Drosophila* KID contains 12 additional amino acids including a potential SH2 motif, Tyr(*1477*)-Leu-Asn. This motif occurs in mammalian IRS-2 where it specifies binding with Grb2 (85) and deletion of IRS-2 extends mouse lifespan (30, 86). *Drosophila* Chico likewise contains SH2 binding motifs for Grb/Drk (Grb/Downstream-of-kinase), and a Y*243*A mutation of this Grb/Drk site extends lifespan (87, 88). *Drosophila* InR itself contains adaptor protein binding sites. The C-terminal tail has SH3-sites that recruit Dock to direct axon guidance, and SH2-sites that recruit Chico to regulate growth (89). At the InR N-terminal juxtamembrane domain, SH2B sites recruit Lnk to modulate interaction with Chico, and mutation of *lnk* extends lifespan (90-93). Based on these facts, we propose the *Drosophila* InR kinase insert domain likewise recruits *Drosophila* Grb/Drk, and Arg*1466*Cys impedes this interaction to induce longevity assurance.

### Caveats

Some of our observations do not fit our interpretations. We predict the *InR*^*E19*(HR)^ protomer can transphosphorylate its opposing protomer but not act as a substrate protomer (biased asymmetry), while the *InR*^*74*(HR)^ protomer can act as enzyme or substrate. Thus, *InR*^*E19*(HR)^/*InR*^*74*(HR)^ is long-lived, but we then expect *InR*^*74*(HR)^/*InR*^*74*(HR)^ to transphosphorylate protomers and produce slow-aging adults. Yet, *InR*^*74*(HR)^/*InR*^*74*(HR)^ is inviable. A second case involves *InR*^*E19*(HR)^/*InR*^*211*(HR)^. We propose both protomers cause biased asymmetry, which implies neither can be transactivated. Yet this genotype is viable and long-lived. Finally, we argue *InR*^*353*(HR)^ can only serve as a substrate protomer with normal kinase activity but with altered KID. This accounts for phenotypes of *InR*^*+*^/*InR*^*353*(HR)^ and *InR*^*E19*(HR)^/*InR*^*353*(HR)^ but does not explain why *InR*^*74*(HR)^/*InR*^*353*(HR)^ has even greater longevity. Overall, a more accurate model requires further biochemical and structural data.

### Synthesis: Drosophila InR modulates aging through distinct structure-defined mechanisms

The phenotypes of *InR*^*+*(HR)^/ *InR*^*353*(HR)^ present a paradox: adults are highly fecundity (and large) and yet long-lived. This positive trait association is seen in other cases involving *C. elegans* and *Drosophila* and challenges life-history theory that expects longevity to trade-off with reproduction (1, 56, 94, 95). Here we propose a mechanistic explanation: InR regulates survival in distinct, independent ways (Fig 7E).

The first involves reproductive costs (69). The alleles *InR*^*E19*(HR)^, *InR*^*74*(HR)^, *InR*^*211*(HR)^, *InR*^*246*(HR)^ slow oogenesis, and previous work shows *Drosophila* longevity is increased when germline stem cell activity is suppressed (96). We propose these *InR* alleles *quantitatively* reduce germline proliferation and thus proportionally relieve survival costs-of-reproduction. This explanation is consistent with how life expectancy is negatively associated with per-ovariole oogenesis when the *InR*^*353*(HR)^ allele is treated as a covariate. How these alleles might modulate oogenesis is unknown but we propose they affect the level of kinase activity in ovarioles and secondarily reduce non-autonomous signals regulating survival-reproduction trade-offs.

The second mechanism confers longevity assurance independent of reproductive costs. We propose the kinase insert domain of *InR*^*353*(HR)^ recruits adaptor proteins that regulate longevity assurance – systems that increase robustness, homeostasis and survival. These effects are additive to effects of reproduction on survival. Thus, while *InR*^*+*(HR)^/*InR*^*353*(HR)^ has *elevated* per-ovariole production and may suffer associated survival costs, the genotype gains ∼13 days of reproduction-independent longevity assurance to yield a net increase in lifespan. *InR*^*E19*(HR)^/*InR*^*353*(HR)^ and *InR*^*211*(HR)^/ *InR*^*353*(HR)^ gain the same longevity assurance but now added to benefits from reduced survival costs-of-reproduction.

This hypothesis resonates with data on *Drosophila chico*. Like *InR*^*+*(HR)^/ *InR*^*353*(HR)^, *Drosophila chico*^*wildtype*^/*chico*^1(null)^ adults are large, fecund and long-lived (97-100). Chico contains phosphotyrosine binding sites for Grb2/Drk and p60 of the PI3K complex (87). Mutation to block Grb/Drk (Y*243*) represses Ras-Erk signaling and extends fly lifespan without reducing growth or fertility (87, 88). This pathway mediates the ETS transcription factor Anterior-open (Aop). In a similar way, the *C. elegans daf-2*(*sa223*) allele is proposed to alter Ras signaling and is long-lived (55, 56).

We hypothesize the proposed SH2 motif Tyr(*1477*)-Leu-Asn of *Drosophila* KID facilitates binding of Grb2/Drk. We reason this SH2 binding event is repressed by Arg*1466*Cys of *InR*^*353*(HR)^. Reduced Grb2/Drk recruitment and activation slows aging by specific mechanisms of longevity assurance transduced by Ras, as does the Chico *Y243* substitution. At the same time the *InR*^*353*(HR)^ protomer still activates PI3K-Akt-Foxo signaling to maintain growth and fecundity. Other *InR* mutant alleles, we propose, inhibit PI3K signaling; their substitutions reduce growth and fecundity, which extends lifespan by lessening survival costs-of-reproduction.

We suggest the longevity benefits of Ras-Erk-Aop and PI3K-Akt-Foxo regulate different cellular and physiological systems. The proposed KID-Grb2/Drk network controls longevity through mechanisms that do not involve costs-of-reproduction. Our data support emerging ideas (45, 89, 101, 102) w here protein tyrosine receptors segregate phenotypes through the action of specific sites within particular structural domains.

## METHODS

### Quantitative complementation test

#### EMS mutant InR alleles

Lines with EMS-induced point mutations at *InR* generated on a ‘*rutipa*’ genetic background, balanced by *TM3,Sb* (provided in 1999 by M. Frasch, The Mount Sinai School of Medicine, ‘Frasch series’) (58, 59). We tested the ability of 17 mutant lines from this collection to produce viable adults when complemented to the EMS-induced mutant allele *InR*^E19^ provided by J. Jack (University of Connecticut School of Medicine, in 1999) (64). We previously reported (3) that the *InR*^E19^ allele recombined to a third chromosome marked with *ri red e*^+^ extends life span of males and females when complemented to the P-element allele *InR*^P5545^ (59). Nine alleles of the Frasch series produced viable adults when complemented to *InR*^E19^ *ri red e*^+^ (*InR*^74^, *InR*^76^, *InR*^211^, *InR*^246^, *InR*^262^, *InR*^327^, *InR*^351^, *InR*^353^, *InR*^373^). Each of these alleles and the *InR*^E19^ allele were crossed six generations into the inbred *Samarkand I-236* (*Sam*) background using the stock *Sam 1; Sam 2; TM3*/*TM6B* (provided in 1999 by T. Mackay, North Carolina State University).

#### Wildtype InR^NC^ alleles

Third chromosome extraction lines were derived from wildtype females caught at the Raleigh’s Farmer’s Market in 1999 (provided by T. Mackay, North Carolina State University) (103). Gravid females were crossed to males of *Sam 1; Sam 2; TM3*/*TM6B* and backcrossed to extract third chromosomes balanced by *TM3* in the Samarkand background. Eighteen lines from this collection formed a sample of wildtype *InR* alleles (the *InR* ^NC^ series); in addition, *InR*^NC442^ was used in each block of demographic analysis to provide a common strain to normalize survival data across test periods.

#### Genotypes for Quantitative Complementation Analysis

We made reciprocal crosses between every *InR*^EMS^/*TM3* and *InR*^NC^/TM3 with *InR*E19/TM3 to produce F_1_ genotypes. Offspring of reciprocal crosses were pooled for phenotype analyses. In complementation analysis, phenotypes of every *InR*^EMS^/*InR*^E19^ and *InR*^NC^/*InR*^E19^ genotype were quantified and compared relative to a contemporaneous cohort of its corresponding *InR*^EMS^/*TM3, InR*^*+*^ *Sb* or *InR*^NC^/*TM3, InR*^*+*^ *Sb*.

##### Demographic quantitative complementation

Life tables were generated for cohorts of each F_1_ genotype (methods detailed in Supplemental Methods). Cohorts for each genotype were initiated with three replicate demography cages of ∼150 newly eclosed adults at ∼1:1 sex ratio. In total we assessed the lifespan of 16,057 males and 17,223 females. Each two days food vials were changed and dead flies were removed and counted. Life tables were constructed by the extinct cohort method with data from replicate cages within a cohort combined for survival analysis.

Life table data were collected across five test blocks (blocks *j* = 1 to 5). To unify data across blocks, in every block we included the wild-derived line *InR*^NC442^. Estimated survival parameters for each genotype within each block were normalized relative to *InR*^NC442^ of the corresponding block. We generated one lifetable for each *InR*^EMS^ mutant and *InR*^NC^ allele, expect for *InR*^211^, *InR*^74^ and *InR*^353^, which were studied twice.

To conduct demographic Quantitative Complementation Testing, within each block *j*, we used proportional hazard analysis (104) executed in JMP (SAS Institute, Cary, NC) to quantify the proportional mortality difference (hazard coefficient *β*_*ij*_) for each *InR* allele *i* (EMS or NC) when complemented to *InR*^E19^ relative to when complemented to TM3, *InR*^*+*^ *Sb*. In every block we estimated *β*_*NC*442,*j*_ and calculated the *adjusted mortality coefficient* for each *InR* allele as 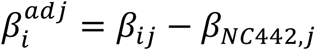. Negative values of 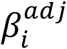 indicate mortality rate is reduced in *InR*^*i*^/*InR*^E19^ relative to *InR*^*i*^/*TM3, InR*^*+*^ *Sb*; values near zero indicate there is no effect on mortality and values greater than zero indicate there is increased mortality; 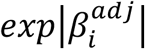 estimates the fold change in mortality risk (hazard ratio) induced by each *InR* allele complemented to *InR*^E19^ relative to when it is complemented to *TM3, InR*^*+*^ *Sb*. Life expectancy was also standardized among blocks: for each block *j* we calculated the difference in life expectancy between *InR*^NC442^ /*InR*^E19^ and *InR*^NC442^ / *TM3, InR*^*+*^ *Sb* denoted 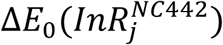. The adjusted difference in life expectancy for each *InR* allele *i* is 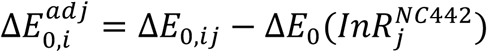.

#### Body size quantitative complementation

Head capsule width was measured from 20 females from each *InR* EMS-mutant and NC-wildtype allele when complemented to *InR*^E19^ and to *TM3, InR*^*+*^ *Sb*. Adults were reared simultaneously on standard diet from vials seeded with 100 eggs, and collected from across three replicate vials. The effect of allele *InR*^*i*^ on body size, *σ*_*i*_, is genotypic mean of *InR*^*i*^ /*TM3, InR^+^ Sb* minus genotypic mean of *InR*^*i*^ /*InR*^E19^.

#### Drosophila insulin receptor alignments with human IR and IGF1R

The annotation of *Drosophila* InR and human IR and IGF1R varies among sources (Fig 3, S1). The human insulin receptor (IR) is referenced as the mature long isoform of Ebina (105), and the Human Genome Variation Society (HGVS, http://hgvs.or/rec.html) that includes an additional 27 a.a. localization sequence (Fig 3, Fig S1). The human IGF1R sequence is numbered as NCBI Reference Sequence: NP_000866.1. *Drosophila InR* numbering begins at the first translation initiation site in *w*Dah wildtype (GenBank accession MT_563159), +45 amino acids relative to the TIS reported by Fernandez (59) (NCBI Reference Sequence: NM_079712.6), which also differs by an insertion/deletion in the L1 ectodomain (*w*Dah lacks H*144*) and the substitution H*146*Q. This polymorphism segregates along latitudinal clines of *Drosophila* natural populations (52). See Supplemental Materials (Fig S3, S4) for confirmed alignments of the homologous recombination allele series.

#### Sequence of EMS InR alleles

The EMS-generated *InR*E19 allele was sequenced using (rare) homozygous F_1_ adults generated from an *InR*^E19^/TM3 stock (64). This allele contained several polymorphisms compared to *InR* of *w*Dah, including the unpublished substitution GTC/GAC, Val*810*Asp (R. Kohanski, personal communication, Feb. 2004). The identity of the *InR*^246^ and *InR*^74^ substitutions were provided by R. Kohanski (unpublished, personal communication, Feb. 2004). The substitutions for EMS alleles *InR*^353^ and *InR*^211^ were reported by Brogiolo (42). In the current work, each of these described substitutions were used to design site directed mutagenesis with homologous recombination to produce coisogenic, single amino acid substitution alleles.

#### InR homologous recombination allelic series

Details of fly stocks and culture, constructs, primers, cloning and work flow for homologous recombination are in Supplemental Materials. In brief, targeting arms for ends-out homologous recombination of *InR* (cloned from *w*Dah) (106) were cloned into the pW25.2 vector (Drosophila Genomics Resource Center, Bloomington, IN) and injected into *w*^*1118*^ embryos by Genetic Services, Inc. (Sudbury, MA). Targeting arms contained single nucleotide substitutions to specify the desired amino acid replacements (Fig 3), while the native *w*Dah sequence provided both arms to produce *InR*^+(HR)^. Flies with the *white*^*+*^ eye color marker mapped to the first or second chromosomes were used for homologous recombination. These lines were crossed to flies expressing flipase and the restriction enzyme *I-Sce1* to execute excision and linearization of the transgenic construct. Replicate, independent (accession) candidate insertion lines were selected based on *white*^*+*^ mapped to chromosome 3. Gene replacement and mutant validation were confirmed by sequencing. Strains with gene replacement strains were crossed to flies expressing *cre* recombinase to remove the *white*^*+*^ marker. The *white*^*+*^ marker was retained to produce *InR*^null(HR)^.

#### Phenotypes of InR homologous recombination allelic series

##### Viability, development time, adult size

Reciprocal, diallel crosses were made among all *InR*^(HR)^ alleles. In replicate vials with yeast-supplemented standard media, four virgin females and males were mated over 24h and removed. Stage of lethality (larvae, prepupae or pupae) was recorded to determine viability. Eclosing adults were recorded every 24 hours. Size (thorax length) was measured from 20 F_1_ females of each viable cross.

##### Lifetable demography

Newly eclosed male and females were collected, transferred to bottles with fresh medium and kept at 25°C for two days (details in Supplemental Materials). Mated females were transferred to demography cages maintained at 25°C at an initial density of 125 females per cage. Three to five replicate cages were initiated for each genotype cohort. Food was changed every two days, at which time dead flies were removed and counted. Life tables were constructed by the extinct cohort method with data from replicate cages within a cohort combined for survival analysis. In total we assessed the lifespan of 17,184 HR females.

##### Fecundity and ovariole number

Fecundity was measured twice daily from eclosion through 15 days old. For each genotype, 15 newly eclosed females were placed with 15 *w*Dah males into three replicate demography cages, modified for egg collection (method in Supplemental Methods). Egg plates were supplemented with yeast paste and changed twice a day at which time eggs were counted. Dead females were removed and counted daily. Daily fecundity (mean eggs per day) is total daily eggs across replicate cages/number of alive females each day. Average fecundity is the daily fecundity averaged across all days. Ovary number was measured from 10-day old females of each genotype. Ovaries were dissected in PBS and fixed with 4% paraformaldehyde. The number of ovarioles per ovary was recorded as the sum over both ovaries from 15 females of each genotype.

## Author contributions

R.Y., M.P. and N.C-J conducted the experiments. R.Y., M.P. and M.T. designed the experiments. M.T. and R.Y. analyzed the data and wrote the paper.

## Acknowledgements

For patient mentoring and discussion on insulin receptor structure and function that made this work possible, MT thanks P. de Meyts, R. Kohanski, B. Garofalo, B. Forbes, Y. Suh, X. Bai, M. Mohammadi, S. Takahashi, and W. Peti. M. Frasch generously shared his archival collection of *InR* EMS mutants. R. Kohanski shared unpublished data in the sites of substitutions in the Frasch alleles. W. Peti provided the structural diagram of substitutions mapped to the kinase domain. Thanks for supporting work in the Tatar lab is due to Aleksandra Norton, Miyo Malouf, Richard Martinez, Dante Ordantella, Colette Toal, Roy Hsu. Funding was provided by NIH R01AG16632 and R37AG024360.

