## Supplemental Methods and Data for "Mapping *Drosophila* insulin receptor structure to the regulation of aging through analysis of amino acid substitutions"

Recombination. Methods Mol Bio 718: 41-73). *InR* genomic DNA from the Dahomey wildtype (*w*Dah) was extracted, PCR amplified using InR Arm primers (**Methods Table A**) and subcloned into the pTopo vector to generate constructs pTopo-Arm1 (3' end of gene) and pTopo-Arm2 (5'end of the gene). The $w^{+}$ marker was targeted into the 9$^{th}$ intron of InR, dividing this sequence into two segments: Arm1 with 2,686 bp and Arm2 with 3,614 bp. To subclone both arms into the pW25.2 transformation vector, Arm1 was flanked by restriction sites *AscI* and *BsIWI* and Arm2 by *Not*I and *Acc65I*. Since *w*Dah contains an internal site of *Acc65I* in Arm2, this site was abolished by site directed mutagenesis without altering its coded amino acid (from GGTACC to GCTACC, at position 1194 of Arm2). All subsequent cloning was made using Arm2 with the abolished *Acc65I* site. Individual clones containing pTopo-Arm1 and pTopo-Arm2 were completely sequenced using primers listed in Table B. Complete sequence of *InR* $w^{Dah}$ Arm1 and Arm2 in **Methods Fig A, B**.

Placement of homologous recombination targeting arms on InR.

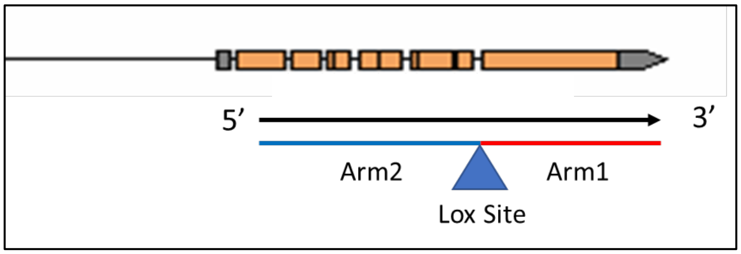

**InR single nucleotide mutations for homologous recombination**

The construct pTopo-Arm2 was used as a template for the site-direct mutagenesis to introduce the InR$^{e19}$ mutation, a modification from T-A at position 1890 of Arm2. The construct pTopo-Arm1 was used as a template for the site-direct mutagenesis to introduce the following independent mutations: InR$^{246}$, InR$^{353}$, InR$^{74}$ and InR$^{211}$. InR$^{246}$ has a G-A modification at position 291, InR$^{353}$ has a C-T modification at position 537, InR$^{74}$ has a A-T modification at position 768 and InR$^{211}$ has a G-A modification at position 933 of Arm1. After mutagenesis, constructs were fully sequenced to confirm the single nucleotide mutation. Location of substituted nucleotides are underlined in **Methods Figures A and B**.

**Arm 2 = 3614 bp**

| Mutation | Position in Arm 2 | Nucleotide change | Aminoacid change |
|---|---|---|---|
| InR[e19] | 1890 | GTC/GAC | Val-Asp |

**Arm 1 = 2686 bp**

| Mutation | Position in Arm1 | Nucleotide change | Aminoacid change |
|---|---|---|---|
| InR[246] | 291 | GTG/ATG | Val/Met |
| InR[353] | 537 | CGT/TGT | Arg/Cys |
| InR[74] | 768 | ATC/TTC | Ile/Phe |
| InR[211] | 933 | GGA/AGA | Gly/Arg |

**Cloning into transformation vector**

The wild type and the mutated InR were sequentially cloned into transformation vector pW25.2. Arm1 wild type or mutated was cloned into the restriction sites AscI and BsiWI. Arm2 wild type of mutated was inserted into restriction sites Acc65I and NotI. Seven different constructs were created, allowing for wild type, single mutants for each one of the five single nucleotide substitutions, as well as one double mutant, InR$^{e19\text{-}74}$. Constructs were then transformed into Drosophila $w^{1118}$ embryos by Genetic Services, Inc. (Sudbury, MA).

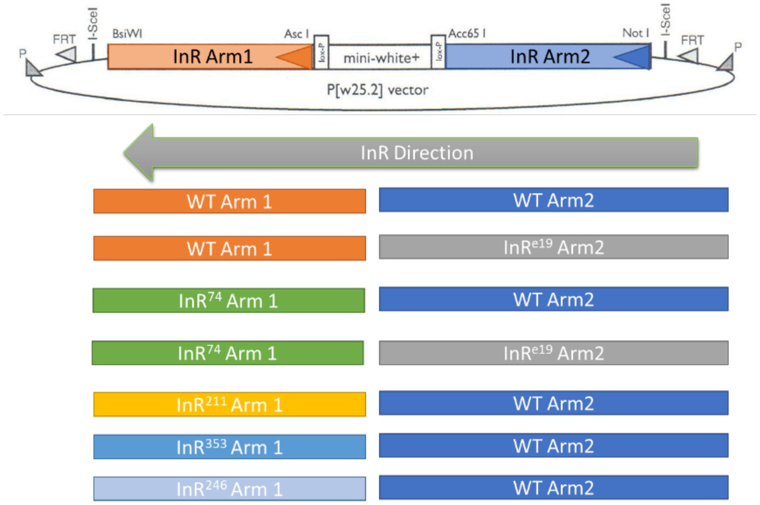

Flies caring the *white*$^+$ eye color marker mapped to the first or second chromosome were used to target homologous recombination by crossing with a line expressing flipase and the restriction enzyme I-Sce1 for excision and linearization of the transgenic construct. Potential lines were selected for *white*$^+$ insertions on chromosome 3, and confirmed by PCR amplification and sequencing. Accession with successful gene replacement were then crossed to flies expressing *cre* recombinase for removal of the *white*$^+$ eye color marker.

**Confirmation of mutations**

Final lines were sequenced to confirm fidelity and homolgous recombination of the replacement allele. using primers internal of homologous recombination construct and primers upstream of Arm2 and downstream of Arm1 into the genomic region. PCR fragments were sequenced to completion (primers in **Methods Table B**). The wild type strain *w*Dah was also sequence to completion, and the following primers were used for its amplification (Arm1: P-InR63 and P-InR60; Arm2: P-InR56 and InR61).

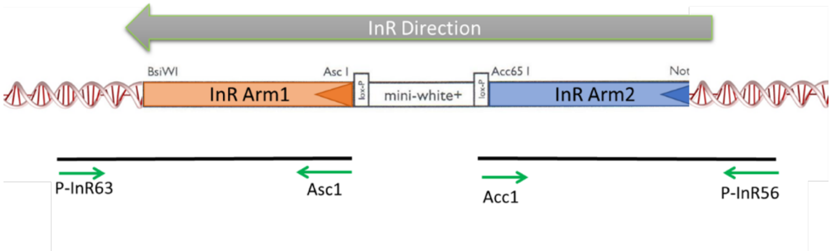

### **Methods Fig A:** w$^{Dah}$ Arm1 sequence cloned into pTOPO

#### w$^{Dah}$ Arm1 sequence cloned in pTOPO

GAATGATCATTTTTAAATGAATTTTTTTTTTTCATTAAATTCAACAGCCTCCGCCGAGCTATGCTAAGGTCTTTTTCTGGCTACTGGGAATCGGCCTAGC < 100

GTTCCTGATCGTTTCCCTGTTCGGCTATGTCTGTTACCTGCACAAGAGGAAGGTTCCCTCTAATGACCTCCATATGAACACAGAGGTGAATCCGTTCTAT < 200
A
GCGAGCATGCAATACATCCCAGACGATTGGGAGGTGCTGCGAGAGAACATCATTCAGTTGGCTCCACTAGGCCAGGGATCCTTTGGCATG**G**TGTATGAGG < 300

GTATCCTGAAGTCCTTTCCACCCAATGGCGTGGATCGCGAGTGTGCCATTAAGACTGTCAACGAAAATGCTACGGATCGCGAGCGAACCAATTTCCTGAG < 400

CGAGGCGAGCGTCATGAAGGAGTTCGATACGTATCATGTCGTAAGATTGCTCGGTGTTTGTTCCAGGGGTCAGCCGGCTCTGGTGGTCATGGAGCTAATG < 500

AAGAAGGGTGATCTTAAGTCCTATTTGCGTGCCCAT**C**GTCCCGAGGAGCGGGATGAGGCCATGATGACGTATCTTAATCGCATCGGAGTGACTGGTAATG < 600

TGCAGCCTCCTACTTATGGAAGAATCTACCAGATGGCCATTGAGATTGCGGATGGCATGGCATATTTGGCCGCCAAGAAGTTCGTCCATCGTGATCTTGC < 700

AGCTCGAAATTGCATGGTTGCTGATGATTTGACGGTGAAAATTGGTGACTTTGGAATGACCCGTGAC**A**TCTATGAGACGGATTACTATCGGAAGGGCACT < 800

AAAGGGCTGCTGCCAGTTCGCTGGATGCCACCGGAGAGCTTGCGAGATGGTGTCTACTCTAGTGCCAGTGATGTATTCAGCTTTGGAGTGGTTCTCTGGG < 900

AAATGGCCACCTTAGCGGCTCAGCCATACCAG**G**GACTTTCCAACGAGCAAGTCCTGCGTTACGTTATCGATGGCGGTGTTATGGAGAGGCCGGAAAATTG < 1000

TCCTGATTTTCTGCATAAACTAATGCAAAGGTGCTGGCATCATAGGTCTTCGGCGAGACCCAGTTTTCTGGATATCATTGCGTATCTCGAACCACAATGC < 1100

CCCAATTCACAATTTAAGGAAGTATCCTTCTATCACTCAGAGGCAGGTCTGCAGCATCGGGAAAAGGAGCGCAAGGAACGCAATCAGCTAGATGCATTCG < 1200

CGGCAGTCCCCTTGGATCAAGATCTGCAGGATCGGGAACAGCAGGAGGATGCTACCACACCTTTACGAATGGGCGATTATCAGCAGAACTCCTCGTTGGA < 1300

TCAACCGCCCGAAAGCCCCATCGCCATGGTTGATGATCAGGGTTCTCACTTGCCATTTAGCCTGCCATCCGGATTCATTGCGAGCAGTACTCCTGATGGG < 1400

CAGACTGTAATGGCTACTGCTTTCCAGAATATTCCAGCAGCGCAAGGTGATATTTCGGCGACTTACGTTGTGCCTGATGCAGACGCTTTGGACGGCGACA < 1500

GGGGATATGAGATCTACGATCCCAGTCCGAAATGTGCAGAGCTGCCGACGAGCAGAAGTGGCAGTACTGGCGGTGGAAAACTCAGCGGAGAACAACATTT < 1600

GCTGCCAAGAAAGGGTCGCCAGCCTACCATCATGAGCAGCTCGATGCCAGATGATGTCATCGGTGGGTCCTCACTGCAACCCTCGACTGCGTCAGCAGCC < 1700

AGTTCTAATGCCAGTTCGCACACAGGACGCCCAAGTCTGAAGAAAACAGTGGCGGATTCGGTTCGCAATAAGGCAAACTTCATTAATCGCCACCTATTTA < 1800

ACCACAAGCGAACGGGCAGCAATGCCAGCCACAAGAGCAATGCCTCCAATGCTCCGAGTACCAGCAGTAACACCAACTTGACAAGTCACCCAGTGGCTAT < 1900

GGGCAATCTTGGAACTATCGAGAGTGGTGGCAGTGGTTCGGCTGGTAGTTATACTGGAACACCCCGCTTCTATACTCCATCAGCGACGCCTGGAGGAGGC < 2000

AGCGGTATGGCCATTAGCGACAATCCTAACTACAGACTACTAGACGAGTCAATAGCCAGCGAACAGGCCACCATCCTAACGACTAGCAGCCCCAATCCCA < 2100

ACTACGAGATGATGCATCCACCAACCAGTCTGGTCAGTACCAATCCGAACTATATGCCCATGAATGAGACTCCAGTGCAGATGGCGGGTGTGACCATTAG < 2200

CCATAATCCGAATTACCAGCCCATGCAGGCGCCGTTGAATGCACGCCAAAGCCAAAGTAGCTCCGACGAGGACAACGAGCAGGAGGAGGACGATGAGGAT < 2300

GAGGACGACGACGTGGACGATGAGCATGTGGAGCACATCAAGATGGAGCGCATGCCATTGAGTCGGCCCAGGCAAAGAGCGTTGCCCAGCAAGACGCAGC < 2400

CGCCTCGCAGTCGCAGCGTCAGCCAAACGAGAAAATCTCCTACGAATCCCAACTCCGGAATCGGAGCGACAGGAGCCGGAAACCGATCCAACTTGCTTAA < 2500

AGAGAACTGGCTGCGACCGGCGAGTACGCCAAGGCCTCCACCACCCAATGGATTCATCGGAAGGGAGGCGTAATCGTTACGAACTGTAGTTCTGTAGAAA < 2600

AACATGTAAATAGATGAGGAGAAGCAGCATGGATATAGATAGAGTAGCAAGTCTTTGCCGCAAGCCAAGCTTCTTGGAATTTGACT < 2686

**Methods Fig B:** w$^{Dah}$ Arm2 sequence cloned into pTOPO. The Acc65I restriction site in region was abolished by site-specific mutagenesis, at position 1194 (marked in red).

### w$^{Dah}$ Arm2 sequence cloned in pTOPO

ATATAGCCGATGGACTGGATAAAACAGCGCTTTCGGTGTCGGGGACGCAATCGCGATGGACAAGGAGCGAATCAAACCCAACAATGCGACTGTCACAAAA < 100

TGTAAAACgtatgtattacctatgatctaccttgcaagttaaaggaacattcccatttgcacgagactgacagatactaatagcttttccattaacacct < 200

tatagCTTGCAAATCCATGGACATCAGGAACATGGTGTCGCACTTCAATCAGCTGGAGAACTGCACGGTCATCGAGGGCTTCCTGCTGATCGATTTGATA < 300

AACGACGCCAGCCCTCTGAACAGAAGCTTTCCAAAACTGACCGAGGTCACAGATTATATCATAATCTACCGTGTGACTGGATTGCACTCGCTGTCAAAGA < 400

TCTTTCCCAATCTGAGCGTCATTAGGGGAAACAAGCTGTTCGACGGATATGCCTTGGTCGTCTACTCGAATTTCGACCTCATGGATTTGGGACTTCACAA < 500

GCTACGATCCATAACCAGAGGCGGTGTGCGGATTGAGAAGAATCATAAGCTGTGCTATGATAGGACCATCGATTGGCTGGAAATTCTGGCGGAAAACGAA < 600

ACCCAACTGGTGGTGCTGACAGAGAACGGCAAGGAGAAGGAGTGCAGGCTTTCCAAGTGCCCGGGGGAGATCAGAATTGAGGAGGGGCACGATACCACGG < 700

CTATTGAGGGAGAGCTTAATGCCAGTTGTCAGCTGCACAATAATAGGCGCCTGTGCTGGAACAGCAAACTCTGCCAGACGAgtgagttggccggtgtaaa < 800

gttataccgttttcttattaacatttttcgcgtttttttcttctccagAATGCCCTGAAAAGTGCAGAAATAACTGCATCGATGAGCACACCTGCTGCAG < 900

CCAGGATTGTTTGGGTGGATGCGTGATCGATAAGAATGGGAATGAGAGCTGCATCTCCTGTCGAAATGTGTCTTTCAACAACATCTGTATGGACTCCTGT < 1000

CCGAAAGGCTATTATCAGgtaactaatgattcacttttagtatacaacaagtagctaactttcaaactcttttgcagTTCGACAGCCGCTGCGTAACGGC < 1100

GAACGAGTGCATCACACTGACAAAGTTTGAAACGAACAGTGTGTATTCCGGTATTCCATACAACGGACAATGTATCACCCACTGTCCAACGGG**C**TACCAG < 1200

AAGTCAGAGAACAAGCGCATGTGCGAACCTTGTCCGGGCGGCAAGTGTGACAAGGAGTGCTCCTCCGGTCTTATCGACAGTTTGGAGCGTGCTCGGGAGT < 1300

TCCACGGCTGCACCATTATAACCGGAACCGAGCCCCTTACCATCAGCATTAAACGTGAAAGCGGCGgtaagtgtttttttgctgctctaaattaaagtata < 1400

atctctatattaaaacttcttgtttcttcagCTCACGTCATGGATGAATTAAAATATGGCCTGGCTGCCGTCCATAAAATTCAGTCGTCCCTAATGGTTC < 1500

ATTTGACCTACGGATTGAAGTCCTTGAAATTCTTTCAATCCCTAACTGAAATTAGCGGCGATCCGCCGATGGACGCGGATAAATATGCTTTGTATGTGCT < 1600

TGATAATCGCGATCTAGATGAGCTCTGGGGACCCAACCAAACGGTGTTCATTAGGAAGGGCGGCGTCTTCTTTCATTTCAACCCAAAACTATGTGTGTCC < 1700

ACCATTAACCAGTTGCTGCCCATGCTGGCCTCCAAGCCAAAGTTTTTTGAAAAGTCAGATGTGGGCGCAGACTCGAATGGAAACCGCGGATCATGTAAGT < 1800

AATCTTTAACGAAGATTTTAGTAAAATTTTAAACTTTGTCAGTACTAACCTAAATGAAACATTTCCAGGTGGAACAGCCGTTCTCAATG**T**CACATTACAA < 1900

TCAGTGGGAGCAAACTCCGCTATGCTGAACGTCACGACAAAAGTTGAAATAGGAGAGCCCCAAAAGCCGAGCAATGCTACAATTGTTTTTAAGGATCCGC < 2000

GCGCCTTCATCGGTTTCGTGTTTTATCATATGATCGATCCGTACGGGAACTCAACTAAAAGCAGTGACGATCCATGCGATGATCGCTGGAAGGTTAGCTC < 2100

TCCGGAAAAGAGCGGGGTCATGGTATTAAGCAATTTGATTCCGTACACTAACTACTCCTACTACGTTCGGACCATGGCTATATCCTCGGAATTGACAAAC < 2200

GCGGAGAGCGACGTGAAGAACTTTAGGACGAATCCCGGACGACCGTCAAAGGTTACGGAGGTGGTAGCAACCGCCATTTCAGATTCGAAAATTGTGAGTA < 2300

TGAAATTGTGCAATAGATTAGTTTAGGATTATAAATGATTACTAAATGTCAAGTAATTGTTCACTTTGACTGATCAACATTTATTGCCTTTTCAGAACGT < 2400

AACATGGAGCTACCTAGATAAGCCTTATGGCGTGCTAACGCGCTATTTTATAAAAGCCAAACTTATAAATCGGCCTACTCGAAACAATAACCGGGATTAC < 2500

TGTACTGAACGTAAGCAACAACATAGCCAAATTGTATCCATAATTAATTAATTTAAATATTTCATCTGTGTAGCTCTCGTCAAGGCCATGGAAAATGACC < 2600

TGCCAGCCACAACGCCTACCAAGAAAATATCAGATCCTTTAGCAGGCGACTGTAAGTGCGTGGAGGGTTCGAAGAAGACTAGCAGTCAGGAATACGATGA < 2700

TCGTAAAGTTCAAGCGGGCATGGAGTTTGAGAACGCGTTGCAAAACTTTATATTTGTTCCAAACATTCGGAAAAGCAAGAATGGATCGTCTGACAAATCA < 2800

GACGGAGCGGAAGGTGCAGCTCTCGATTCTAATGCTATTCCAAATGGAGGAGCTACTAACCCTTCACGTAGAAGGAGAGACGTTGCGCTCGAGCCAGAGC < 2900

TCGACGATGTAGAGGGCAGTGTACTTCTACGCCATGTGCGCTCCATCACAGACGATACCGATGCATTTTTCGAAAAGGACGACGAAAATACCTATAAAGA < 3000

CGAAGAAGACTTGTCCTCCAACAAACAATTCTATGAGGTGTTTGCCAAGGAATTGCCACCAAATCAAACACATTTTGTCTTTGAAAAACTGCGCCACTTC < 3100

ACCCGCTACGCTATCTTCGTGGTAGCCTGTAGAGAAGAAATCCCCAGCGAAAAATTAAGGGACACCAGTTTTAAGAAGTCGCTCTGCAGCGATTATGACA < 3200

CCGTTTTCCAAACTACAAAGAGAAAGAGTAGGTGGACTTGAGAGCGTGTTTACTTTCATTACTAACGATTTGGTTTTACTTATAGAATTTGCCGACATAG < 3300

TCATGGACCTAAAAGTAGATTTAGAACACGCCAACAACACCGAGTCCCCAGTACGGGTTCGCTGGACGCCACCAGTAGATCCCAACGGAGAAATTGTCAC < 3400

CTATGAAGTGGCCTACAAGTTGCAAAAACCCGATCAAGTGGAAGAAAAGAAGTGCATTCCGGCTGCTGACTTCAACCAGACTGCCGGTTATTTAATAAAG < 3500

CTCAACGAGGGCCTTTACAGCTTCAGGGTGCGAGCCAATTCAATAGCGGGATACGGCGATTTCACGGAAGTCGAACATATAAAAGTTGAGGTAGGTTGAA < 3600

TAATTTTTATCTGC < 3614

**Methods Table A:** Primers for cloning InR Arm1 and Arm2, and for site-direct mutagenesis. Sequences 5'-3'.

| **InR primers – used for cloning and site-direct mutagenesis** | | |
|---|---|---|
| **Primer Name** | **Primer Sequence** | **Use of primer** |
| P- InR 13 | TAATTAATTGGCGCGCCGAATGATCATTTTTAAATGAATTTTTTTTTTTCATTAAATTCA | 5' HR Region for InR Arm1; forward |
| P- InR 14 | ATATCGTACGAGTCAAATTCCAAGAAGCTTGGCTTGCGGCAAAG | 3' HR Region for InR Arm1; reverse |
| P- InR 15 | TAATTAATTGCGGCCGCATATAGCCGATGGACTGGATAAAACAGCGCTTTCGGTG | 5' HR Region for InR Arm2; forward |
| P- InR 16 | ATATGGTACCGCAGATAAAAATTATTCAACCTACCTCAACTTTTATATGTTCGACTTCCG | 3' HR Region for InR Arm2; reverse |
| P- InR 29 | CCACTGTCCAACGGG**c**TACCAGAAGTCAGAG | Primer for Acc65I modification in Arm2; forward |
| P- InR 30 | CTCTGACTTCTGGTA**g**CCCGTTGGACAGTGG | Primer for Acc65I modification in Arm2; reverse |
| P- InR 39 | GGAACAGCCGTTCTCAATG**a**CACATTACAATCAGTGGGA | InR$^{e19}$ mutagenesis Arm2, forward |
| P- InR 40 | TCCCACTGATTGTAATGTG**t**CATTGAGAACGGCTGTTCC | InR$^{e19}$ mutagenesis Arm2, reverse |
| P- InR 41 | ACTTTGGAATGACCCGTGAC**t**TCTATGAGACGGATTACTAT | InR$^{74}$ mutagenesis Arm1, forward |
| P- InR 42 | ATAGTAATCCGTCTCATAGA**a**GTCACGGGTCATTCCAAAGT | InR$^{74}$ mutagenesis Arm1, reverse |
| P- InR 43 | GGCTCAGCCATACCAG**a**GACTTTCCAACGAGCA | InR$^{211}$ mutagenesis Arm1, forward |
| P- InR 44 | TGCTCGTTGGAAAGTC**t**CTGGTATGGCTGAGCC | InR$^{211}$ mutagenesis Arm1, reverse |
| P- InR 45 | TATTTGCGTGCCCAT**t**GTCCCGAGGAGCGGG | InR$^{353}$ mutagenesis Arm1, forward |
| P- InR 46 | CCCGCTCCTCGGGAC**a**ATGGGCACGCAAATA | InR$^{353}$ mutagenesis Arm1, reverse |
| P- InR 51 | GCCAGGGATCCTTTGGCATG**a**TGTATGAGGGTATCCTGAAG | InR$^{246}$ mutagenesis Arm1, forward |
| P- InR 52 | CTTCAGGATACCCTCATACA**t**CATGCCAAAGGATCCCTGGC | InR$^{246}$ mutagenesis Arm1, reverse |
| P- InR 56 | CGGCTGCAACAGAGTGTGAGAGCGGGACGA | Arm2 HR confirmation, forward |
| P- InR 60 | TCGGCCTACTCGAAACAATAACCGGGATTACTGTACTGAACgtaagc | w$^{Dah}$ Arm1, forward |
| P- InR 61 | CTTTTCCCGATGCTGCAGACCTGCCTCTGAGTGATAGAAGGATACTTCCTTAAATTGTGA | w$^{Dah}$ Arm2, Reverse |
| P- InR 63 | CACTATCGTTTTCTGCTCTCGTTTTGTATTGCGTTAGAGGCTCTAGTGCATTA | Arm1 HR confirmation, reverse |
| P- AscI | GTATGCTATACGAAGTTATCTAGACTAGTCTAGGGCG | Arm1 HR confirmation, forward |
| P- AccI | CATTATACGAAGTTATCTAGACTAGTCTAGGGTAC | Arm2 HR confirmation, reverse |
| P- SP6 | GATTTAGGTGACACTATAG | pTopo sequencing |
| P- T7 | TAATACGACTCACTATAGGG | pTopo sequencing |

**Methods Table B:** Primers for sequencing Arm1 and Arm2. All primers were used for sequencing confirmation of homologous recombination events. Primers used to sequence DNA cloned in pTopo are highlighted in the right column.

| **InR primers – used for sequencing** | | | |
|---|---|---|---|
| **Primer Name** | **Primer Sequence** | **Use of primer** | **pTopo** |
| P- InR 02 | GCACAATTACTCTTACTCGCCTGG | Arm2; forward | |
| P- InR 06 | TCAACGAGGGCCTTTACAGCTTCA | Arm2; forward | |
| P- InR 08 | GTTACAATGGATTGCTAGCCAGTC | Arm1; reverse | |
| P- InR 17 | CACAAGCTACGATCCATAACCAGAGG | Arm2; forward | √ |
| P- InR 18 | TGTATGGACTCCTGTCCGAAAGGCTA | Arm2; forward | √ |
| P- InR 19 | CAGTCGTCCCTAATGGTTCATTTGAC | Arm2; forward | √ |
| P- InR 20 | ATGATCGATCCGTACGGGAACTCAAC | Arm2; forward | √ |
| P- InR 21 | CCGGGATTACTGTACTGAACGTAAGC | Arm2; forward | √ |
| P- InR 22 | GACGAAGAAGACTTGTCCTCCAAC | Arm2; forward | √ |
| P- InR 23 | TCTACAGGCTACCACGAAGATAGCGT | Arm2; reverse | |
| P- InR 24 | CAGGTCATTTTCCATGGCCTTGACGA | Arm2; reverse | |
| P- InR 25 | AACCTTCCAGCGATCATCGCATGGAT | Arm2; reverse | √ |
| P- InR 26 | GGACGGCAGCCAGGCCATATTTTAAT | Arm2; reverse | √ |
| P- InR 27 | GACACATTTCGACAGGAGATGCAGCT | Arm2; reverse | √ |
| P- InR 28 | GAGTGCAATCCAGTCACACGGTAGAT | Arm2; reverse | √ |
| P- InR 31 | GGTGGTCATGGAGCTAATGAAGAAGG | Arm1; forward | √ |
| P- InR 32 | GGCGAGACCCAGTTTTCTGGATATCA | Arm1; forward | √ |
| P- InR 33 | GATCTACGATCCCAGTCCGAAATGTG | Arm1; forward | √ |
| P- InR 34 | GACTACTAGACGAGTCAATAGCCAGC | Arm1; forward | √ |
| P- InR 35 | GCCATCTGCACTGGAGTCTCATTCAT | Arm1; reverse | |
| P- InR 36 | GACCCACCGATGACATCATCTGGCAT | Arm1; reverse | |
| P- InR 37 | ACTGCCGCGAATGCATCTAGCTGATT | Arm1; reverse | √ |
| P- InR 38 | CACATTACCAGTCACTCCGATGCGAT | Arm1; reverse | √ |
| P- InR 64 | ACAAAGCATATTTATCCGCGTCC | Arm2; reverse | |
| P- InR 65 | CAACACCGAGTCCCCAGTACGG | Arm2; forward | |
| P- InR 66 | CCGTACTGGGGACTCGGTGTTG | Arm2; forward | |
| P- InR 67 | TATGTTCGACTTCCGTGAAATCGC | Arm2; reverse | |
| P- InR 68 | TTCAACAGCCTCCGCCGAGC | Arm1; forward | |
| P- InR 69 | GCAAACACCGAGCAATCTTACG | Arm1; reverse | |
| P- InR 70 | TGGCCGCCAAGAAGTTCGTC | Arm1; forward | |
| P- InR 71 | TGCTGGTACTCGGAGCATTG | Arm1; reverse | |
| P- InR 72 | CACCAACTTGACAAGTCACCC | Arm1; forward | |
| P- InR 73 | ATGAATGAGACTCCAGTGCAGATGGC | Arm1; forward | |
| P- InR 74 | GCGTTAGAGGCTCTAGTGCAT | Arm1; reverse | |
| P- InR 75 | TGTGACAGTCGCATTGTTGGGT | Arm2; reverse | |
| P- InR 76 | CATTGTGCAGCAGGTGCTGCTGA | Arm2; reverse | |
| P- InR 77 | GCGGGACGAAAGATGATTACCTAG | Arm2; forward | |

| | | |
|---|---|---|
| P- InR 78 | ACGATATGGCAGCAGCAGCAAC | Arm2; forward |
| P- InR 79 | TGTTGCTTTTGTTGCGGTGTGGC | Arm2; reverse |
| P- InR 80 | GAAGTTTTCGCGCTGTTGTCG | Arm2; reverse |

Figure S1. Partial sequence of Drosophila insulin-like receptor (dInR) aligned with human insulin receptor (hIR, HGVS annotation, with signal peptide) and insulin growth factor-1 receptor (hIGF1R). Drosophila sequence translation initiation site of wDah (parent wildtype allele of homologous recombination, GenBank accession MT_563159) and of Fernandez (NCBI Reference Sequence: NM_079712.6). **Red**: Drosophila InR homologous recombination substitutions corresponding to allele. **Blue**: substitution of the Exelisis dominant negative *InR* transgene P{UAS-InR.K1409A} (BDSC). **Gold**: hIGF1R polymorphisms of Suh {Suh, 2008 #5452} enriched in centenarians, and of the *C. elegans* longevity allele *daf-2*(e1370). Domains: distal portion of L2, proximal portion of FnIII-1, Kinase insert domain (KID), Activation loop (A-Loop) with conserved autophosphorylation tyrosine (**Y**). The proposed SH2 binding motif of the Drosophila KID is highlighted, with Tyr1477.

```
dInR      MFNMPRGVTKSKSKRGKIKMENDMAAAATTTACTLGHICVLCRQEMLLDTCCCRQAVEAV    60
          wDah TIS                                      Reference TIS

dInR      DSPASSEEAYSSSNSSSCQASSEISAEEVWFLSHDDIVLCRRPKFDEVETTGKKRDVKCS    120

dInR      GHQCSNECDDGSTKNNRQQRENFNIFSNCHNILRTLQSLLLLMFNCGIFNKRRRRQHQQQ    180

dInR      HHHHYQHHQHHHQQHLQRQQANVSYTKFLLLLQTLAAATTRLSLSPKNYKQQQQLQHNQQ    240

hIR       ---------------------------------------------------------MAT    3
hIGF1R    ---------------------------------------------------------MKS    3
dInR      LPRATPQQKQQEKDRHKCFHYKHNYSYSPGISLLLFILLANTLAIQAVVLPAHQQHLLHN    300
                                                                   :  .

hIR       GGRRGAAAAPLLVAVAALLLGAAGHLYP------------GEV-CPGMDIRNNLTRLHEL    50
hIGF1R    GSGGGSPT-S---LWGLLFLSAALSLWP----------TSGEICGPGIDIRNDYQQLKRL    49
dInR      DIADGLDK-------TALSVSGTQSRWTRSESNPTMRLSQNVKPCKSMDIRNMVSHFNQL    353
          .   *            * :..:   :         .     .:****   :::.*

hIR       ENCSVIEGHLQILLMFKTRPEDFRDLSFPKLIMITDYLLLFRVYGLESLKDLFPNLTVIR    110
hIGF1R    ENCTVIEGYLHILLIS--KAEDYRSYRFPKLTVITEYLLLFRVAGLESLGDLFPNLTVIR    107
dInR      ENCTVIEGFLLIDLINDASP---LNRSFPKLTEVTDYIIIYRVTGLHSLSKIFPNLSVIR    410
           ***:****.* * *:    ^     .  ****  :*:*::::** **.** .:****:***
                        hIGF1R A67T
//= = = = = = = = = = = = = = = = = = = = = = = = = = = = = = = = //

hIR       YLKIRRSYALVSLSFFRKLRLIRGETL-EIGNYSFYALDNQNLRQLWDWSKHNLTITQGK    452
hIGF1R    YVKIRHSHALVSLSFLKNLRLILGEEQ-LEGNYSFYVLDNQNLQQLWDWDHRNLTIKAGK    445
dInR      SLMVHLTYGLKSLKFFQSLTEISGDPPMDADKYALYVLDNRDLDELWGPNQ-TVFIRKGG    759
           : :: ::.* **.*::.*  * *:      .:*::*.***::* :**. .:^.: *  *
                                                     hIGF1R R437H

                                  L2
hIR       LFFHYNPKLCLSEIHKMEEVSGTKG-RQERNDIALKTNGDQASCENELLKFSYIRTSFDK    511
hIGF1R    MYFAFNPKLCVSEIYRMEEVTGTKG-RQSKGDINTRNNGERASCESDVLHFTSTTTSKNR    504
dInR      VFFHFNPKLCVSTINQLLPMLASKPKFFEKSDVGADSNGNRGSCGTAVLNVTLQSVGANS    819
          ::* :*****:* * ::  : .:*    .:.*:   .**::.** . :*:^:    .. :
                                                      dInR^e19 V810D

                                FnIII-1
hIR       ILLRWE-------PY-------WPPDFRDLLGFMLFYKEAPYQNVTEFDGQDACGSNSWT    557
hIGF1R    IIITWH-------RY-------RPPDYRDLISFTVYYKEAPFKNVTEYDGQDACGSNSWN    550
dInR      AMLNVTTKVEIGEPQKPSNATIVFKDPRAFIGFVFYHMIDPYGNSTKSS-DDPC-DDRWK    877
           ::                        * * ::.* .::   *: * *: . :* * .^ *.
                                                        hIGF1R N547H
//= = = = = = = = = = = = = = = = = = = = = = = = = = = = = = = = //

hIR       GMVYEGNARDIIKGEAETRVAVKTVNESASLRERIEFLNEASVMKGFTCHHVVRLLGVVS    1094
hIGF1R    GMVYEGVAKGVVKDEPETRVAIKTVNEAASMRERIEFLNEASVMKEFNCHHVVRLLGVVS    1070
dInR      GMVYEGILKSFPPNGVDRECAIKTVNENATDRERTNFLSEASVMKEFDTYHVVRLLGVCS    1441
          **^***   :..   .   :  . *:***** *: *** :**.****** *   :******** *
          dInR^246 V1384M    UAS-dInR-DN K1409A

                                              KID
hIR       KGQPTLVVMELMAHGDLKSYLRSLRPEAENNP------------GRPPPTLQEMIQMAAE    1142
hIGF1R    QGQPTLVIMELMTRGDLKSYLRSLRPEMENNP------------VLAPPSLSKMIQMAGE    1118
dInR      RGQPALVVMELMKKGDLKSYLRAHRPEERDEAMMTYLNRIGVTGNVQPPTYGRIYQMAIE    1501
          :***:**:****  :********: ^**  .::                **:   .: *** *
                            dInR^353 R1466C

                                                     A-Loop
hIR       IADGMAYLNAKKFVHRDLAARNCMVAHDFTVKIGDFGMTRDIYETDYYRKGGKGLLPVRW    1202
hIGF1R    IADGMAYLNANKFVHRDLAARNCMVAEDFTVKIGDFGMTRDIYETDYYRKGGKGLLPVRW    1178
dInR      IADGMAYLAAKKFVHRDLAARNCMVADDLTVKIGDFGMTRDIYETDYYRKGTKGLLPVRW    1561
          ******** *:***************.*:************^********* ********
                                                dInR^74 I1543F

hIR       MAPESLKDGVFTTSSDMWSFGVVLWEITSLAEQPYQGLSNEQVLKFVMDGGYLDQPDNCP    1262
hIGF1R    MSPESLKDGVFTTYSDVWSFGVVLWEIATLAEQPYQGLSNEQVLRFVMEGGLLDKPDNCP    1238
dInR      MPPESLRDGVYSSASDVFSFGVVLWEMATLAAQPYQGLSNEQVLRYVIDGGVMERPENCP    1621
          * ****:***::: **::********:::** *^**^*******::*::**  :::*:***
                           daf2(e1370)P1465S   dInR^211 G1598R
```

**Supplemental Figure S2. Distribution of eclosion time.** Six genotypes, emerging males and females from from day of egg deposition (0). Sum of two replicate vials. $F_1$ eggs after 24 hours mating, from crosses of 4 males and 4 females: *InR*(mutant allele 1)/TM6*Sb* x *InR*(mutant allele 2)/TM6*Sb*. Emerging adults with marker *Sb* are TM6 heterozygotes with either mutant *InR* allele. Flies without the marker are trans-heterozygote mutant *InR* genotypes.

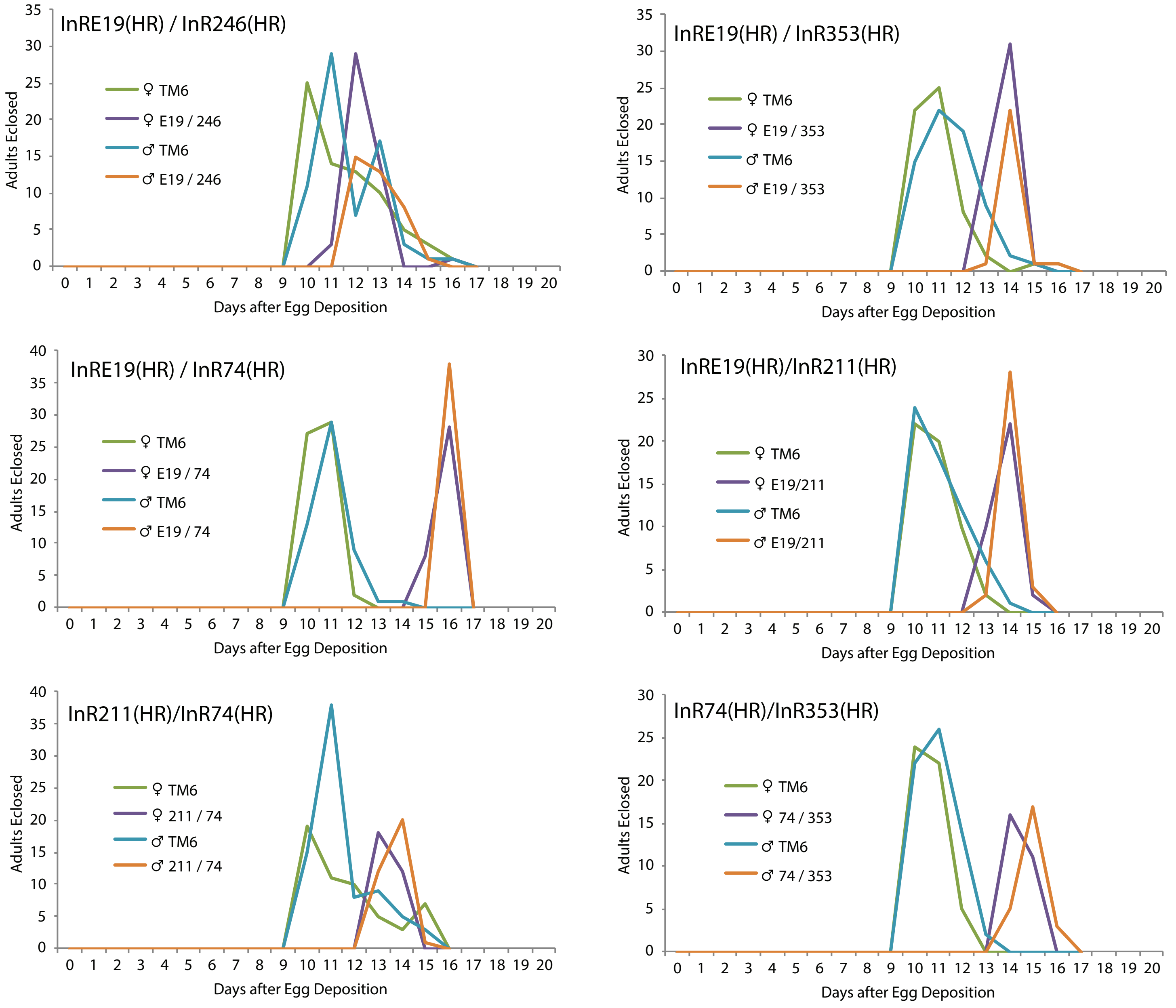

**Fig S3. Master DNA alignment HR alleles with wildtypes *w*DAH and 29B(HR)**

```
DAH         agcgggacgaaagatgattacctagcgggggtgagagagacccaaaaagcgaaagcagaa
29B         agcgggacgaaagatgattacctagcgggggtgagagagacccaaaaagcgaaagcagaa
e19         agcgggacgaaagatgattacctagcgggggtgagagagacccaaaaagcgaaagcagaa
74e19       agcgggacgaaagatgattacctagcgggggtgagagagacccaaaaagcgaaagcagaa
74          agcgggacgaaagatgattacctagcgggggtgagagagacccaaaaagcgaaagcagaa
211         agcgggacgaaagatgattacctagcgggggtgagagagacccaaaaagcgaaagcagaa
246         agcgggacgaaagatgattacctagcgggggtgagagagacccaaaaagcgaaagcagaa
353         agcgggacgaaagatgattacctagcgggggtgagagagacccaaaaagcgaaagcagaa
            ************************************************************

DAH         ctcgtggaaagctgaaaatttgaggaaacaatttcataatacgaatagaaacaccgcaat
29B         ctcgtggaaagctgaaaatttgaggaaacaatttcataatacgaatagaaacaccgcaat
e19         ctcgtggaaagctgaaaatttgaggaaacaatttcataatacgaatagaaacaccgcaat
74e19       ctcgtggaaagctgaaaatttgaggaaacaatttcataatacgaatagaaacaccgcaat
74          ctcgtggaaagctgaaaatttgaggaaacaatttcataatacgaatagaaacaccgcaat
211         ctcgtggaaagctgaaaatttgaggaaacaatttcataatacgaatagaaacaccgcaat
246         ctcgtggaaagctgaaaatttgaggaaacaatttcataatacgaatagaaacaccgcaat
353         ctcgtggaaagctgaaaatttgaggaaacaatttcataatacgaatagaaacaccgcaat
            ************************************************************

DAH         ATGTTCAATATGCCACGGGGAGTGACAAAAAGTAAATCCAAGCGTGGGAAAATTAAGATG     60
29B         ATGTTCAATATGCCACGGGGAGTGACAAAAAGTAAATCCAAGCGTGGGAAAATTAAGATG     60
e19         ATGTTCAATATGCCACGGGGAGTGACAAAAAGTAAATCCAAGCGTGGGAAAATTAAGATG     60
74e19       ATGTTCAATATGCCACGGGGAGTGACAAAAAGTAAATCCAAGCGTGGGAAAATTAAGATG     60
74          ATGTTCAATATGCCACGGGGAGTGACAAAAAGTAAATCCAAGCGTGGGAAAATTAAGATG     60
211         ATGTTCAATATGCCACGGGGAGTGACAAAAAGTAAATCCAAGCGTGGGAAAATTAAGATG     60
246         ATGTTCAATATGCCACGGGGAGTGACAAAAAGTAAATCCAAGCGTGGGAAAATTAAGATG     60
353         ATGTTCAATATGCCACGGGGAGTGACAAAAAGTAAATCCAAGCGTGGGAAAATTAAGATG     60
            ************************************************************

DAH         GAAAACGATATGGCAGCAGCAGCAACAACAACAGCCTGCACGCTTGGACACATTTGTGTT     120
29B         GAAAACGATATGGCAGCAGCAGCAACAACAACAGCCTGCACGCTTGGACACATTTGTGTT     120
e19         GAAAACGATATGGCAGCAGCAGCAACAACAACAGCCTGCACGCTTGGACACATTTGTGTT     120
74e19       GAAAACGATATGGCAGCAGCAGCAACAACAACAGCCTGCACGCTTGGACACATTTGTGTT     120
74          GAAAACGATATGGCAGCAGCAGCAACAACAACAGCCTGCACGCTTGGACACATTTGTGTT     120
211         GAAAACGATATGGCAGCAGCAGCAACAACAACAGCCTGCACGCTTGGACACATTTGTGTT     120
246         GAAAACGATATGGCAGCAGCAGCAACAACAACAGCCTGCACGCTTGGACACATTTGTGTT     120
353         GAAAACGATATGGCAGCAGCAGCAACAACAACAGCCTGCACGCTTGGACACATTTGTGTT     120
            ************************************************************

DAH         TTGTGCCGGCAAGAAATGTTGCTGGATACATGTTGCTGCCGGCAAGCAGTAGAAGCAGTT     180
29B         TTGTGCCGGCAAGAAATGTTGCTGGATACATGTTGCTGCCGGCAAGCAGTAGAAGCAGTT     180
e19         TTGTGCCGGCAAGAAATGTTGCTGGATACATGTTGCTGCCGGCAAGCAGTAGAAGCAGTT     180
74e19       TTGTGCCGGCAAGAAATGTTGCTGGATACATGTTGCTGCCGGCAAGCAGTAGAAGCAGTT     180
74          TTGTGCCGGCAAGAAATGTTGCTGGATACATGTTGCTGCCGGCAAGCAGTAGAAGCAGTT     180
211         TTGTGCCGGCAAGAAATGTTGCTGGATACATGTTGCTGCCGGCAAGCAGTAGAAGCAGTT     180
246         TTGTGCCGGCAAGAAATGTTGCTGGATACATGTTGCTGCCGGCAAGCAGTAGAAGCAGTT     180
353         TTGTGCCGGCAAGAAATGTTGCTGGATACATGTTGCTGCCGGCAAGCAGTAGAAGCAGTT     180
            ************************************************************
```

### Supplemental Methods and Data

#### 1. Drosophila culture, stocks, demography

**Culture**

Flies were reared and aged at 25°C, 40% RH, 12L:12D on standard food media: cornmeal (5.2%), sugar (11.0%), autolyzed yeast (2.5%; SAF brand, Lesaffre Yeast Corp., Milwaukee, WI, USA.), agar (0.79%) (w/v in 100 mL water) with 0.2% Tegosep (methyl4-hydroxybenzoate, Sigma, St Louis, MO, USA).

Oviposition plates. 5 cm dishes with grape juice media and excess live yeast were affixed to the food tube port of demography cages. Plates were changed twice daily, with total eggs summed each day. Grape juice plate media: 25 g Bacto-Agar (BD cat # 214010), 250 ml grape juice (Welsh concentrate), 750 ml water and 15 ml 20% Tegosep.

**Lifetable demography**

Cages for lifetable demography were made from 1 L clear food service containers with a ventilated lid, a gasket covered aperture to provide access to remove dead flies and a port near the cage bottom that opened to a plastic tube affixed a standard glass media vial with 3 ml of standard Drosophila diet. Each two days, dead flies were removed by aspiration and counted, and fresh food vials were provided.

#### 2. Ends out homologous recombination

**Reagents**

| |
|---|
| Genomic DNA extraction: Promega Wizard SV Genomic Purification System. Promega, cat # A2361 |
| PCR: Phusion High-Fidelity PCR Master Mis with GC Buffer. New England Biolabs, cat # M0532 |
| Cloning: Zero Blunt Topo PCR cloning kit, Invitrogen, cat # 45-0245 |
| pW25.2 (Drosophila Genomics Resource Center) |
| Bacteria Transformation: One Shot Top10 electrocompetent cells (Invitrogen), cat # C404052 |
| Plasmid DNA prep: QiaPrep Spin miniprep kit, Qiagen, cat # 27104 |
| Gel purification: QIAquick Gel Extraction Kit. Qiagen, cat # 28740 |
| Site-direct mutagenesis: GeneArt site direct mutagenesis system, Invitrogen, cat #A13282, using AccuPrime Pfx DNA polymerase, Invitrogen, cat #12344-024 |
| Primer synthesis: Integrated DNA Technologies (IDT, IA, USA) |
| Sequencing: Genewiz (South Plainfield, NJ, USA) |
| Restriction Enzymes: New England Biolabs |
| Embryo Injection: Genetic Services, Inc. (Sudbury, MA) |

**Drosophila Stocks for homologous recombination**

| *Genotype* | *Stock* |
|---|---|
| $y^1w^*$; $P\{ry^{+t7.2} = 70FLP\}11\ P\{v^{+t1.8} = 70I\text{-}SceI\}2B\ noc^{Sco}$ / $CyO, S^2$ | BSC 6934 |
| $y^{d2}\ w^{1118}\ P\{ry^{+t7.2}$=ey-FLP.N}2 | BSC 5580 |
| $y^1\ w^{67c23}$; $sna^{Sco}$/CyO, $P\{w^{+mC}$=Crew}DH1 | BSC 1092 |
| *wDahomey* | L. Partridge (UCL, UK) |

All lines backcrossed to $w^{Dah}$ for six generations.

**Targeting arms for homologous recombination**

*InR* gene replacement to produce single nucleotide substitution followed Staber *et al* (Staber CJ, Gell S, Jepson JE, Reenan RA. 2011. Perturbing A-to-I RNA Editing Using Genetics and Homologous

```
DAH      GACAGCCCCGCAAGCAGTGAAGAAGCGTATAGCAGTAGCAACAGCAGCAGCTGTCAAGCA    240
29B      GACAGCCCCGCAAGCAGTGAAGAAGCGTATAGCAGTAGCAACAGCAGCAGCTGTCAAGCA    240
e19      GACAGCCCCGCAAGCAGTGAAGAAGCGTATAGCAGTAGCAACAGCAGCAGCTGTCAAGCA    240
74e19    GACAGCCCCGCAAGCAGTGAAGAAGCGTATAGCAGTAGCAACAGCAGCAGCTGTCAAGCA    240
74       GACAGCCCCGCAAGCAGTGAAGAAGCGTATAGCAGTAGCAACAGCAGCAGCTGTCAAGCA    240
211      GACAGCCCCGCAAGCAGTGAAGAAGCGTATAGCAGTAGCAACAGCAGCAGCTGTCAAGCA    240
246      GACAGCCCCGCAAGCAGTGAAGAAGCGTATAGCAGTAGCAACAGCAGCAGCTGTCAAGCA    240
353      GACAGCCCCGCAAGCAGTGAAGAAGCGTATAGCAGTAGCAACAGCAGCAGCTGTCAAGCA    240
         ************************************************************

DAH      AGCAGTGAAATCAGTGCGGAGGAGGTCTGGTTTCTCAGTCATGATGATATCGTACTGTGC    300
29B      AGCAGTGAAATCAGTGCGGAGGAGGTCTGGTTTCTCAGTCATGATGATATCGTACTGTGC    300
e19      AGCAGTGAAATCAGTGCGGAGGAGGTCTGGTTTCTCAGTCATGATGATATCGTACTGTGC    300
74e19    AGCAGTGAAATCAGTGCGGAGGAGGTCTGGTTTCTCAGTCATGATGATATCGTACTGTGC    300
74       AGCAGTGAAATCAGTGCGGAGGAGGTCTGGTTTCTCAGTCATGATGATATCGTACTGTGC    300
211      AGCAGTGAAATCAGTGCGGAGGAGGTCTGGTTTCTCAGTCATGATGATATCGTACTGTGC    300
246      AGCAGTGAAATCAGTGCGGAGGAGGTCTGGTTTCTCAGTCATGATGATATCGTACTGTGC    300
353      AGCAGTGAAATCAGTGCGGAGGAGGTCTGGTTTCTCAGTCATGATGATATCGTACTGTGC    300
         ************************************************************

DAH      CGCAGACCAAAATTTGACGAAGTGGAGACGACGGGTAAAAAGAGGGACGTTAAATGCAGC    360
29B      CGCAGACCAAAATTTGACGAAGTGGAGACGACGGGTAAAAAGAGGGACGTTAAATGCAGC    360
e19      CGCAGACCAAAATTTGACGAAGTGGAGACGACGGGTAAAAAGAGGGACGTTAAATGCAGC    360
74e19    CGCAGACCAAAATTTGACGAAGTGGAGACGACGGGTAAAAAGAGGGACGTTAAATGCAGC    360
74       CGCAGACCAAAATTTGACGAAGTGGAGACGACGGGTAAAAAGAGGGACGTTAAATGCAGC    360
211      CGCAGACCAAAATTTGACGAAGTGGAGACGACGGGTAAAAAGAGGGACGTTAAATGCAGC    360
246      CGCAGACCAAAATTTGACGAAGTGGAGACGACGGGTAAAAAGAGGGACGTTAAATGCAGC    360
353      CGCAGACCAAAATTTGACGAAGTGGAGACGACGGGTAAAAAGAGGGACGTTAAATGCAGC    360
         ************************************************************

DAH      GGGCATCAGTGCAGCAATGAATGCGACGATGGCAGCACGAAAAACAATCGACAACAGCGC    420
29B      GGGCATCAGTGCAGCAATGAATGCGACGATGGCAGCACGAAAAACAATCGACAACAGCGC    420
e19      GGGCATCAGTGCAGCAATGAATGCGACGATGGCAGCACGAAAAACAATCGACAACAGCGC    420
74e19    GGGCATCAGTGCAGCAATGAATGCGACGATGGCAGCACGAAAAACAATCGACAACAGCGC    420
74       GGGCATCAGTGCAGCAATGAATGCGACGATGGCAGCACGAAAAACAATCGACAACAGCGC    420
211      GGGCATCAGTGCAGCAATGAATGCGACGATGGCAGCACGAAAAACAATCGACAACAGCGC    420
246      GGGCATCAGTGCAGCAATGAATGCGACGATGGCAGCACGAAAAACAATCGACAACAGCGC    420
353      GGGCATCAGTGCAGCAATGAATGCGACGATGGCAGCACGAAAAACAATCGACAACAGCGC    420
         ************************************************************

DAH      GAAAACTTCAATATCTTTAGCAACTGTCACAATATTTTGCGAACATTGCAATCGCTGCTG    480
29B      GAAAACTTCAATATCTTTAGCAACTGTCACAATATTTTGCGAACATTGCAATCGCTGCTG    480
e19      GAAAACTTCAATATCTTTAGCAACTGTCACAATATTTTGCGAACATTGCAATCGCTGCTG    480
74e19    GAAAACTTCAATATCTTTAGCAACTGTCACAATATTTTGCGAACATTGCAATCGCTGCTG    480
74       GAAAACTTCAATATCTTTAGCAACTGTCACAATATTTTGCGAACATTGCAATCGCTGCTG    480
211      GAAAACTTCAATATCTTTAGCAACTGTCACAATATTTTGCGAACATTGCAATCGCTGCTG    480
246      GAAAACTTCAATATCTTTAGCAACTGTCACAATATTTTGCGAACATTGCAATCGCTGCTG    480
353      GAAAACTTCAATATCTTTAGCAACTGTCACAATATTTTGCGAACATTGCAATCGCTGCTG    480
         ************************************************************

DAH      CTGCTCATGTTCAATTGCGGCATTTTCAACAAGCGACGCAGGCGGCAGCATCAGCAGCAG    540
29B      CTGCTCATGTTCAATTGCGGCATTTTCAACAAGCGACGCAGGCGGCAGCATCAGCAGCAG    540
e19      CTGCTCATGTTCAATTGCGGCATTTTCAACAAGCGACGCAGGCGGCAGCATCAGCAGCAG    540
74e19    CTGCTCATGTTCAATTGCGGCATTTTCAACAAGCGACGCAGGCGGCAGCATCAGCAGCAG    540
74       CTGCTCATGTTCAATTGCGGCATTTTCAACAAGCGACGCAGGCGGCAGCATCAGCAGCAG    540
211      CTGCTCATGTTCAATTGCGGCATTTTCAACAAGCGACGCAGGCGGCAGCATCAGCAGCAG    540
246      CTGCTCATGTTCAATTGCGGCATTTTCAACAAGCGACGCAGGCGGCAGCATCAGCAGCAG    540
353      CTGCTCATGTTCAATTGCGGCATTTTCAACAAGCGACGCAGGCGGCAGCATCAGCAGCAG    540
         ************************************************************
```

```
DAH      CATCATCATCATTATCAGCATCATCAGCATCATCATCAGCAGCATCTTCAGCGGCAGCAA   600
29B      CATCATCATCATTATCAGCATCATCAGCATCATCATCAGCAGCATCTTCAGCGGCAGCAA   600
e19      CATCATCATCATTATCAGCATCATCAGCATCATCATCAGCAGCATCTTCAGCGGCAGCAA   600
74e19    CATCATCATCATTATCAGCATCATCAGCATCATCATCAGCAGCATCTTCAGCGGCAGCAA   600
74       CATCATCATCATTATCAGCATCATCAGCATCATCATCAGCAGCATCTTCAGCGGCAGCAA   600
211      CATCATCATCATTATCAGCATCATCAGCATCATCATCAGCAGCATCTTCAGCGGCAGCAA   600
246      CATCATCATCATTATCAGCATCATCAGCATCATCATCAGCAGCATCTTCAGCGGCAGCAA   600
353      CATCATCATCATTATCAGCATCATCAGCATCATCATCAGCAGCATCTTCAGCGGCAGCAA   600
         ************************************************************

DAH      GCCAATGTTAGTTACACAAAATTCCTATTGCTGCTACAAACACTGGCAGCAGCAACCACA   660
29B      GCCAATGTTAGTTACACAAAATTCCTATTGCTGCTACAAACACTGGCAGCAGCAACCACA   660
e19      GCCAATGTTAGTTACACAAAATTCCTATTGCTGCTACAAACACTGGCAGCAGCAACCACA   660
74e19    GCCAATGTTAGTTACACAAAATTCCTATTGCTGCTACAAACACTGGCAGCAGCAACCACA   660
74       GCCAATGTTAGTTACACAAAATTCCTATTGCTGCTACAAACACTGGCAGCAGCAACCACA   660
211      GCCAATGTTAGTTACACAAAATTCCTATTGCTGCTACAAACACTGGCAGCAGCAACCACA   660
246      GCCAATGTTAGTTACACAAAATTCCTATTGCTGCTACAAACACTGGCAGCAGCAACCACA   660
353      GCCAATGTTAGTTACACAAAATTCCTATTGCTGCTACAAACACTGGCAGCAGCAACCACA   660
         ************************************************************

DAH      AGACTGAGTTTAAGCCCTAAAAACTACAAACAACAACAACAACTACAGCATAACCAACAG   720
29B      AGACTGAGTTTAAGCCCTAAAAACTACAAACAACAACAACAACTACAGCATAACCAACAG   720
e19      AGACTGAGTTTAAGCCCTAAAAACTACAAACAACAACAACAACTACAGCATAACCAACAG   720
74e19    AGACTGAGTTTAAGCCCTAAAAACTACAAACAACAACAACAACTACAGCATAACCAACAG   720
74       AGACTGAGTTTAAGCCCTAAAAACTACAAACAACAACAACAACTACAGCATAACCAACAG   720
211      AGACTGAGTTTAAGCCCTAAAAACTACAAACAACAACAACAACTACAGCATAACCAACAG   720
246      AGACTGAGTTTAAGCCCTAAAAACTACAAACAACAACAACAACTACAGCATAACCAACAG   720
353      AGACTGAGTTTAAGCCCTAAAAACTACAAACAACAACAACAACTACAGCATAACCAACAG   720
         ************************************************************

DAH      CTGCCACGTGCCACACCGCAACAAAAGCAACAAGAGAAAGATAGGCATAAGTGCTTTCAC   780
29B      CTGCCACGTGCCACACCGCAACAAAAGCAACAAGAGAAAGATAGGCATAAGTGCTTTCAC   780
e19      CTGCCACGTGCCACACCGCAACAAAAGCAACAAGAGAAAGATAGGCATAAGTGCTTTCAC   780
74e19    CTGCCACGTGCCACACCGCAACAAAAGCAACAAGAGAAAGATAGGCATAAGTGCTTTCAC   780
74       CTGCCACGTGCCACACCGCAACAAAAGCAACAAGAGAAAGATAGGCATAAGTGCTTTCAC   780
211      CTGCCACGTGCCACACCGCAACAAAAGCAACAAGAGAAAGATAGGCATAAGTGCTTTCAC   780
246      CTGCCACGTGCCACACCGCAACAAAAGCAACAAGAGAAAGATAGGCATAAGTGCTTTCAC   780
353      CTGCCACGTGCCACACCGCAACAAAAGCAACAAGAGAAAGATAGGCATAAGTGCTTTCAC   780
         ************************************************************

DAH      TACAAGCACAATTACTCTTACTCGCCTGGCATTAGCCTTCTACTCTTTATCCTACTGGCC   840
29B      TACAAGCACAATTACTCTTACTCGCCTGGCATTAGCCTTCTACTCTTTATCCTACTGGCC   840
e19      TACAAGCACAATTACTCTTACTCGCCTGGCATTAGCCTTCTACTCTTTATCCTACTGGCC   840
74e19    TACAAGCACAATTACTCTTACTCGCCTGGCATTAGCCTTCTACTCTTTATCCTACTGGCC   840
74       TACAAGCACAATTACTCTTACTCGCCTGGCATTAGCCTTCTACTCTTTATCCTACTGGCC   840
211      TACAAGCACAATTACTCTTACTCGCCTGGCATTAGCCTTCTACTCTTTATCCTACTGGCC   840
246      TACAAGCACAATTACTCTTACTCGCCTGGCATTAGCCTTCTACTCTTTATCCTACTGGCC   840
353      TACAAGCACAATTACTCTTACTCGCCTGGCATTAGCCTTCTACTCTTTATCCTACTGGCC   840
         ************************************************************

DAH      AACACATTGGCCATCCAAGCGGTCGTGTTGCCAGCACATCAGCAGCACCTGCTGCACAAT   900
29B      AACACATTGGCCATCCAAGCGGTCGTGTTGCCAGCACATCAGCAGCACCTGCTGCACAAT   900
e19      AACACATTGGCCATCCAAGCGGTCGTGTTGCCAGCACATCAGCAGCACCTGCTGCACAAT   900
74e19    AACACATTGGCCATCCAAGCGGTCGTGTTGCCAGCACATCAGCAGCACCTGCTGCACAAT   900
74       AACACATTGGCCATCCAAGCGGTCGTGTTGCCAGCACATCAGCAGCACCTGCTGCACAAT   900
211      AACACATTGGCCATCCAAGCGGTCGTGTTGCCAGCACATCAGCAGCACCTGCTGCACAAT   900
246      AACACATTGGCCATCCAAGCGGTCGTGTTGCCAGCACATCAGCAGCACCTGCTGCACAAT   900
353      AACACATTGGCCATCCAAGCGGTCGTGTTGCCAGCACATCAGCAGCACCTGCTGCACAAT   900
         ************************************************************
```

```
DAH      GATATAGCCGATGGACTGGATAAAACAGCGCTTTCGGTGTCGGGGACGCAATCGCGATGG   960
29B      GATATAGCCGATGGACTGGATAAAACAGCGCTTTCGGTGTCGGGGACGCAATCGCGATGG   960
e19      GATATAGCCGATGGACTGGATAAAACAGCGCTTTCGGTGTCGGGGACGCAATCGCGATGG   960
74e19    GATATAGCCGATGGACTGGATAAAACAGCGCTTTCGGTGTCGGGGACGCAATCGCGATGG   960
74       GATATAGCCGATGGACTGGATAAAACAGCGCTTTCGGTGTCGGGGACGCAATCGCGATGG   960
211      GATATAGCCGATGGACTGGATAAAACAGCGCTTTCGGTGTCGGGGACGCAATCGCGATGG   960
246      GATATAGCCGATGGACTGGATAAAACAGCGCTTTCGGTGTCGGGGACGCAATCGCGATGG   960
353      GATATAGCCGATGGACTGGATAAAACAGCGCTTTCGGTGTCGGGGACGCAATCGCGATGG   960
         ************************************************************

DAH      ACAAGGAGCGAATCAAACCCAACAATGCGACTGTCACAAAATGTAAAACgtatgtattac   1020
29B      ACAAGGAGCGAATCAAACCCAACAATGCGACTGTCACAAAATGTAAAACgtatgtattac   1020
e19      ACAAGGAGCGAATCAAACCCAACAATGCGACTGTCACAAAATGTAAAACgtatgtattac   1020
74e19    ACAAGGAGCGAATCAAACCCAACAATGCGACTGTCACAAAATGTAAAACgtatgtattac   1020
74       ACAAGGAGCGAATCAAACCCAACAATGCGACTGTCACAAAATGTAAAACgtatgtattac   1020
211      ACAAGGAGCGAATCAAACCCAACAATGCGACTGTCACAAAATGTAAAACgtatgtattac   1020
246      ACAAGGAGCGAATCAAACCCAACAATGCGACTGTCACAAAATGTAAAACgtatgtattac   1020
353      ACAAGGAGCGAATCAAACCCAACAATGCGACTGTCACAAAATGTAAAACgtatgtattac   1020
         ************************************************************

DAH      ctatgatctaccttgcaagttaaaggaacattcccatttgcacgagactgacagatacta   1080
29B      ctatgatctaccttgcaagttaaaggaacattcccatttgcacgagactgacagatacta   1080
e19      ctatgatctaccttgcaagttaaaggaacattcccatttgcacgagactgacagatacta   1080
74e19    ctatgatctaccttgcaagttaaaggaacattcccatttgcacgagactgacagatacta   1080
74       ctatgatctaccttgcaagttaaaggaacattcccatttgcacgagactgacagatacta   1080
211      ctatgatctaccttgcaagttaaaggaacattcccatttgcacgagactgacagatacta   1080
246      ctatgatctaccttgcaagttaaaggaacattcccatttgcacgagactgacagatacta   1080
353      ctatgatctaccttgcaagttaaaggaacattcccatttgcacgagactgacagatacta   1080
         ************************************************************

DAH      atagcttttccattaacaccttatagCTTGCAAATCCATGGACATCAGGAACATGGTGTC   1140
29B      atagcttttccattaacaccttatagCTTGCAAATCCATGGACATCAGGAACATGGTGTC   1140
e19      atagcttttccattaacaccttatagCTTGCAAATCCATGGACATCAGGAACATGGTGTC   1140
74e19    atagcttttccattaacaccttatagCTTGCAAATCCATGGACATCAGGAACATGGTGTC   1140
74       atagcttttccattaacaccttatagCTTGCAAATCCATGGACATCAGGAACATGGTGTC   1140
211      atagcttttccattaacaccttatagCTTGCAAATCCATGGACATCAGGAACATGGTGTC   1140
246      atagcttttccattaacaccttatagCTTGCAAATCCATGGACATCAGGAACATGGTGTC   1140
353      atagcttttccattaacaccttatagCTTGCAAATCCATGGACATCAGGAACATGGTGTC   1140
         ************************************************************

DAH      GCACTTCAATCAGCTGGAGAACTGCACGGTCATCGAGGGCTTCCTGCTGATCGATTTGAT   1200
29B      GCACTTCAATCAGCTGGAGAACTGCACGGTCATCGAGGGCTTCCTGCTGATCGATTTGAT   1200
e19      GCACTTCAATCAGCTGGAGAACTGCACGGTCATCGAGGGCTTCCTGCTGATCGATTTGAT   1200
74e19    GCACTTCAATCAGCTGGAGAACTGCACGGTCATCGAGGGCTTCCTGCTGATCGATTTGAT   1200
74       GCACTTCAATCAGCTGGAGAACTGCACGGTCATCGAGGGCTTCCTGCTGATCGATTTGAT   1200
211      GCACTTCAATCAGCTGGAGAACTGCACGGTCATCGAGGGCTTCCTGCTGATCGATTTGAT   1200
246      GCACTTCAATCAGCTGGAGAACTGCACGGTCATCGAGGGCTTCCTGCTGATCGATTTGAT   1200
353      GCACTTCAATCAGCTGGAGAACTGCACGGTCATCGAGGGCTTCCTGCTGATCGATTTGAT   1200
         ************************************************************

DAH      AAACGACGCCAGCCCTCTGAACAGAAGCTTTCCAAAACTGACCGAGGTCACAGATTATAT   1260
29B      AAACGACGCCAGCCCTCTGAACAGAAGCTTTCCAAAACTGACCGAGGTCACAGATTATAT   1260
e19      AAACGACGCCAGCCCTCTGAACAGAAGCTTTCCAAAACTGACCGAGGTCACAGATTATAT   1260
74e19    AAACGACGCCAGCCCTCTGAACAGAAGCTTTCCAAAACTGACCGAGGTCACAGATTATAT   1260
74       AAACGACGCCAGCCCTCTGAACAGAAGCTTTCCAAAACTGACCGAGGTCACAGATTATAT   1260
211      AAACGACGCCAGCCCTCTGAACAGAAGCTTTCCAAAACTGACCGAGGTCACAGATTATAT   1260
246      AAACGACGCCAGCCCTCTGAACAGAAGCTTTCCAAAACTGACCGAGGTCACAGATTATAT   1260
353      AAACGACGCCAGCCCTCTGAACAGAAGCTTTCCAAAACTGACCGAGGTCACAGATTATAT   1260
         ************************************************************
```

```
DAH       CATAATCTACCGTGTGACTGGATTGCACTCGCTGTCAAAGATCTTTCCCAATCTGAGCGT    1320
29B       CATAATCTACCGTGTGACTGGATTGCACTCGCTGTCAAAGATCTTTCCCAATCTGAGCGT    1320
e19       CATAATCTACCGTGTGACTGGATTGCACTCGCTGTCAAAGATCTTTCCCAATCTGAGCGT    1320
74e19     CATAATCTACCGTGTGACTGGATTGCACTCGCTGTCAAAGATCTTTCCCAATCTGAGCGT    1320
74        CATAATCTACCGTGTGACTGGATTGCACTCGCTGTCAAAGATCTTTCCCAATCTGAGCGT    1320
211       CATAATCTACCGTGTGACTGGATTGCACTCGCTGTCAAAGATCTTTCCCAATCTGAGCGT    1320
246       CATAATCTACCGTGTGACTGGATTGCACTCGCTGTCAAAGATCTTTCCCAATCTGAGCGT    1320
353       CATAATCTACCGTGTGACTGGATTGCACTCGCTGTCAAAGATCTTTCCCAATCTGAGCGT    1320
          ************************************************************

DAH       CATTAGGGGAAACAAGCTGTTCGACGGATATGCCTTGGTCGTCTACTCGAATTTCGACCT    1380
29B       CATTAGGGGAAACAAGCTGTTCGACGGATATGCCTTGGTCGTCTACTCGAATTTCGACCT    1380
e19       CATTAGGGGAAACAAGCTGTTCGACGGATATGCCTTGGTCGTCTACTCGAATTTCGACCT    1380
74e19     CATTAGGGGAAACAAGCTGTTCGACGGATATGCCTTGGTCGTCTACTCGAATTTCGACCT    1380
74        CATTAGGGGAAACAAGCTGTTCGACGGATATGCCTTGGTCGTCTACTCGAATTTCGACCT    1380
211       CATTAGGGGAAACAAGCTGTTCGACGGATATGCCTTGGTCGTCTACTCGAATTTCGACCT    1380
246       CATTAGGGGAAACAAGCTGTTCGACGGATATGCCTTGGTCGTCTACTCGAATTTCGACCT    1380
353       CATTAGGGGAAACAAGCTGTTCGACGGATATGCCTTGGTCGTCTACTCGAATTTCGACCT    1380
          ************************************************************

DAH       CATGGATTTGGGACTTCACAAGCTACGATCCATAACCAGAGGCGGTGTGCGGATTGAGAA    1440
29B       CATGGATTTGGGACTTCACAAGCTACGATCCATAACCAGAGGCGGTGTGCGGATTGAGAA    1440
e19       CATGGATTTGGGACTTCACAAGCTACGATCCATAACCAGAGGCGGTGTGCGGATTGAGAA    1440
74e19     CATGGATTTGGGACTTCACAAGCTACGATCCATAACCAGAGGCGGTGTGCGGATTGAGAA    1440
74        CATGGATTTGGGACTTCACAAGCTACGATCCATAACCAGAGGCGGTGTGCGGATTGAGAA    1440
211       CATGGATTTGGGACTTCACAAGCTACGATCCATAACCAGAGGCGGTGTGCGGATTGAGAA    1440
246       CATGGATTTGGGACTTCACAAGCTACGATCCATAACCAGAGGCGGTGTGCGGATTGAGAA    1440
353       CATGGATTTGGGACTTCACAAGCTACGATCCATAACCAGAGGCGGTGTGCGGATTGAGAA    1440
          ************************************************************

DAH       GAATCATAAGCTGTGCTATGATAGGACCATCGATTGGCTGGAAATTCTGGCGGAAAACGA    1500
29B       GAATCATAAGCTGTGCTATGATAGGACCATCGATTGGCTGGAAATTCTGGCGGAAAACGA    1500
e19       GAATCATAAGCTGTGCTATGATAGGACCATCGATTGGCTGGAAATTCTGGCGGAAAACGA    1500
74e19     GAATCATAAGCTGTGCTATGATAGGACCATCGATTGGCTGGAAATTCTGGCGGAAAACGA    1500
74        GAATCATAAGCTGTGCTATGATAGGACCATCGATTGGCTGGAAATTCTGGCGGAAAACGA    1500
211       GAATCATAAGCTGTGCTATGATAGGACCATCGATTGGCTGGAAATTCTGGCGGAAAACGA    1500
246       GAATCATAAGCTGTGCTATGATAGGACCATCGATTGGCTGGAAATTCTGGCGGAAAACGA    1500
353       GAATCATAAGCTGTGCTATGATAGGACCATCGATTGGCTGGAAATTCTGGCGGAAAACGA    1500
          ************************************************************

DAH       AACCCAACTGGTGGTGCTGACAGAGAACGGCAAGGAGAAGGAGTGCAGGCTTTCCAAGTG    1560
29B       AACCCAACTGGTGGTGCTGACAGAGAACGGCAAGGAGAAGGAGTGCAGGCTTTCCAAGTG    1560
e19       AACCCAACTGGTGGTGCTGACAGAGAACGGCAAGGAGAAGGAGTGCAGGCTTTCCAAGTG    1560
74e19     AACCCAACTGGTGGTGCTGACAGAGAACGGCAAGGAGAAGGAGTGCAGGCTTTCCAAGTG    1560
74        AACCCAACTGGTGGTGCTGACAGAGAACGGCAAGGAGAAGGAGTGCAGGCTTTCCAAGTG    1560
211       AACCCAACTGGTGGTGCTGACAGAGAACGGCAAGGAGAAGGAGTGCAGGCTTTCCAAGTG    1560
246       AACCCAACTGGTGGTGCTGACAGAGAACGGCAAGGAGAAGGAGTGCAGGCTTTCCAAGTG    1560
353       AACCCAACTGGTGGTGCTGACAGAGAACGGCAAGGAGAAGGAGTGCAGGCTTTCCAAGTG    1560
          ************************************************************

DAH       CCCGGGGGAGATCAGAATTGAGGAGGGGCACGATACCACGGCTATTGAGGGAGAGCTTAA    1620
29B       CCCGGGGGAGATCAGAATTGAGGAGGGGCACGATACCACGGCTATTGAGGGAGAGCTTAA    1620
e19       CCCGGGGGAGATCAGAATTGAGGAGGGGCACGATACCACGGCTATTGAGGGAGAGCTTAA    1620
74e19     CCCGGGGGAGATCAGAATTGAGGAGGGGCACGATACCACGGCTATTGAGGGAGAGCTTAA    1620
74        CCCGGGGGAGATCAGAATTGAGGAGGGGCACGATACCACGGCTATTGAGGGAGAGCTTAA    1620
211       CCCGGGGGAGATCAGAATTGAGGAGGGGCACGATACCACGGCTATTGAGGGAGAGCTTAA    1620
246       CCCGGGGGAGATCAGAATTGAGGAGGGGCACGATACCACGGCTATTGAGGGAGAGCTTAA    1620
353       CCCGGGGGAGATCAGAATTGAGGAGGGGCACGATACCACGGCTATTGAGGGAGAGCTTAA    1620
          ************************************************************
```

```
DAH       TGCCAGTTGTCAGCTGCACAATAATAGGCGCCTGTGCTGGAACAGCAAACTCTGCCAGAC    1680
29B       TGCCAGTTGTCAGCTGCACAATAATAGGCGCCTGTGCTGGAACAGCAAACTCTGCCAGAC    1680
e19       TGCCAGTTGTCAGCTGCACAATAATAGGCGCCTGTGCTGGAACAGCAAACTCTGCCAGAC    1680
74e19     TGCCAGTTGTCAGCTGCACAATAATAGGCGCCTGTGCTGGAACAGCAAACTCTGCCAGAC    1680
74        TGCCAGTTGTCAGCTGCACAATAATAGGCGCCTGTGCTGGAACAGCAAACTCTGCCAGAC    1680
211       TGCCAGTTGTCAGCTGCACAATAATAGGCGCCTGTGCTGGAACAGCAAACTCTGCCAGAC    1680
246       TGCCAGTTGTCAGCTGCACAATAATAGGCGCCTGTGCTGGAACAGCAAACTCTGCCAGAC    1680
353       TGCCAGTTGTCAGCTGCACAATAATAGGCGCCTGTGCTGGAACAGCAAACTCTGCCAGAC    1680
          ************************************************************

DAH       GAgtgagttggccggtgtaaagttataccgttttcttattaacatttttcgcgttttttt    1740
29B       GAgtgagttggccggtgtaaagttataccgttttcttattaacatttttcgcgttttttt    1740
e19       GAgtgagttggccggtgtaaagttataccgttttcttattaacatttttcgcgttttttt    1740
74e19     GAgtgagttggccggtgtaaagttataccgttttcttattaacatttttcgcgttttttt    1740
74        GAgtgagttggccggtgtaaagttataccgttttcttattaacatttttcgcgttttttt    1740
211       GAgtgagttggccggtgtaaagttataccgttttcttattaacatttttcgcgttttttt    1740
246       GAgtgagttggccggtgtaaagttataccgttttcttattaacatttttcgcgttttttt    1740
353       GAgtgagttggccggtgtaaagttataccgttttcttattaacatttttcgcgttttttt    1740
          ************************************************************

DAH       cttctccagAATGCCCTGAAAAGTGCAGAAATAACTGCATCGATGAGCACACCTGCTGCA    1800
29B       cttctccagAATGCCCTGAAAAGTGCAGAAATAACTGCATCGATGAGCACACCTGCTGCA    1800
e19       cttctccagAATGCCCTGAAAAGTGCAGAAATAACTGCATCGATGAGCACACCTGCTGCA    1800
74e19     cttctccagAATGCCCTGAAAAGTGCAGAAATAACTGCATCGATGAGCACACCTGCTGCA    1800
74        cttctccagAATGCCCTGAAAAGTGCAGAAATAACTGCATCGATGAGCACACCTGCTGCA    1800
211       cttctccagAATGCCCTGAAAAGTGCAGAAATAACTGCATCGATGAGCACACCTGCTGCA    1800
246       cttctccagAATGCCCTGAAAAGTGCAGAAATAACTGCATCGATGAGCACACCTGCTGCA    1800
353       cttctccagAATGCCCTGAAAAGTGCAGAAATAACTGCATCGATGAGCACACCTGCTGCA    1800
          ************************************************************

DAH       GCCAGGATTGTTTGGGTGGATGCGTGATCGATAAGAATGGGAATGAGAGCTGCATCTCCT    1860
29B       GCCAGGATTGTTTGGGTGGATGCGTGATCGATAAGAATGGGAATGAGAGCTGCATCTCCT    1860
e19       GCCAGGATTGTTTGGGTGGATGCGTGATCGATAAGAATGGGAATGAGAGCTGCATCTCCT    1860
74e19     GCCAGGATTGTTTGGGTGGATGCGTGATCGATAAGAATGGGAATGAGAGCTGCATCTCCT    1860
74        GCCAGGATTGTTTGGGTGGATGCGTGATCGATAAGAATGGGAATGAGAGCTGCATCTCCT    1860
211       GCCAGGATTGTTTGGGTGGATGCGTGATCGATAAGAATGGGAATGAGAGCTGCATCTCCT    1860
246       GCCAGGATTGTTTGGGTGGATGCGTGATCGATAAGAATGGGAATGAGAGCTGCATCTCCT    1860
353       GCCAGGATTGTTTGGGTGGATGCGTGATCGATAAGAATGGGAATGAGAGCTGCATCTCCT    1860
          ************************************************************

DAH       GTCGAAATGTGTCTTTCAACAACATCTGTATGGACTCCTGTCCGAAAGGCTATTATCAGg    1920
29B       GTCGAAATGTGTCTTTCAACAACATCTGTATGGACTCCTGTCCGAAAGGCTATTATCAGg    1920
e19       GTCGAAATGTGTCTTTCAACAACATCTGTATGGACTCCTGTCCGAAAGGCTATTATCAGg    1920
74e19     GTCGAAATGTGTCTTTCAACAACATCTGTATGGACTCCTGTCCGAAAGGCTATTATCAGg    1920
74        GTCGAAATGTGTCTTTCAACAACATCTGTATGGACTCCTGTCCGAAAGGCTATTATCAGg    1920
211       GTCGAAATGTGTCTTTCAACAACATCTGTATGGACTCCTGTCCGAAAGGCTATTATCAGg    1920
246       GTCGAAATGTGTCTTTCAACAACATCTGTATGGACTCCTGTCCGAAAGGCTATTATCAGg    1920
353       GTCGAAATGTGTCTTTCAACAACATCTGTATGGACTCCTGTCCGAAAGGCTATTATCAGg    1920
          ************************************************************

DAH       taactaatgattcacttttagtatacaacaagtagctaactttcaaactcttttgcagTT    1980
29B       taactaatgattcacttttagtatacaacaagtagctaactttcaaactcttttgcagTT    1980
e19       taactaatgattcacttttagtatacaacaagtagctaactttcaaactcttttgcagTT    1980
74e19     taactaatgattcacttttagtatacaacaagtagctaactttcaaactcttttgcagTT    1980
74        taactaatgattcacttttagtatacaacaagtagctaactttcaaactcttttgcagTT    1980
211       taactaatgattcacttttagtatacaacaagtagctaactttcaaactcttttgcagTT    1980
246       taactaatgattcacttttagtatacaacaagtagctaactttcaaactcttttgcagTT    1980
353       taactaatgattcacttttagtatacaacaagtagctaactttcaaactcttttgcagTT    1980
          ************************************************************
```

```
DAH      CGACAGCCGCTGCGTAACGGCGAACGAGTGCATCACACTGACAAAGTTTGAAACGAACAG   2040
29B      CGACAGCCGCTGCGTAACGGCGAACGAGTGCATCACACTGACAAAGTTTGAAACGAACAG   2040
e19      CGACAGCCGCTGCGTAACGGCGAACGAGTGCATCACACTGACAAAGTTTGAAACGAACAG   2040
74e19    CGACAGCCGCTGCGTAACGGCGAACGAGTGCATCACACTGACAAAGTTTGAAACGAACAG   2040
74       CGACAGCCGCTGCGTAACGGCGAACGAGTGCATCACACTGACAAAGTTTGAAACGAACAG   2040
211      CGACAGCCGCTGCGTAACGGCGAACGAGTGCATCACACTGACAAAGTTTGAAACGAACAG   2040
246      CGACAGCCGCTGCGTAACGGCGAACGAGTGCATCACACTGACAAAGTTTGAAACGAACAG   2040
353      CGACAGCCGCTGCGTAACGGCGAACGAGTGCATCACACTGACAAAGTTTGAAACGAACAG   2040
         ************************************************************

DAH      TGTGTATTCCGGTATTCCATACAACGGACAATGTATCACCCACTGTCCAACGGGGTACCA   2100
29B      TGTGTATTCCGGTATTCCATACAACGGACAATGTATCACCCACTGTCCAACGGGGTACCA   2100
e19      TGTGTATTCCGGTATTCCATACAACGGACAATGTATCACCCACTGTCCAACGGGGTACCA   2100
74e19    TGTGTATTCCGGTATTCCATACAACGGACAATGTATCACCCACTGTCCAACGGGGTACCA   2100
74       TGTGTATTCCGGTATTCCATACAACGGACAATGTATCACCCACTGTCCAACGGGGTACCA   2100
211      TGTGTATTCCGGTATTCCATACAACGGACAATGTATCACCCACTGTCCAACGGGGTACCA   2100
246      TGTGTATTCCGGTATTCCATACAACGGACAATGTATCACCCACTGTCCAACGGGGTACCA   2100
353      TGTGTATTCCGGTATTCCATACAACGGACAATGTATCACCCACTGTCCAACGGGGTACCA   2100
         ************************************************************

DAH      GAAGTCAGAGAACAAGCGCATGTGCGAACCTTGTCCGGGCGGCAAGTGTGACAAGGAGTG   2160
29B      GAAGTCAGAGAACAAGCGCATGTGCGAACCTTGTCCGGGCGGCAAGTGTGACAAGGAGTG   2160
e19      GAAGTCAGAGAACAAGCGCATGTGCGAACCTTGTCCGGGCGGCAAGTGTGACAAGGAGTG   2160
74e19    GAAGTCAGAGAACAAGCGCATGTGCGAACCTTGTCCGGGCGGCAAGTGTGACAAGGAGTG   2160
74       GAAGTCAGAGAACAAGCGCATGTGCGAACCTTGTCCGGGCGGCAAGTGTGACAAGGAGTG   2160
211      GAAGTCAGAGAACAAGCGCATGTGCGAACCTTGTCCGGGCGGCAAGTGTGACAAGGAGTG   2160
246      GAAGTCAGAGAACAAGCGCATGTGCGAACCTTGTCCGGGCGGCAAGTGTGACAAGGAGTG   2160
353      GAAGTCAGAGAACAAGCGCATGTGCGAACCTTGTCCGGGCGGCAAGTGTGACAAGGAGTG   2160
         ************************************************************

DAH      CTCCTCCGGTCTTATCGACAGTTTGGAGCGTGCTCGGGAGTTCCACGGCTGCACCATTAT   2220
29B      CTCCTCCGGTCTTATCGACAGTTTGGAGCGTGCTCGGGAGTTCCACGGCTGCACCATTAT   2220
e19      CTCCTCCGGTCTTATCGACAGTTTGGAGCGTGCTCGGGAGTTCCACGGCTGCACCATTAT   2220
74e19    CTCCTCCGGTCTTATCGACAGTTTGGAGCGTGCTCGGGAGTTCCACGGCTGCACCATTAT   2220
74       CTCCTCCGGTCTTATCGACAGTTTGGAGCGTGCTCGGGAGTTCCACGGCTGCACCATTAT   2220
211      CTCCTCCGGTCTTATCGACAGTTTGGAGCGTGCTCGGGAGTTCCACGGCTGCACCATTAT   2220
246      CTCCTCCGGTCTTATCGACAGTTTGGAGCGTGCTCGGGAGTTCCACGGCTGCACCATTAT   2220
353      CTCCTCCGGTCTTATCGACAGTTTGGAGCGTGCTCGGGAGTTCCACGGCTGCACCATTAT   2220
         ************************************************************

DAH      AACCGGAACCGAGCCCCTTACCATCAGCATTAAACGTGAAAGCGGCGgtaagtgtttttt   2280
29B      AACCGGAACCGAGCCCCTTACCATCAGCATTAAACGTGAAAGCGGCGgtaagtgtttttt   2280
e19      AACCGGAACCGAGCCCCTTACCATCAGCATTAAACGTGAAAGCGGCGgtaagtgtttttt   2280
74e19    AACCGGAACCGAGCCCCTTACCATCAGCATTAAACGTGAAAGCGGCGgtaagtgtttttt   2280
74       AACCGGAACCGAGCCCCTTACCATCAGCATTAAACGTGAAAGCGGCGgtaagtgtttttt   2280
211      AACCGGAACCGAGCCCCTTACCATCAGCATTAAACGTGAAAGCGGCGgtaagtgtttttt   2280
246      AACCGGAACCGAGCCCCTTACCATCAGCATTAAACGTGAAAGCGGCGgtaagtgtttttt   2280
353      AACCGGAACCGAGCCCCTTACCATCAGCATTAAACGTGAAAGCGGCGgtaagtgtttttt   2280
         ************************************************************

DAH      gctgctctaaattaaagtataatctctatattaaaacttcttgtttcttcagCTCACGTC   2340
29B      gctgctctaaattaaagtataatctctatattaaaacttcttgtttcttcagCTCACGTC   2340
e19      gctgctctaaattaaagtataatctctatattaaaacttcttgtttcttcagCTCACGTC   2340
74e19    gctgctctaaattaaagtataatctctatattaaaacttcttgtttcttcagCTCACGTC   2340
74       gctgctctaaattaaagtataatctctatattaaaacttcttgtttcttcagCTCACGTC   2340
211      gctgctctaaattaaagtataatctctatattaaaacttcttgtttcttcagCTCACGTC   2340
246      gctgctctaaattaaagtataatctctatattaaaacttcttgtttcttcagCTCACGTC   2340
353      gctgctctaaattaaagtataatctctatattaaaacttcttgtttcttcagCTCACGTC   2340
         ************************************************************
```

```
DAH     ATGGATGAATTAAAATATGGCCTGGCTGCCGTCCATAAAATTCAGTCGTCCCTAATGGTT  2400
29B     ATGGATGAATTAAAATATGGCCTGGCTGCCGTCCATAAAATTCAGTCGTCCCTAATGGTT  2400
e19     ATGGATGAATTAAAATATGGCCTGGCTGCCGTCCATAAAATTCAGTCGTCCCTAATGGTT  2400
74e19   ATGGATGAATTAAAATATGGCCTGGCTGCCGTCCATAAAATTCAGTCGTCCCTAATGGTT  2400
74      ATGGATGAATTAAAATATGGCCTGGCTGCCGTCCATAAAATTCAGTCGTCCCTAATGGTT  2400
211     ATGGATGAATTAAAATATGGCCTGGCTGCCGTCCATAAAATTCAGTCGTCCCTAATGGTT  2400
246     ATGGATGAATTAAAATATGGCCTGGCTGCCGTCCATAAAATTCAGTCGTCCCTAATGGTT  2400
353     ATGGATGAATTAAAATATGGCCTGGCTGCCGTCCATAAAATTCAGTCGTCCCTAATGGTT  2400
        ************************************************************

DAH     CATTTGACCTACGGATTGAAGTCCTTGAAATTCTTTCAATCCCTAACTGAAATTAGCGGC  2460
29B     CATTTGACCTACGGATTGAAGTCCTTGAAATTCTTTCAATCCCTAACTGAAATTAGCGGC  2460
e19     CATTTGACCTACGGATTGAAGTCCTTGAAATTCTTTCAATCCCTAACTGAAATTAGCGGC  2460
74e19   CATTTGACCTACGGATTGAAGTCCTTGAAATTCTTTCAATCCCTAACTGAAATTAGCGGC  2460
74      CATTTGACCTACGGATTGAAGTCCTTGAAATTCTTTCAATCCCTAACTGAAATTAGCGGC  2460
211     CATTTGACCTACGGATTGAAGTCCTTGAAATTCTTTCAATCCCTAACTGAAATTAGCGGC  2460
246     CATTTGACCTACGGATTGAAGTCCTTGAAATTCTTTCAATCCCTAACTGAAATTAGCGGC  2460
353     CATTTGACCTACGGATTGAAGTCCTTGAAATTCTTTCAATCCCTAACTGAAATTAGCGGC  2460
        ************************************************************

DAH     GATCCGCCGATGGACGCGGATAAATATGCTTTGTATGTGCTTGATAATCGCGATCTAGAT  2520
29B     GATCCGCCGATGGACGCGGATAAATATGCTTTGTATGTGCTTGATAATCGCGATCTAGAT  2520
e19     GATCCGCCGATGGACGCGGATAAATATGCTTTGTATGTGCTTGATAATCGCGATCTAGAT  2520
74e19   GATCCGCCGATGGACGCGGATAAATATGCTTTGTATGTGCTTGATAATCGCGATCTAGAT  2520
74      GATCCGCCGATGGACGCGGATAAATATGCTTTGTATGTGCTTGATAATCGCGATCTAGAT  2520
211     GATCCGCCGATGGACGCGGATAAATATGCTTTGTATGTGCTTGATAATCGCGATCTAGAT  2520
246     GATCCGCCGATGGACGCGGATAAATATGCTTTGTATGTGCTTGATAATCGCGATCTAGAT  2520
353     GATCCGCCGATGGACGCGGATAAATATGCTTTGTATGTGCTTGATAATCGCGATCTAGAT  2520
        ************************************************************

DAH     GAGCTCTGGGGACCCAACCAAACGGTGTTCATTAGGAAGGGCGGCGTCTTCTTTCATTTC  2580
29B     GAGCTCTGGGGACCCAACCAAACGGTGTTCATTAGGAAGGGCGGCGTCTTCTTTCATTTC  2580
e19     GAGCTCTGGGGACCCAACCAAACGGTGTTCATTAGGAAGGGCGGCGTCTTCTTTCATTTC  2580
74e19   GAGCTCTGGGGACCCAACCAAACGGTGTTCATTAGGAAGGGCGGCGTCTTCTTTCATTTC  2580
74      GAGCTCTGGGGACCCAACCAAACGGTGTTCATTAGGAAGGGCGGCGTCTTCTTTCATTTC  2580
211     GAGCTCTGGGGACCCAACCAAACGGTGTTCATTAGGAAGGGCGGCGTCTTCTTTCATTTC  2580
246     GAGCTCTGGGGACCCAACCAAACGGTGTTCATTAGGAAGGGCGGCGTCTTCTTTCATTTC  2580
353     GAGCTCTGGGGACCCAACCAAACGGTGTTCATTAGGAAGGGCGGCGTCTTCTTTCATTTC  2580
        ************************************************************

DAH     AACCCAAAACTATGTGTGTCCACCATTAACCAGTTGCTGCCCATGCTGGCCTCCAAGCCA  2640
29B     AACCCAAAACTATGTGTGTCCACCATTAACCAGTTGCTGCCCATGCTGGCCTCCAAGCCA  2640
e19     AACCCAAAACTATGTGTGTCCACCATTAACCAGTTGCTGCCCATGCTGGCCTCCAAGCCA  2640
74e19   AACCCAAAACTATGTGTGTCCACCATTAACCAGTTGCTGCCCATGCTGGCCTCCAAGCCA  2640
74      AACCCAAAACTATGTGTGTCCACCATTAACCAGTTGCTGCCCATGCTGGCCTCCAAGCCA  2640
211     AACCCAAAACTATGTGTGTCCACCATTAACCAGTTGCTGCCCATGCTGGCCTCCAAGCCA  2640
246     AACCCAAAACTATGTGTGTCCACCATTAACCAGTTGCTGCCCATGCTGGCCTCCAAGCCA  2640
353     AACCCAAAACTATGTGTGTCCACCATTAACCAGTTGCTGCCCATGCTGGCCTCCAAGCCA  2640
        ************************************************************

DAH     AAGTTTTTTGAAAAGTCAGATGTGGGCGCAGACTCGAATGGAAACCGCGGATCATgtaag  2700
29B     AAGTTTTTTGAAAAGTCAGATGTGGGCGCAGACTCGAATGGAAACCGCGGATCATgtaag  2700
e19     AAGTTTTTTGAAAAGTCAGATGTGGGCGCAGACTCGAATGGAAACCGCGGATCATgtaag  2700
74e19   AAGTTTTTTGAAAAGTCAGATGTGGGCGCAGACTCGAATGGAAACCGCGGATCATgtaag  2700
74      AAGTTTTTTGAAAAGTCAGATGTGGGCGCAGACTCGAATGGAAACCGCGGATCATgtaag  2700
211     AAGTTTTTTGAAAAGTCAGATGTGGGCGCAGACTCGAATGGAAACCGCGGATCATgtaag  2700
246     AAGTTTTTTGAAAAGTCAGATGTGGGCGCAGACTCGAATGGAAACCGCGGATCATgtaag  2700
353     AAGTTTTTTGAAAAGTCAGATGTGGGCGCAGACTCGAATGGAAACCGCGGATCATgtaag  2700
        ************************************************************
```

```
DAH      taatctttaacgaagattttagtaaaattttaaactttgtcagtactaacctaaatgaaa   2760
29B      taatctttaacgaagattttagtaaaattttaaactttgtcagtactaacctaaatgaaa   2760
e19      taatctttaacgaagattttagtaaaattttaaactttgtcagtactaacctaaatgaaa   2760
74e19    taatctttaacgaagattttagtaaaattttaaactttgtcagtactaacctaaatgaaa   2760
74       taatctttaacgaagattttagtaaaattttaaactttgtcagtactaacctaaatgaaa   2760
211      taatctttaacgaagattttagtaaaattttaaactttgtcagtactaacctaaatgaaa   2760
246      taatctttaacgaagattttagtaaaattttaaactttgtcagtactaacctaaatgaaa   2760
353      taatctttaacgaagattttagtaaaattttaaactttgtcagtactaacctaaatgaaa   2760
         ************************************************************

DAH      catttccagGTGGAACAGCCGTTCTCAATGTCACATTACAATCAGTGGGAGCAAACTCCG   2820
29B      catttccagGTGGAACAGCCGTTCTCAATGTCACATTACAATCAGTGGGAGCAAACTCCG   2820
e19      catttccagGTGGAACAGCCGTTCTCAATGACACATTACAATCAGTGGGAGCAAACTCCG   2820
74e19    catttccagGTGGAACAGCCGTTCTCAATGACACATTACAATCAGTGGGAGCAAACTCCG   2820
74       catttccagGTGGAACAGCCGTTCTCAATGTCACATTACAATCAGTGGGAGCAAACTCCG   2820
211      catttccagGTGGAACAGCCGTTCTCAATGTCACATTACAATCAGTGGGAGCAAACTCCG   2820
246      catttccagGTGGAACAGCCGTTCTCAATGTCACATTACAATCAGTGGGAGCAAACTCCG   2820
353      catttccagGTGGAACAGCCGTTCTCAATGTCACATTACAATCAGTGGGAGCAAACTCCG   2820
         ****************************** *****************************

DAH      CTATGCTGAACGTCACGACAAAAGTTGAAATAGGAGAGCCCCAAAAGCCGAGCAATGCTA   2880
29B      CTATGCTGAACGTCACGACAAAAGTTGAAATAGGAGAGCCCCAAAAGCCGAGCAATGCTA   2880
e19      CTATGCTGAACGTCACGACAAAAGTTGAAATAGGAGAGCCCCAAAAGCCGAGCAATGCTA   2880
74e19    CTATGCTGAACGTCACGACAAAAGTTGAAATAGGAGAGCCCCAAAAGCCGAGCAATGCTA   2880
74       CTATGCTGAACGTCACGACAAAAGTTGAAATAGGAGAGCCCCAAAAGCCGAGCAATGCTA   2880
211      CTATGCTGAACGTCACGACAAAAGTTGAAATAGGAGAGCCCCAAAAGCCGAGCAATGCTA   2880
246      CTATGCTGAACGTCACGACAAAAGTTGAAATAGGAGAGCCCCAAAAGCCGAGCAATGCTA   2880
353      CTATGCTGAACGTCACGACAAAAGTTGAAATAGGAGAGCCCCAAAAGCCGAGCAATGCTA   2880
         ************************************************************

DAH      CAATTGTTTTTAAGGATCCGCGCGCCTTCATCGGTTTCGTGTTTTATCATATGATCGATC   2940
29B      CAATTGTTTTTAAGGATCCGCGCGCCTTCATCGGTTTCGTGTTTTATCATATGATCGATC   2940
e19      CAATTGTTTTTAAGGATCCGCGCGCCTTCATCGGTTTCGTGTTTTATCATATGATCGATC   2940
74e19    CAATTGTTTTTAAGGATCCGCGCGCCTTCATCGGTTTCGTGTTTTATCATATGATCGATC   2940
74       CAATTGTTTTTAAGGATCCGCGCGCCTTCATCGGTTTCGTGTTTTATCATATGATCGATC   2940
211      CAATTGTTTTTAAGGATCCGCGCGCCTTCATCGGTTTCGTGTTTTATCATATGATCGATC   2940
246      CAATTGTTTTTAAGGATCCGCGCGCCTTCATCGGTTTCGTGTTTTATCATATGATCGATC   2940
353      CAATTGTTTTTAAGGATCCGCGCGCCTTCATCGGTTTCGTGTTTTATCATATGATCGATC   2940
         ************************************************************

DAH      CGTACGGGAACTCAACTAAAAGCAGTGACGATCCATGCGATGATCGCTGGAAGGTTAGCT   3000
29B      CGTACGGGAACTCAACTAAAAGCAGTGACGATCCATGCGATGATCGCTGGAAGGTTAGCT   3000
e19      CGTACGGGAACTCAACTAAAAGCAGTGACGATCCATGCGATGATCGCTGGAAGGTTAGCT   3000
74e19    CGTACGGGAACTCAACTAAAAGCAGTGACGATCCATGCGATGATCGCTGGAAGGTTAGCT   3000
74       CGTACGGGAACTCAACTAAAAGCAGTGACGATCCATGCGATGATCGCTGGAAGGTTAGCT   3000
211      CGTACGGGAACTCAACTAAAAGCAGTGACGATCCATGCGATGATCGCTGGAAGGTTAGCT   3000
246      CGTACGGGAACTCAACTAAAAGCAGTGACGATCCATGCGATGATCGCTGGAAGGTTAGCT   3000
353      CGTACGGGAACTCAACTAAAAGCAGTGACGATCCATGCGATGATCGCTGGAAGGTTAGCT   3000
         ************************************************************

DAH      CTCCGGAAAAGAGCGGGGTCATGGTATTAAGCAATTTGATTCCGTACACTAACTACTCCT   3060
29B      CTCCGGAAAAGAGCGGGGTCATGGTATTAAGCAATTTGATTCCGTACACTAACTACTCCT   3060
e19      CTCCGGAAAAGAGCGGGGTCATGGTATTAAGCAATTTGATTCCGTACACTAACTACTCCT   3060
74e19    CTCCGGAAAAGAGCGGGGTCATGGTATTAAGCAATTTGATTCCGTACACTAACTACTCCT   3060
74       CTCCGGAAAAGAGCGGGGTCATGGTATTAAGCAATTTGATTCCGTACACTAACTACTCCT   3060
211      CTCCGGAAAAGAGCGGGGTCATGGTATTAAGCAATTTGATTCCGTACACTAACTACTCCT   3060
246      CTCCGGAAAAGAGCGGGGTCATGGTATTAAGCAATTTGATTCCGTACACTAACTACTCCT   3060
353      CTCCGGAAAAGAGCGGGGTCATGGTATTAAGCAATTTGATTCCGTACACTAACTACTCCT   3060
         ************************************************************
```

```
DAH      ACTACGTTCGGACCATGGCTATATCCTCGGAATTGACAAACGCGGAGAGCGACGTGAAGA   3120
29B      ACTACGTTCGGACCATGGCTATATCCTCGGAATTGACAAACGCGGAGAGCGACGTGAAGA   3120
e19      ACTACGTTCGGACCATGGCTATATCCTCGGAATTGACAAACGCGGAGAGCGACGTGAAGA   3120
74e19    ACTACGTTCGGACCATGGCTATATCCTCGGAATTGACAAACGCGGAGAGCGACGTGAAGA   3120
74       ACTACGTTCGGACCATGGCTATATCCTCGGAATTGACAAACGCGGAGAGCGACGTGAAGA   3120
211      ACTACGTTCGGACCATGGCTATATCCTCGGAATTGACAAACGCGGAGAGCGACGTGAAGA   3120
246      ACTACGTTCGGACCATGGCTATATCCTCGGAATTGACAAACGCGGAGAGCGACGTGAAGA   3120
353      ACTACGTTCGGACCATGGCTATATCCTCGGAATTGACAAACGCGGAGAGCGACGTGAAGA   3120
         ************************************************************

DAH      ACTTTAGGACGAATCCCGGACGACCGTCAAAGGTTACGGAGGTGGTAGCAACCGCCATTT   3180
29B      ACTTTAGGACGAATCCCGGACGACCGTCAAAGGTTACGGAGGTGGTAGCAACCGCCATTT   3180
e19      ACTTTAGGACGAATCCCGGACGACCGTCAAAGGTTACGGAGGTGGTAGCAACCGCCATTT   3180
74e19    ACTTTAGGACGAATCCCGGACGACCGTCAAAGGTTACGGAGGTGGTAGCAACCGCCATTT   3180
74       ACTTTAGGACGAATCCCGGACGACCGTCAAAGGTTACGGAGGTGGTAGCAACCGCCATTT   3180
211      ACTTTAGGACGAATCCCGGACGACCGTCAAAGGTTACGGAGGTGGTAGCAACCGCCATTT   3180
246      ACTTTAGGACGAATCCCGGACGACCGTCAAAGGTTACGGAGGTGGTAGCAACCGCCATTT   3180
353      ACTTTAGGACGAATCCCGGACGACCGTCAAAGGTTACGGAGGTGGTAGCAACCGCCATTT   3180
         ************************************************************

DAH      CAGATTCGAAAATTgtgagtatgaaattgtgcaatagattagtttaggattataaatgat   3240
29B      CAGATTCGAAAATTgtgagtatgaaattgtgcaatagattagtttaggattataaatgat   3240
e19      CAGATTCGAAAATTgtgagtatgaaattgtgcaatagattagtttaggattataaatgat   3240
74e19    CAGATTCGAAAATTgtgagtatgaaattgtgcaatagattagtttaggattataaatgat   3240
74       CAGATTCGAAAATTgtgagtatgaaattgtgcaatagattagtttaggattataaatgat   3240
211      CAGATTCGAAAATTgtgagtatgaaattgtgcaatagattagtttaggattataaatgat   3240
246      CAGATTCGAAAATTgtgagtatgaaattgtgcaatagattagtttaggattataaatgat   3240
353      CAGATTCGAAAATTgtgagtatgaaattgtgcaatagattagtttaggattataaatgat   3240
         ************************************************************

DAH      tactaaatgtcaagtaattgttcactttgactgatcaacatttattgccttttcagAACG   3300
29B      tactaaatgtcaagtaattgttcactttgactgatcaacatttattgccttttcagAACG   3300
e19      tactaaatgtcaagtaattgttcactttgactgatcaacatttattgccttttcagAACG   3300
74e19    tactaaatgtcaagtaattgttcactttgactgatcaacatttattgccttttcagAACG   3300
74       tactaaatgtcaagtaattgttcactttgactgatcaacatttattgccttttcagAACG   3300
211      tactaaatgtcaagtaattgttcactttgactgatcaacatttattgccttttcagAACG   3300
246      tactaaatgtcaagtaattgttcactttgactgatcaacatttattgccttttcagAACG   3300
353      tactaaatgtcaagtaattgttcactttgactgatcaacatttattgccttttcagAACG   3300
         ************************************************************

DAH      TAACATGGAGCTACCTAGATAAGCCTTATGGCGTGCTAACGCGCTATTTTATAAAAGCCA   3360
29B      TAACATGGAGCTACCTAGATAAGCCTTATGGCGTGCTAACGCGCTATTTTATAAAAGCCA   3360
e19      TAACATGGAGCTACCTAGATAAGCCTTATGGCGTGCTAACGCGCTATTTTATAAAAGCCA   3360
74e19    TAACATGGAGCTACCTAGATAAGCCTTATGGCGTGCTAACGCGCTATTTTATAAAAGCCA   3360
74       TAACATGGAGCTACCTAGATAAGCCTTATGGCGTGCTAACGCGCTATTTTATAAAAGCCA   3360
211      TAACATGGAGCTACCTAGATAAGCCTTATGGCGTGCTAACGCGCTATTTTATAAAAGCCA   3360
246      TAACATGGAGCTACCTAGATAAGCCTTATGGCGTGCTAACGCGCTATTTTATAAAAGCCA   3360
353      TAACATGGAGCTACCTAGATAAGCCTTATGGCGTGCTAACGCGCTATTTTATAAAAGCCA   3360
         ************************************************************

DAH      AACTTATAAATCGGCCTACTCGAAACAATAACCGGGATTACTGTACTGAACgtaagcaac   3420
29B      AACTTATAAATCGGCCTACTCGAAACAATAACCGGGATTACTGTACTGAACgtaagcaac   3420
e19      AACTTATAAATCGGCCTACTCGAAACAATAACCGGGATTACTGTACTGAACgtaagcaac   3420
74e19    AACTTATAAATCGGCCTACTCGAAACAATAACCGGGATTACTGTACTGAACgtaagcaac   3420
74       AACTTATAAATCGGCCTACTCGAAACAATAACCGGGATTACTGTACTGAACgtaagcaac   3420
211      AACTTATAAATCGGCCTACTCGAAACAATAACCGGGATTACTGTACTGAACgtaagcaac   3420
246      AACTTATAAATCGGCCTACTCGAAACAATAACCGGGATTACTGTACTGAACgtaagcaac   3420
353      AACTTATAAATCGGCCTACTCGAAACAATAACCGGGATTACTGTACTGAACgtaagcaac   3420
         ************************************************************
```

```
DAH       aacatagccaaattgtatccataattaattaatttaaatatttcatctgtgtagCTCTCG      3480
29B       aacatagccaaattgtatccataattaattaatttaaatatttcatctgtgtagCTCTCG      3480
e19       aacatagccaaattgtatccataattaattaatttaaatatttcatctgtgtagCTCTCG      3480
74e19     aacatagccaaattgtatccataattaattaatttaaatatttcatctgtgtagCTCTCG      3480
74        aacatagccaaattgtatccataattaattaatttaaatatttcatctgtgtagCTCTCG      3480
211       aacatagccaaattgtatccataattaattaatttaaatatttcatctgtgtagCTCTCG      3480
246       aacatagccaaattgtatccataattaattaatttaaatatttcatctgtgtagCTCTCG      3480
353       aacatagccaaattgtatccataattaattaatttaaatatttcatctgtgtagCTCTCG      3480
          ************************************************************

DAH       TCAAGGCCATGGAAAATGACCTGCCAGCCACAACGCCTACCAAGAAAATATCAGATCCTT      3540
29B       TCAAGGCCATGGAAAATGACCTGCCAGCCACAACGCCTACCAAGAAAATATCAGATCCTT      3540
e19       TCAAGGCCATGGAAAATGACCTGCCAGCCACAACGCCTACCAAGAAAATATCAGATCCTT      3540
74e19     TCAAGGCCATGGAAAATGACCTGCCAGCCACAACGCCTACCAAGAAAATATCAGATCCTT      3540
74        TCAAGGCCATGGAAAATGACCTGCCAGCCACAACGCCTACCAAGAAAATATCAGATCCTT      3540
211       TCAAGGCCATGGAAAATGACCTGCCAGCCACAACGCCTACCAAGAAAATATCAGATCCTT      3540
246       TCAAGGCCATGGAAAATGACCTGCCAGCCACAACGCCTACCAAGAAAATATCAGATCCTT      3540
353       TCAAGGCCATGGAAAATGACCTGCCAGCCACAACGCCTACCAAGAAAATATCAGATCCTT      3540
          ************************************************************

DAH       TAGCAGGCGACTGTAAGTGCGTGGAGGGTTCGAAGAAGACTAGCAGTCAGGAATACGATG      3600
29B       TAGCAGGCGACTGTAAGTGCGTGGAGGGTTCGAAGAAGACTAGCAGTCAGGAATACGATG      3600
e19       TAGCAGGCGACTGTAAGTGCGTGGAGGGTTCGAAGAAGACTAGCAGTCAGGAATACGATG      3600
74e19     TAGCAGGCGACTGTAAGTGCGTGGAGGGTTCGAAGAAGACTAGCAGTCAGGAATACGATG      3600
74        TAGCAGGCGACTGTAAGTGCGTGGAGGGTTCGAAGAAGACTAGCAGTCAGGAATACGATG      3600
211       TAGCAGGCGACTGTAAGTGCGTGGAGGGTTCGAAGAAGACTAGCAGTCAGGAATACGATG      3600
246       TAGCAGGCGACTGTAAGTGCGTGGAGGGTTCGAAGAAGACTAGCAGTCAGGAATACGATG      3600
353       TAGCAGGCGACTGTAAGTGCGTGGAGGGTTCGAAGAAGACTAGCAGTCAGGAATACGATG      3600
          ************************************************************

DAH       ATCGTAAAGTTCAAGCGGGCATGGAGTTTGAGAACGCGTTGCAAAACTTTATATTTGTTC      3660
29B       ATCGTAAAGTTCAAGCGGGCATGGAGTTTGAGAACGCGTTGCAAAACTTTATATTTGTTC      3660
e19       ATCGTAAAGTTCAAGCGGGCATGGAGTTTGAGAACGCGTTGCAAAACTTTATATTTGTTC      3660
74e19     ATCGTAAAGTTCAAGCGGGCATGGAGTTTGAGAACGCGTTGCAAAACTTTATATTTGTTC      3660
74        ATCGTAAAGTTCAAGCGGGCATGGAGTTTGAGAACGCGTTGCAAAACTTTATATTTGTTC      3660
211       ATCGTAAAGTTCAAGCGGGCATGGAGTTTGAGAACGCGTTGCAAAACTTTATATTTGTTC      3660
246       ATCGTAAAGTTCAAGCGGGCATGGAGTTTGAGAACGCGTTGCAAAACTTTATATTTGTTC      3660
353       ATCGTAAAGTTCAAGCGGGCATGGAGTTTGAGAACGCGTTGCAAAACTTTATATTTGTTC      3660
          ************************************************************

DAH       CAAACATTCGGAAAAGCAAGAATGGATCGTCTGACAAATCAGACGGAGCGGAAGGTGCAG      3720
29B       CAAACATTCGGAAAAGCAAGAATGGATCGTCTGACAAATCAGACGGAGCGGAAGGTGCAG      3720
e19       CAAACATTCGGAAAAGCAAGAATGGATCGTCTGACAAATCAGACGGAGCGGAAGGTGCAG      3720
74e19     CAAACATTCGGAAAAGCAAGAATGGATCGTCTGACAAATCAGACGGAGCGGAAGGTGCAG      3720
74        CAAACATTCGGAAAAGCAAGAATGGATCGTCTGACAAATCAGACGGAGCGGAAGGTGCAG      3720
211       CAAACATTCGGAAAAGCAAGAATGGATCGTCTGACAAATCAGACGGAGCGGAAGGTGCAG      3720
246       CAAACATTCGGAAAAGCAAGAATGGATCGTCTGACAAATCAGACGGAGCGGAAGGTGCAG      3720
353       CAAACATTCGGAAAAGCAAGAATGGATCGTCTGACAAATCAGACGGAGCGGAAGGTGCAG      3720
          ************************************************************

DAH       CTCTCGATTCTAATGCTATTCCAAATGGAGGAGCTACTAACCCTTCACGTAGAAGGAGAG      3780
29B       CTCTCGATTCTAATGCTATTCCAAATGGAGGAGCTACTAACCCTTCACGTAGAAGGAGAG      3780
e19       CTCTCGATTCTAATGCTATTCCAAATGGAGGAGCTACTAACCCTTCACGTAGAAGGAGAG      3780
74e19     CTCTCGATTCTAATGCTATTCCAAATGGAGGAGCTACTAACCCTTCACGTAGAAGGAGAG      3780
74        CTCTCGATTCTAATGCTATTCCAAATGGAGGAGCTACTAACCCTTCACGTAGAAGGAGAG      3780
211       CTCTCGATTCTAATGCTATTCCAAATGGAGGAGCTACTAACCCTTCACGTAGAAGGAGAG      3780
246       CTCTCGATTCTAATGCTATTCCAAATGGAGGAGCTACTAACCCTTCACGTAGAAGGAGAG      3780
353       CTCTCGATTCTAATGCTATTCCAAATGGAGGAGCTACTAACCCTTCACGTAGAAGGAGAG      3780
          ************************************************************
```

60

```
DAH       ACGTTGCGCTCGAGCCAGAGCTCGACGATGTAGAGGGCAGTGTACTTCTACGCCATGTGC     3840
29B       ACGTTGCGCTCGAGCCAGAGCTCGACGATGTAGAGGGCAGTGTACTTCTACGCCATGTGC     3840
e19       ACGTTGCGCTCGAGCCAGAGCTCGACGATGTAGAGGGCAGTGTACTTCTACGCCATGTGC     3840
74e19     ACGTTGCGCTCGAGCCAGAGCTCGACGATGTAGAGGGCAGTGTACTTCTACGCCATGTGC     3840
74        ACGTTGCGCTCGAGCCAGAGCTCGACGATGTAGAGGGCAGTGTACTTCTACGCCATGTGC     3840
211       ACGTTGCGCTCGAGCCAGAGCTCGACGATGTAGAGGGCAGTGTACTTCTACGCCATGTGC     3840
246       ACGTTGCGCTCGAGCCAGAGCTCGACGATGTAGAGGGCAGTGTACTTCTACGCCATGTGC     3840
353       ACGTTGCGCTCGAGCCAGAGCTCGACGATGTAGAGGGCAGTGTACTTCTACGCCATGTGC     3840
          ************************************************************

DAH       GCTCCATCACAGACGATACCGATGCATTTTTCGAAAAGGACGACGAAAATACCTATAAAG     3900
29B       GCTCCATCACAGACGATACCGATGCATTTTTCGAAAAGGACGACGAAAATACCTATAAAG     3900
e19       GCTCCATCACAGACGATACCGATGCATTTTTCGAAAAGGACGACGAAAATACCTATAAAG     3900
74e19     GCTCCATCACAGACGATACCGATGCATTTTTCGAAAAGGACGACGAAAATACCTATAAAG     3900
74        GCTCCATCACAGACGATACCGATGCATTTTTCGAAAAGGACGACGAAAATACCTATAAAG     3900
211       GCTCCATCACAGACGATACCGATGCATTTTTCGAAAAGGACGACGAAAATACCTATAAAG     3900
246       GCTCCATCACAGACGATACCGATGCATTTTTCGAAAAGGACGACGAAAATACCTATAAAG     3900
353       GCTCCATCACAGACGATACCGATGCATTTTTCGAAAAGGACGACGAAAATACCTATAAAG     3900
          ************************************************************

DAH       ACGAAGAAGACTTGTCCTCCAACAAACAATTCTATGAGGTGTTTGCCAAGGAATTGCCAC     3960
29B       ACGAAGAAGACTTGTCCTCCAACAAACAATTCTATGAGGTGTTTGCCAAGGAATTGCCAC     3960
e19       ACGAAGAAGACTTGTCCTCCAACAAACAATTCTATGAGGTGTTTGCCAAGGAATTGCCAC     3960
74e19     ACGAAGAAGACTTGTCCTCCAACAAACAATTCTATGAGGTGTTTGCCAAGGAATTGCCAC     3960
74        ACGAAGAAGACTTGTCCTCCAACAAACAATTCTATGAGGTGTTTGCCAAGGAATTGCCAC     3960
211       ACGAAGAAGACTTGTCCTCCAACAAACAATTCTATGAGGTGTTTGCCAAGGAATTGCCAC     3960
246       ACGAAGAAGACTTGTCCTCCAACAAACAATTCTATGAGGTGTTTGCCAAGGAATTGCCAC     3960
353       ACGAAGAAGACTTGTCCTCCAACAAACAATTCTATGAGGTGTTTGCCAAGGAATTGCCAC     3960
          ************************************************************

DAH       CAAATCAAACACATTTTGTCTTTGAAAAACTGCGCCACTTCACCCGCTACGCTATCTTCG     4020
29B       CAAATCAAACACATTTTGTCTTTGAAAAACTGCGCCACTTCACCCGCTACGCTATCTTCG     4020
e19       CAAATCAAACACATTTTGTCTTTGAAAAACTGCGCCACTTCACCCGCTACGCTATCTTCG     4020
74e19     CAAATCAAACACATTTTGTCTTTGAAAAACTGCGCCACTTCACCCGCTACGCTATCTTCG     4020
74        CAAATCAAACACATTTTGTCTTTGAAAAACTGCGCCACTTCACCCGCTACGCTATCTTCG     4020
211       CAAATCAAACACATTTTGTCTTTGAAAAACTGCGCCACTTCACCCGCTACGCTATCTTCG     4020
246       CAAATCAAACACATTTTGTCTTTGAAAAACTGCGCCACTTCACCCGCTACGCTATCTTCG     4020
353       CAAATCAAACACATTTTGTCTTTGAAAAACTGCGCCACTTCACCCGCTACGCTATCTTCG     4020
          ************************************************************

DAH       TGGTAGCCTGTAGAGAAGAAATCCCCAGCGAAAAATTAAGGGACACCAGTTTTAAGAAGT     4080
29B       TGGTAGCCTGTAGAGAAGAAATCCCCAGCGAAAAATTAAGGGACACCAGTTTTAAGAAGT     4080
e19       TGGTAGCCTGTAGAGAAGAAATCCCCAGCGAAAAATTAAGGGACACCAGTTTTAAGAAGT     4080
74e19     TGGTAGCCTGTAGAGAAGAAATCCCCAGCGAAAAATTAAGGGACACCAGTTTTAAGAAGT     4080
74        TGGTAGCCTGTAGAGAAGAAATCCCCAGCGAAAAATTAAGGGACACCAGTTTTAAGAAGT     4080
211       TGGTAGCCTGTAGAGAAGAAATCCCCAGCGAAAAATTAAGGGACACCAGTTTTAAGAAGT     4080
246       TGGTAGCCTGTAGAGAAGAAATCCCCAGCGAAAAATTAAGGGACACCAGTTTTAAGAAGT     4080
353       TGGTAGCCTGTAGAGAAGAAATCCCCAGCGAAAAATTAAGGGACACCAGTTTTAAGAAGT     4080
          ************************************************************

DAH       CGCTCTGCAGCGATTATGACACCGTTTTCCAAACTACAAAGAGAAAGAgtaggtggactt     4140
29B       CGCTCTGCAGCGATTATGACACCGTTTTCCAAACTACAAAGAGAAAGAgtaggtggactt     4140
e19       CGCTCTGCAGCGATTATGACACCGTTTTCCAAACTACAAAGAGAAAGAgtaggtggactt     4140
74e19     CGCTCTGCAGCGATTATGACACCGTTTTCCAAACTACAAAGAGAAAGAgtaggtggactt     4140
74        CGCTCTGCAGCGATTATGACACCGTTTTCCAAACTACAAAGAGAAAGAgtaggtggactt     4140
211       CGCTCTGCAGCGATTATGACACCGTTTTCCAAACTACAAAGAGAAAGAgtaggtggactt     4140
246       CGCTCTGCAGCGATTATGACACCGTTTTCCAAACTACAAAGAGAAAGAgtaggtggactt     4140
353       CGCTCTGCAGCGATTATGACACCGTTTTCCAAACTACAAAGAGAAAGAgtaggtggactt     4140
          ************************************************************
```

```
DAH      gagagcgtgtttactttcattactaacgatttggttttacttatagAATTTGCCGACATA   4200
29B      gagagcgtgtttactttcattactaacgatttggttttacttatagAATTTGCCGACATA   4200
e19      gagagcgtgtttactttcattactaacgatttggttttacttatagAATTTGCCGACATA   4200
74e19    gagagcgtgtttactttcattactaacgatttggttttacttatagAATTTGCCGACATA   4200
74       gagagcgtgtttactttcattactaacgatttggttttacttatagAATTTGCCGACATA   4200
211      gagagcgtgtttactttcattactaacgatttggttttacttatagAATTTGCCGACATA   4200
246      gagagcgtgtttactttcattactaacgatttggttttacttatagAATTTGCCGACATA   4200
353      gagagcgtgtttactttcattactaacgatttggttttacttatagAATTTGCCGACATA   4200
         ************************************************************

DAH      GTCATGGACCTAAAAGTAGATTTAGAACACGCCAACAACACCGAGTCCCCAGTACGGGTT   4260
29B      GTCATGGACCTAAAAGTAGATTTAGAACACGCCAACAACACCGAGTCCCCAGTACGGGTT   4260
e19      GTCATGGACCTAAAAGTAGATTTAGAACACGCCAACAACACCGAGTCCCCAGTACGGGTT   4260
74e19    GTCATGGACCTAAAAGTAGATTTAGAACACGCCAACAACACCGAGTCCCCAGTACGGGTT   4260
74       GTCATGGACCTAAAAGTAGATTTAGAACACGCCAACAACACCGAGTCCCCAGTACGGGTT   4260
211      GTCATGGACCTAAAAGTAGATTTAGAACACGCCAACAACACCGAGTCCCCAGTACGGGTT   4260
246      GTCATGGACCTAAAAGTAGATTTAGAACACGCCAACAACACCGAGTCCCCAGTACGGGTT   4260
353      GTCATGGACCTAAAAGTAGATTTAGAACACGCCAACAACACCGAGTCCCCAGTACGGGTT   4260
         ************************************************************

DAH      CGCTGGACGCCACCAGTAGATCCCAACGGAGAAATTGTCACCTATGAAGTGGCCTACAAG   4320
29B      CGCTGGACGCCACCAGTAGATCCCAACGGAGAAATTGTCACCTATGAAGTGGCCTACAAG   4320
e19      CGCTGGACGCCACCAGTAGATCCCAACGGAGAAATTGTCACCTATGAAGTGGCCTACAAG   4320
74e19    CGCTGGACGCCACCAGTAGATCCCAACGGAGAAATTGTCACCTATGAAGTGGCCTACAAG   4320
74       CGCTGGACGCCACCAGTAGATCCCAACGGAGAAATTGTCACCTATGAAGTGGCCTACAAG   4320
211      CGCTGGACGCCACCAGTAGATCCCAACGGAGAAATTGTCACCTATGAAGTGGCCTACAAG   4320
246      CGCTGGACGCCACCAGTAGATCCCAACGGAGAAATTGTCACCTATGAAGTGGCCTACAAG   4320
353      CGCTGGACGCCACCAGTAGATCCCAACGGAGAAATTGTCACCTATGAAGTGGCCTACAAG   4320
         ************************************************************

DAH      TTGCAAAAACCCGATCAAGTGGAAGAAAAGAAGTGCATTCCGGCTGCTGACTTCAACCAG   4380
29B      TTGCAAAAACCCGATCAAGTGGAAGAAAAGAAGTGCATTCCGGCTGCTGACTTCAACCAG   4380
e19      TTGCAAAAACCCGATCAAGTGGAAGAAAAGAAGTGCATTCCGGCTGCTGACTTCAACCAG   4380
74e19    TTGCAAAAACCCGATCAAGTGGAAGAAAAGAAGTGCATTCCGGCTGCTGACTTCAACCAG   4380
74       TTGCAAAAACCCGATCAAGTGGAAGAAAAGAAGTGCATTCCGGCTGCTGACTTCAACCAG   4380
211      TTGCAAAAACCCGATCAAGTGGAAGAAAAGAAGTGCATTCCGGCTGCTGACTTCAACCAG   4380
246      TTGCAAAAACCCGATCAAGTGGAAGAAAAGAAGTGCATTCCGGCTGCTGACTTCAACCAG   4380
353      TTGCAAAAACCCGATCAAGTGGAAGAAAAGAAGTGCATTCCGGCTGCTGACTTCAACCAG   4380
         ************************************************************

DAH      ACTGCCGGTTATTTAATAAAGCTCAACGAGGGCCTTTACAGCTTCAGGGTGCGAGCCAAT   4440
29B      ACTGCCGGTTATTTAATAAAGCTCAACGAGGGCCTTTACAGCTTCAGGGTGCGAGCCAAT   4440
e19      ACTGCCGGTTATTTAATAAAGCTCAACGAGGGCCTTTACAGCTTCAGGGTGCGAGCCAAT   4440
74e19    ACTGCCGGTTATTTAATAAAGCTCAACGAGGGCCTTTACAGCTTCAGGGTGCGAGCCAAT   4440
74       ACTGCCGGTTATTTAATAAAGCTCAACGAGGGCCTTTACAGCTTCAGGGTGCGAGCCAAT   4440
211      ACTGCCGGTTATTTAATAAAGCTCAACGAGGGCCTTTACAGCTTCAGGGTGCGAGCCAAT   4440
246      ACTGCCGGTTATTTAATAAAGCTCAACGAGGGCCTTTACAGCTTCAGGGTGCGAGCCAAT   4440
353      ACTGCCGGTTATTTAATAAAGCTCAACGAGGGCCTTTACAGCTTCAGGGTGCGAGCCAAT   4440
         ************************************************************

DAH      TCAATAGCGGGATACGGCGATTTCACGGAAGTCGAACATATAAAAGTTGAGgtaggttga   4500
29B      TCAATAGCGGGATACGGCGATTTCACGGAAGTCGAACATATAAAAGTTGAGgtaggttga   4500
e19      TCAATAGCGGGATACGGCGATTTCACGGAAGTCGAACATATAAAAGTTGAGgtaggttga   4500
74e19    TCAATAGCGGGATACGGCGATTTCACGGAAGTCGAACATATAAAAGTTGAGgtaggttga   4500
74       TCAATAGCGGGATACGGCGATTTCACGGAAGTCGAACATATAAAAGTTGAGgtaggttga   4500
211      TCAATAGCGGGATACGGCGATTTCACGGAAGTCGAACATATAAAAGTTGAGgtaggttga   4500
246      TCAATAGCGGGATACGGCGATTTCACGGAAGTCGAACATATAAAAGTTGAGgtaggttga   4500
353      TCAATAGCGGGATACGGCGATTTCACGGAAGTCGAACATATAAAAGTTGAGgtaggttga   4500
         ************************************************************
```

```
DAH       ataatttttatctgc---------------------------------------------   4515
29B       ataatttttatctgcggtaccctagactagtctagataacttcgtataatgtatgctata   4560
e19       ataatttttatctgcggtaccctagactagtctagataacttcgtataatgtatgctata   4560
74e19     ataatttttatctgcggtaccctagactagtctagataacttcgtataatgtatgctata   4560
74        ataatttttatctgcggtaccctagactagtctagataacttcgtataatgtatgctata   4560
211       ataatttttatctgcggtaccctagactagtctagataacttcgtataatgtatgctata   4560
246       ataatttttatctgcggtaccctagactagtctagataacttcgtataatgtatgctata   4560
353       ataatttttatctgcggtaccctagactagtctagataacttcgtataatgtatgctata   4560
          ***************

DAH       -------------------------------gaatgatcatttttaaatgaatttttttt   4544
29B       cgaagttatctagactagtctagggcgcgccgaatgatcatttttaaatgaatttttttt   4620
e19       cgaagttatctagactagtctagggcgcgccgaatgatcatttttaaatgaatttttttt   4620
74e19     cgaagttatctagactagtctagggcgcgccgaatgatcatttttaaatgaatttttttt   4620
74        cgaagttatctagactagtctagggcgcgccgaatgatcatttttaaatgaatttttttt   4620
211       cgaagttatctagactagtctagggcgcgccgaatgatcatttttaaatgaatttttttt   4620
246       cgaagttatctagactagtctagggcgcgccgaatgatcatttttaaatgaatttttttt   4620
353       cgaagttatctagactagtctagggcgcgccgaatgatcatttttaaatgaatttttttt   4620
                                         *****************************

DAH       tttcattaaattcaacagCCTCCGCCGAGCTATGCTAAGGTCTTTTTCTGGCTACTGGGA   4604
29B       tttcattaaattcaacagCCTCCGCCGAGCTATGCTAAGGTCTTTTTCTGGCTACTGGGA   4680
e19       tttcattaaattcaacagCCTCCGCCGAGCTATGCTAAGGTCTTTTTCTGGCTACTGGGA   4680
74e19     tttcattaaattcaacagCCTCCGCCGAGCTATGCTAAGGTCTTTTTCTGGCTACTGGGA   4680
74        tttcattaaattcaacagCCTCCGCCGAGCTATGCTAAGGTCTTTTTCTGGCTACTGGGA   4680
211       tttcattaaattcaacagCCTCCGCCGAGCTATGCTAAGGTCTTTTTCTGGCTACTGGGA   4680
246       tttcattaaattcaacagCCTCCGCCGAGCTATGCTAAGGTCTTTTTCTGGCTACTGGGA   4680
353       tttcattaaattcaacagCCTCCGCCGAGCTATGCTAAGGTCTTTTTCTGGCTACTGGGA   4680
          ************************************************************

DAH       ATCGGCCTAGCGTTCCTGATCGTTTCCCTGTTCGGCTATGTCTGTTACCTGCACAAGAGG   4664
29B       ATCGGCCTAGCGTTCCTGATCGTTTCCCTGTTCGGCTATGTCTGTTACCTGCACAAGAGG   4740
e19       ATCGGCCTAGCGTTCCTGATCGTTTCCCTGTTCGGCTATGTCTGTTACCTGCACAAGAGG   4740
74e19     ATCGGCCTAGCGTTCCTGATCGTTTCCCTGTTCGGCTATGTCTGTTACCTGCACAAGAGG   4740
74        ATCGGCCTAGCGTTCCTGATCGTTTCCCTGTTCGGCTATGTCTGTTACCTGCACAAGAGG   4740
211       ATCGGCCTAGCGTTCCTGATCGTTTCCCTGTTCGGCTATGTCTGTTACCTGCACAAGAGG   4740
246       ATCGGCCTAGCGTTCCTGATCGTTTCCCTGTTCGGCTATGTCTGTTACCTGCACAAGAGG   4740
353       ATCGGCCTAGCGTTCCTGATCGTTTCCCTGTTCGGCTATGTCTGTTACCTGCACAAGAGG   4740
          ************************************************************

DAH       AAGGTTCCCTCTAATGACCTCCATATGAACACAGAGGTGAATCCGTTCTATGCGAGCATG   4724
29B       AAGGTTCCCTCTAATGACCTCCATATGAACACAGAGGTGAATCCGTTCTATGCGAGCATG   4800
e19       AAGGTTCCCTCTAATGACCTCCATATGAACACAGAGGTGAATCCGTTCTATGCGAGCATG   4800
74e19     AAGGTTCCCTCTAATGACCTCCATATGAACACAGAGGTGAATCCGTTCTATGCGAGCATG   4800
74        AAGGTTCCCTCTAATGACCTCCATATGAACACAGAGGTGAATCCGTTCTATGCGAGCATG   4800
211       AAGGTTCCCTCTAATGACCTCCATATGAACACAGAGGTGAATCCGTTCTATGCGAGCATG   4800
246       AAGGTTCCCTCTAATGACCTCCATATGAACACAGAGGTGAATCCGTTCTATGCGAGCATG   4800
353       AAGGTTCCCTCTAATGACCTCCATATGAACACAGAGGTGAATCCGTTCTATGCGAGCATG   4800
          ************************************************************

DAH       CAATACATCCCAGACGATTGGGAGGTGCTGCGAGAGAACATCATTCAGTTGGCTCCACTA   4784
29B       CAATACATCCCAGACGATTGGGAGGTGCTGCGAGAGAACATCATTCAGTTGGCTCCACTA   4860
e19       CAATACATCCCAGACGATTGGGAGGTGCTGCGAGAGAACATCATTCAGTTGGCTCCACTA   4860
74e19     CAATACATCCCAGACGATTGGGAGGTGCTGCGAGAGAACATCATTCAGTTGGCTCCACTA   4860
74        CAATACATCCCAGACGATTGGGAGGTGCTGCGAGAGAACATCATTCAGTTGGCTCCACTA   4860
211       CAATACATCCCAGACGATTGGGAGGTGCTGCGAGAGAACATCATTCAGTTGGCTCCACTA   4860
246       CAATACATCCCAGACGATTGGGAGGTGCTGCGAGAGAACATCATTCAGTTGGCTCCACTA   4860
353       CAATACATCCCAGACGATTGGGAGGTGCTGCGAGAGAACATCATTCAGTTGGCTCCACTA   4860
          ************************************************************
```

```
DAH      GGCCAGGGATCCTTTGGCATGGTGTATGAGGGTATCCTGAAGTCCTTTCCACCCAATGGC   4844
29B      GGCCAGGGATCCTTTGGCATGGTGTATGAGGGTATCCTGAAGTCCTTTCCACCCAATGGC   4920
e19      GGCCAGGGATCCTTTGGCATGGTGTATGAGGGTATCCTGAAGTCCTTTCCACCCAATGGC   4920
74e19    GGCCAGGGATCCTTTGGCATGGTGTATGAGGGTATCCTGAAGTCCTTTCCACCCAATGGC   4920
74       GGCCAGGGATCCTTTGGCATGGTGTATGAGGGTATCCTGAAGTCCTTTCCACCCAATGGC   4920
211      GGCCAGGGATCCTTTGGCATGGTGTATGAGGGTATCCTGAAGTCCTTTCCACCCAATGGC   4920
246      GGCCAGGGATCCTTTGGCATGATGTATGAGGGTATCCTGAAGTCCTTTCCACCCAATGGC   4920
353      GGCCAGGGATCCTTTGGCATGGTGTATGAGGGTATCCTGAAGTCCTTTCCACCCAATGGC   4920
         ********************* **************************************

DAH      GTGGATCGCGAGTGTGCCATTAAGACTGTCAACGAAAATGCTACGGATCGCGAGCGAACC   4904
29B      GTGGATCGCGAGTGTGCCATTAAGACTGTCAACGAAAATGCTACGGATCGCGAGCGAACC   4980
e19      GTGGATCGCGAGTGTGCCATTAAGACTGTCAACGAAAATGCTACGGATCGCGAGCGAACC   4980
74e19    GTGGATCGCGAGTGTGCCATTAAGACTGTCAACGAAAATGCTACGGATCGCGAGCGAACC   4980
74       GTGGATCGCGAGTGTGCCATTAAGACTGTCAACGAAAATGCTACGGATCGCGAGCGAACC   4980
211      GTGGATCGCGAGTGTGCCATTAAGACTGTCAACGAAAATGCTACGGATCGCGAGCGAACC   4980
246      GTGGATCGCGAGTGTGCCATTAAGACTGTCAACGAAAATGCTACGGATCGCGAGCGAACC   4980
353      GTGGATCGCGAGTGTGCCATTAAGACTGTCAACGAAAATGCTACGGATCGCGAGCGAACC   4980
         ************************************************************

DAH      AATTTCCTGAGCGAGGCGAGCGTCATGAAGGAGTTCGATACGTATCATGTCGTAAGATTG   4964
29B      AATTTCCTGAGCGAGGCGAGCGTCATGAAGGAGTTCGATACGTATCATGTCGTAAGATTG   5040
e19      AATTTCCTGAGCGAGGCGAGCGTCATGAAGGAGTTCGATACGTATCATGTCGTAAGATTG   5040
74e19    AATTTCCTGAGCGAGGCGAGCGTCATGAAGGAGTTCGATACGTATCATGTCGTAAGATTG   5040
74       AATTTCCTGAGCGAGGCGAGCGTCATGAAGGAGTTCGATACGTATCATGTCGTAAGATTG   5040
211      AATTTCCTGAGCGAGGCGAGCGTCATGAAGGAGTTCGATACGTATCATGTCGTAAGATTG   5040
246      AATTTCCTGAGCGAGGCGAGCGTCATGAAGGAGTTCGATACGTATCATGTCGTAAGATTG   5040
353      AATTTCCTGAGCGAGGCGAGCGTCATGAAGGAGTTCGATACGTATCATGTCGTAAGATTG   5040
         ************************************************************

DAH      CTCGGTGTTTGTTCCAGGGGTCAGCCGGCTCTGGTGGTCATGGAGCTAATGAAGAAGGGT   5024
29B      CTCGGTGTTTGTTCCAGGGGTCAGCCGGCTCTGGTGGTCATGGAGCTAATGAAGAAGGGT   5100
e19      CTCGGTGTTTGTTCCAGGGGTCAGCCGGCTCTGGTGGTCATGGAGCTAATGAAGAAGGGT   5100
74e19    CTCGGTGTTTGTTCCAGGGGTCAGCCGGCTCTGGTGGTCATGGAGCTAATGAAGAAGGGT   5100
74       CTCGGTGTTTGTTCCAGGGGTCAGCCGGCTCTGGTGGTCATGGAGCTAATGAAGAAGGGT   5100
211      CTCGGTGTTTGTTCCAGGGGTCAGCCGGCTCTGGTGGTCATGGAGCTAATGAAGAAGGGT   5100
246      CTCGGTGTTTGTTCCAGGGGTCAGCCGGCTCTGGTGGTCATGGAGCTAATGAAGAAGGGT   5100
353      CTCGGTGTTTGTTCCAGGGGTCAGCCGGCTCTGGTGGTCATGGAGCTAATGAAGAAGGGT   5100
         ************************************************************

DAH      GATCTTAAGTCCTATTTGCGTGCCCATCGTCCCGAGGAGCGGGATGAGGCCATGATGACG   5084
29B      GATCTTAAGTCCTATTTGCGTGCCCATCGTCCCGAGGAGCGGGATGAGGCCATGATGACG   5160
e19      GATCTTAAGTCCTATTTGCGTGCCCATCGTCCCGAGGAGCGGGATGAGGCCATGATGACG   5160
74e19    GATCTTAAGTCCTATTTGCGTGCCCATCGTCCCGAGGAGCGGGATGAGGCCATGATGACG   5160
74       GATCTTAAGTCCTATTTGCGTGCCCATCGTCCCGAGGAGCGGGATGAGGCCATGATGACG   5160
211      GATCTTAAGTCCTATTTGCGTGCCCATCGTCCCGAGGAGCGGGATGAGGCCATGATGACG   5160
246      GATCTTAAGTCCTATTTGCGTGCCCATCGTCCCGAGGAGCGGGATGAGGCCATGATGACG   5160
353      GATCTTAAGTCCTATTTGCGTGCCCATTGTCCCGAGGAGCGGGATGAGGCCATGATGACG   5160
         *************************** ********************************

DAH      TATCTTAATCGCATCGGAGTGACTGGTAATGTGCAGCCTCCTACTTATGGAAGAATCTAC   5144
29B      TATCTTAATCGCATCGGAGTGACTGGTAATGTGCAGCCTCCTACTTATGGAAGAATCTAC   5220
e19      TATCTTAATCGCATCGGAGTGACTGGTAATGTGCAGCCTCCTACTTATGGAAGAATCTAC   5220
74e19    TATCTTAATCGCATCGGAGTGACTGGTAATGTGCAGCCTCCTACTTATGGAAGAATCTAC   5220
74       TATCTTAATCGCATCGGAGTGACTGGTAATGTGCAGCCTCCTACTTATGGAAGAATCTAC   5220
211      TATCTTAATCGCATCGGAGTGACTGGTAATGTGCAGCCTCCTACTTATGGAAGAATCTAC   5220
246      TATCTTAATCGCATCGGAGTGACTGGTAATGTGCAGCCTCCTACTTATGGAAGAATCTAC   5220
353      TATCTTAATCGCATCGGAGTGACTGGTAATGTGCAGCCTCCTACTTATGGAAGAATCTAC   5220
         ************************************************************
```

```
DAH      CAGATGGCCATTGAGATTGCGGATGGCATGGCATATTTGGCCGCCAAGAAGTTCGTCCAT   5204
29B      CAGATGGCCATTGAGATTGCGGATGGCATGGCATATTTGGCCGCCAAGAAGTTCGTCCAT   5280
e19      CAGATGGCCATTGAGATTGCGGATGGCATGGCATATTTGGCCGCCAAGAAGTTCGTCCAT   5280
74e19    CAGATGGCCATTGAGATTGCGGATGGCATGGCATATTTGGCCGCCAAGAAGTTCGTCCAT   5280
74       CAGATGGCCATTGAGATTGCGGATGGCATGGCATATTTGGCCGCCAAGAAGTTCGTCCAT   5280
211      CAGATGGCCATTGAGATTGCGGATGGCATGGCATATTTGGCCGCCAAGAAGTTCGTCCAT   5280
246      CAGATGGCCATTGAGATTGCGGATGGCATGGCATATTTGGCCGCCAAGAAGTTCGTCCAT   5280
353      CAGATGGCCATTGAGATTGCGGATGGCATGGCATATTTGGCCGCCAAGAAGTTCGTCCAT   5280
         ************************************************************

DAH      CGTGATCTTGCAGCTCGAAATTGCATGGTTGCTGATGATTTGACGGTGAAAATTGGTGAC   5264
29B      CGTGATCTTGCAGCTCGAAATTGCATGGTTGCTGATGATTTGACGGTGAAAATTGGTGAC   5340
e19      CGTGATCTTGCAGCTCGAAATTGCATGGTTGCTGATGATTTGACGGTGAAAATTGGTGAC   5340
74e19    CGTGATCTTGCAGCTCGAAATTGCATGGTTGCTGATGATTTGACGGTGAAAATTGGTGAC   5340
74       CGTGATCTTGCAGCTCGAAATTGCATGGTTGCTGATGATTTGACGGTGAAAATTGGTGAC   5340
211      CGTGATCTTGCAGCTCGAAATTGCATGGTTGCTGATGATTTGACGGTGAAAATTGGTGAC   5340
246      CGTGATCTTGCAGCTCGAAATTGCATGGTTGCTGATGATTTGACGGTGAAAATTGGTGAC   5340
353      CGTGATCTTGCAGCTCGAAATTGCATGGTTGCTGATGATTTGACGGTGAAAATTGGTGAC   5340
         ************************************************************

DAH      TTTGGAATGACCCGTGACATCTATGAGACGGATTACTATCGGAAGGGCACTAAAGGGCTG   5324
29B      TTTGGAATGACCCGTGACATCTATGAGACGGATTACTATCGGAAGGGCACTAAAGGGCTG   5400
e19      TTTGGAATGACCCGTGACATCTATGAGACGGATTACTATCGGAAGGGCACTAAAGGGCTG   5400
74e19    TTTGGAATGACCCGTGACTTCTATGAGACGGATTACTATCGGAAGGGCACTAAAGGGCTG   5400
74       TTTGGAATGACCCGTGACTTCTATGAGACGGATTACTATCGGAAGGGCACTAAAGGGCTG   5400
211      TTTGGAATGACCCGTGACATCTATGAGACGGATTACTATCGGAAGGGCACTAAAGGGCTG   5400
246      TTTGGAATGACCCGTGACATCTATGAGACGGATTACTATCGGAAGGGCACTAAAGGGCTG   5400
353      TTTGGAATGACCCGTGACATCTATGAGACGGATTACTATCGGAAGGGCACTAAAGGGCTG   5400
         ****************** *****************************************

DAH      CTGCCAGTTCGCTGGATGCCACCGGAGAGCTTGCGAGATGGTGTCTACTCTAGTGCCAGT   5384
29B      CTGCCAGTTCGCTGGATGCCACCGGAGAGCTTGCGAGATGGTGTCTACTCTAGTGCCAGT   5460
e19      CTGCCAGTTCGCTGGATGCCACCGGAGAGCTTGCGAGATGGTGTCTACTCTAGTGCCAGT   5460
74e19    CTGCCAGTTCGCTGGATGCCACCGGAGAGCTTGCGAGATGGTGTCTACTCTAGTGCCAGT   5460
74       CTGCCAGTTCGCTGGATGCCACCGGAGAGCTTGCGAGATGGTGTCTACTCTAGTGCCAGT   5460
211      CTGCCAGTTCGCTGGATGCCACCGGAGAGCTTGCGAGATGGTGTCTACTCTAGTGCCAGT   5460
246      CTGCCAGTTCGCTGGATGCCACCGGAGAGCTTGCGAGATGGTGTCTACTCTAGTGCCAGT   5460
353      CTGCCAGTTCGCTGGATGCCACCGGAGAGCTTGCGAGATGGTGTCTACTCTAGTGCCAGT   5460
         ************************************************************

DAH      GATGTATTCAGCTTTGGAGTGGTTCTCTGGGAAATGGCCACCTTAGCGGCTCAGCCATAC   5444
29B      GATGTATTCAGCTTTGGAGTGGTTCTCTGGGAAATGGCCACCTTAGCGGCTCAGCCATAC   5520
e19      GATGTATTCAGCTTTGGAGTGGTTCTCTGGGAAATGGCCACCTTAGCGGCTCAGCCATAC   5520
74e19    GATGTATTCAGCTTTGGAGTGGTTCTCTGGGAAATGGCCACCTTAGCGGCTCAGCCATAC   5520
74       GATGTATTCAGCTTTGGAGTGGTTCTCTGGGAAATGGCCACCTTAGCGGCTCAGCCATAC   5520
211      GATGTATTCAGCTTTGGAGTGGTTCTCTGGGAAATGGCCACCTTAGCGGCTCAGCCATAC   5520
246      GATGTATTCAGCTTTGGAGTGGTTCTCTGGGAAATGGCCACCTTAGCGGCTCAGCCATAC   5520
353      GATGTATTCAGCTTTGGAGTGGTTCTCTGGGAAATGGCCACCTTAGCGGCTCAGCCATAC   5520
         ************************************************************

DAH      CAGGGACTTTCCAACGAGCAAGTCCTGCGTTACGTTATCGATGGCGGTGTTATGGAGAGG   5504
29B      CAGGGACTTTCCAACGAGCAAGTCCTGCGTTACGTTATCGATGGCGGTGTTATGGAGAGG   5580
e19      CAGGGACTTTCCAACGAGCAAGTCCTGCGTTACGTTATCGATGGCGGTGTTATGGAGAGG   5580
74e19    CAGGGACTTTCCAACGAGCAAGTCCTGCGTTACGTTATCGATGGCGGTGTTATGGAGAGG   5580
74       CAGGGACTTTCCAACGAGCAAGTCCTGCGTTACGTTATCGATGGCGGTGTTATGGAGAGG   5580
211      CAGAGACTTTCCAACGAGCAAGTCCTGCGTTACGTTATCGATGGCGGTGTTATGGAGAGG   5580
246      CAGGGACTTTCCAACGAGCAAGTCCTGCGTTACGTTATCGATGGCGGTGTTATGGAGAGG   5580
353      CAGGGACTTTCCAACGAGCAAGTCCTGCGTTACGTTATCGATGGCGGTGTTATGGAGAGG   5580
         *** ********************************************************
```

```
DAH       CCGGAAAATTGTCCTGATTTTCTGCATAAACTAATGCAAAGGTGCTGGCATCATAGGTCT    5564
29B       CCGGAAAATTGTCCTGATTTTCTGCATAAACTAATGCAAAGGTGCTGGCATCATAGGTCT    5640
e19       CCGGAAAATTGTCCTGATTTTCTGCATAAACTAATGCAAAGGTGCTGGCATCATAGGTCT    5640
74e19     CCGGAAAATTGTCCTGATTTTCTGCATAAACTAATGCAAAGGTGCTGGCATCATAGGTCT    5640
74        CCGGAAAATTGTCCTGATTTTCTGCATAAACTAATGCAAAGGTGCTGGCATCATAGGTCT    5640
211       CCGGAAAATTGTCCTGATTTTCTGCATAAACTAATGCAAAGGTGCTGGCATCATAGGTCT    5640
246       CCGGAAAATTGTCCTGATTTTCTGCATAAACTAATGCAAAGGTGCTGGCATCATAGGTCT    5640
353       CCGGAAAATTGTCCTGATTTTCTGCATAAACTAATGCAAAGGTGCTGGCATCATAGGTCT    5640
          ************************************************************

DAH       TCGGCGAGACCCAGTTTTCTGGATATCATTGCGTATCTCGAACCACAATGCCCCAATTCA    5624
29B       TCGGCGAGACCCAGTTTTCTGGATATCATTGCGTATCTCGAACCACAATGCCCCAATTCA    5700
e19       TCGGCGAGACCCAGTTTTCTGGATATCATTGCGTATCTCGAACCACAATGCCCCAATTCA    5700
74e19     TCGGCGAGACCCAGTTTTCTGGATATCATTGCGTATCTCGAACCACAATGCCCCAATTCA    5700
74        TCGGCGAGACCCAGTTTTCTGGATATCATTGCGTATCTCGAACCACAATGCCCCAATTCA    5700
211       TCGGCGAGACCCAGTTTTCTGGATATCATTGCGTATCTCGAACCACAATGCCCCAATTCA    5700
246       TCGGCGAGACCCAGTTTTCTGGATATCATTGCGTATCTCGAACCACAATGCCCCAATTCA    5700
353       TCGGCGAGACCCAGTTTTCTGGATATCATTGCGTATCTCGAACCACAATGCCCCAATTCA    5700
          ************************************************************

DAH       CAATTTAAGGAAGTATCCTTCTATCACTCAGAGGCAGGTCTGCAGCATCGGGAAAAGGAG    5684
29B       CAATTTAAGGAAGTATCCTTCTATCACTCAGAGGCAGGTCTGCAGCATCGGGAAAAGGAG    5760
e19       CAATTTAAGGAAGTATCCTTCTATCACTCAGAGGCAGGTCTGCAGCATCGGGAAAAGGAG    5760
74e19     CAATTTAAGGAAGTATCCTTCTATCACTCAGAGGCAGGTCTGCAGCATCGGGAAAAGGAG    5760
74        CAATTTAAGGAAGTATCCTTCTATCACTCAGAGGCAGGTCTGCAGCATCGGGAAAAGGAG    5760
211       CAATTTAAGGAAGTATCCTTCTATCACTCAGAGGCAGGTCTGCAGCATCGGGAAAAGGAG    5760
246       CAATTTAAGGAAGTATCCTTCTATCACTCAGAGGCAGGTCTGCAGCATCGGGAAAAGGAG    5760
353       CAATTTAAGGAAGTATCCTTCTATCACTCAGAGGCAGGTCTGCAGCATCGGGAAAAGGAG    5760
          ************************************************************

DAH       CGCAAGGAACGCAATCAGCTAGATGCATTCGCGGCAGTCCCCTTGGATCAAGATCTGCAG    5744
29B       CGCAAGGAACGCAATCAGCTAGATGCATTCGCGGCAGTCCCCTTGGATCAAGATCTGCAG    5820
e19       CGCAAGGAACGCAATCAGCTAGATGCATTCGCGGCAGTCCCCTTGGATCAAGATCTGCAG    5820
74e19     CGCAAGGAACGCAATCAGCTAGATGCATTCGCGGCAGTCCCCTTGGATCAAGATCTGCAG    5820
74        CGCAAGGAACGCAATCAGCTAGATGCATTCGCGGCAGTCCCCTTGGATCAAGATCTGCAG    5820
211       CGCAAGGAACGCAATCAGCTAGATGCATTCGCGGCAGTCCCCTTGGATCAAGATCTGCAG    5820
246       CGCAAGGAACGCAATCAGCTAGATGCATTCGCGGCAGTCCCCTTGGATCAAGATCTGCAG    5820
353       CGCAAGGAACGCAATCAGCTAGATGCATTCGCGGCAGTCCCCTTGGATCAAGATCTGCAG    5820
          ************************************************************

DAH       GATCGGGAACAGCAGGAGGATGCTACCACACCTTTACGAATGGGCGATTATCAGCAGAAC    5804
29B       GATCGGGAACAGCAGGAGGATGCTACCACACCTTTACGAATGGGCGATTATCAGCAGAAC    5880
e19       GATCGGGAACAGCAGGAGGATGCTACCACACCTTTACGAATGGGCGATTATCAGCAGAAC    5880
74e19     GATCGGGAACAGCAGGAGGATGCTACCACACCTTTACGAATGGGCGATTATCAGCAGAAC    5880
74        GATCGGGAACAGCAGGAGGATGCTACCACACCTTTACGAATGGGCGATTATCAGCAGAAC    5880
211       GATCGGGAACAGCAGGAGGATGCTACCACACCTTTACGAATGGGCGATTATCAGCAGAAC    5880
246       GATCGGGAACAGCAGGAGGATGCTACCACACCTTTACGAATGGGCGATTATCAGCAGAAC    5880
353       GATCGGGAACAGCAGGAGGATGCTACCACACCTTTACGAATGGGCGATTATCAGCAGAAC    5880
          ************************************************************

DAH       TCCTCGTTGGATCAACCGCCCGAAAGCCCCATCGCCATGGTTGATGATCAGGGTTCTCAC    5864
29B       TCCTCGTTGGATCAACCGCCCGAAAGCCCCATCGCCATGGTTGATGATCAGGGTTCTCAC    5940
e19       TCCTCGTTGGATCAACCGCCCGAAAGCCCCATCGCCATGGTTGATGATCAGGGTTCTCAC    5940
74e19     TCCTCGTTGGATCAACCGCCCGAAAGCCCCATCGCCATGGTTGATGATCAGGGTTCTCAC    5940
74        TCCTCGTTGGATCAACCGCCCGAAAGCCCCATCGCCATGGTTGATGATCAGGGTTCTCAC    5940
211       TCCTCGTTGGATCAACCGCCCGAAAGCCCCATCGCCATGGTTGATGATCAGGGTTCTCAC    5940
246       TCCTCGTTGGATCAACCGCCCGAAAGCCCCATCGCCATGGTTGATGATCAGGGTTCTCAC    5940
353       TCCTCGTTGGATCAACCGCCCGAAAGCCCCATCGCCATGGTTGATGATCAGGGTTCTCAC    5940
          ************************************************************
```

```
DAH       TTGCCATTTAGCCTGCCATCCGGATTCATTGCGAGCAGTACTCCTGATGGGCAGACTGTA   5924
29B       TTGCCATTTAGCCTGCCATCCGGATTCATTGCGAGCAGTACTCCTGATGGGCAGACTGTA   6000
e19       TTGCCATTTAGCCTGCCATCCGGATTCATTGCGAGCAGTACTCCTGATGGGCAGACTGTA   6000
74e19     TTGCCATTTAGCCTGCCATCCGGATTCATTGCGAGCAGTACTCCTGATGGGCAGACTGTA   6000
74        TTGCCATTTAGCCTGCCATCCGGATTCATTGCGAGCAGTACTCCTGATGGGCAGACTGTA   6000
211       TTGCCATTTAGCCTGCCATCCGGATTCATTGCGAGCAGTACTCCTGATGGGCAGACTGTA   6000
246       TTGCCATTTAGCCTGCCATCCGGATTCATTGCGAGCAGTACTCCTGATGGGCAGACTGTA   6000
353       TTGCCATTTAGCCTGCCATCCGGATTCATTGCGAGCAGTACTCCTGATGGGCAGACTGTA   6000
          ************************************************************

DAH       ATGGCTACTGCTTTCCAGAATATTCCAGCAGCGCAAGGTGATATTTCGGCGACTTACGTT   5984
29B       ATGGCTACTGCTTTCCAGAATATTCCAGCAGCGCAAGGTGATATTTCGGCGACTTACGTT   6060
e19       ATGGCTACTGCTTTCCAGAATATTCCAGCAGCGCAAGGTGATATTTCGGCGACTTACGTT   6060
74e19     ATGGCTACTGCTTTCCAGAATATTCCAGCAGCGCAAGGTGATATTTCGGCGACTTACGTT   6060
74        ATGGCTACTGCTTTCCAGAATATTCCAGCAGCGCAAGGTGATATTTCGGCGACTTACGTT   6060
211       ATGGCTACTGCTTTCCAGAATATTCCAGCAGCGCAAGGTGATATTTCGGCGACTTACGTT   6060
246       ATGGCTACTGCTTTCCAGAATATTCCAGCAGCGCAAGGTGATATTTCGGCGACTTACGTT   6060
353       ATGGCTACTGCTTTCCAGAATATTCCAGCAGCGCAAGGTGATATTTCGGCGACTTACGTT   6060
          ************************************************************

DAH       GTGCCTGATGCAGACGCTTTGGACGGCGACAGGGGATATGAGATCTACGATCCCAGTCCG   6044
29B       GTGCCTGATGCAGACGCTTTGGACGGCGACAGGGGATATGAGATCTACGATCCCAGTCCG   6120
e19       GTGCCTGATGCAGACGCTTTGGACGGCGACAGGGGATATGAGATCTACGATCCCAGTCCG   6120
74e19     GTGCCTGATGCAGACGCTTTGGACGGCGACAGGGGATATGAGATCTACGATCCCAGTCCG   6120
74        GTGCCTGATGCAGACGCTTTGGACGGCGACAGGGGATATGAGATCTACGATCCCAGTCCG   6120
211       GTGCCTGATGCAGACGCTTTGGACGGCGACAGGGGATATGAGATCTACGATCCCAGTCCG   6120
246       GTGCCTGATGCAGACGCTTTGGACGGCGACAGGGGATATGAGATCTACGATCCCAGTCCG   6120
353       GTGCCTGATGCAGACGCTTTGGACGGCGACAGGGGATATGAGATCTACGATCCCAGTCCG   6120
          ************************************************************

DAH       AAATGTGCAGAGCTGCCGACGAGCAGAAGTGGCAGTACTGGCGGTGGAAAACTCAGCGGA   6104
29B       AAATGTGCAGAGCTGCCGACGAGCAGAAGTGGCAGTACTGGCGGTGGAAAACTCAGCGGA   6180
e19       AAATGTGCAGAGCTGCCGACGAGCAGAAGTGGCAGTACTGGCGGTGGAAAACTCAGCGGA   6180
74e19     AAATGTGCAGAGCTGCCGACGAGCAGAAGTGGCAGTACTGGCGGTGGAAAACTCAGCGGA   6180
74        AAATGTGCAGAGCTGCCGACGAGCAGAAGTGGCAGTACTGGCGGTGGAAAACTCAGCGGA   6180
211       AAATGTGCAGAGCTGCCGACGAGCAGAAGTGGCAGTACTGGCGGTGGAAAACTCAGCGGA   6180
246       AAATGTGCAGAGCTGCCGACGAGCAGAAGTGGCAGTACTGGCGGTGGAAAACTCAGCGGA   6180
353       AAATGTGCAGAGCTGCCGACGAGCAGAAGTGGCAGTACTGGCGGTGGAAAACTCAGCGGA   6180
          ************************************************************

DAH       GAACAACATTTGCTGCCAAGAAAGGGTCGCCAGCCTACCATCATGAGCAGCTCGATGCCA   6164
29B       GAACAACATTTGCTGCCAAGAAAGGGTCGCCAGCCTACCATCATGAGCAGCTCGATGCCA   6240
e19       GAACAACATTTGCTGCCAAGAAAGGGTCGCCAGCCTACCATCATGAGCAGCTCGATGCCA   6240
74e19     GAACAACATTTGCTGCCAAGAAAGGGTCGCCAGCCTACCATCATGAGCAGCTCGATGCCA   6240
74        GAACAACATTTGCTGCCAAGAAAGGGTCGCCAGCCTACCATCATGAGCAGCTCGATGCCA   6240
211       GAACAACATTTGCTGCCAAGAAAGGGTCGCCAGCCTACCATCATGAGCAGCTCGATGCCA   6240
246       GAACAACATTTGCTGCCAAGAAAGGGTCGCCAGCCTACCATCATGAGCAGCTCGATGCCA   6240
353       GAACAACATTTGCTGCCAAGAAAGGGTCGCCAGCCTACCATCATGAGCAGCTCGATGCCA   6240
          ************************************************************

DAH       GATGATGTCATCGGTGGGTCCTCACTGCAACCCTCGACTGCGTCAGCAGCCAGTTCTAAT   6224
29B       GATGATGTCATCGGTGGGTCCTCACTGCAACCCTCGACTGCGTCAGCAGCCAGTTCTAAT   6300
e19       GATGATGTCATCGGTGGGTCCTCACTGCAACCCTCGACTGCGTCAGCAGCCAGTTCTAAT   6300
74e19     GATGATGTCATCGGTGGGTCCTCACTGCAACCCTCGACTGCGTCAGCAGCCAGTTCTAAT   6300
74        GATGATGTCATCGGTGGGTCCTCACTGCAACCCTCGACTGCGTCAGCAGCCAGTTCTAAT   6300
211       GATGATGTCATCGGTGGGTCCTCAGTGCAACCCTCGACTGCGTCAGCAGCCAGTTCTAAT   6300
246       GATGATGTCATCGGTGGGTCCTCACTGCAACCCTCGACTGCGTCAGCAGCCAGTTCTAAT   6300
353       GATGATGTCATCGGTGGGTCCTCACTGCAACCCTCGACTGCGTCAGCAGCCAGTTCTAAT   6300
          ************************ ***********************************
```

```
DAH      GCCAGTTCGCACACAGGACGCCCAAGTCTGAAGAAAACAGTGGCGGATTCGGTTCGCAAT    6284
29B      GCCAGTTCGCACACAGGACGCCCAAGTCTGAAGAAAACAGTGGCGGATTCGGTTCGCAAT    6360
e19      GCCAGTTCGCACACAGGACGCCCAAGTCTGAAGAAAACAGTGGCGGATTCGGTTCGCAAT    6360
74e19    GCCAGTTCGCACACAGGACGCCCAAGTCTGAAGAAAACAGTGGCGGATTCGGTTCGCAAT    6360
74       GCCAGTTCGCACACAGGACGCCCAAGTCTGAAGAAAACAGTGGCGGATTCGGTTCGCAAT    6360
211      GCCAGTTCGCACACAGGACGCCCAAGTCTGAAGAAAACAGTGGCGGATTCGGTTCGCAAT    6360
246      GCCAGTTCGCACACAGGACGCCCAAGTCTGAAGAAAACAGTGGCGGATTCGGTTCGCAAT    6360
353      GCCAGTTCGCACACAGGACGCCCAAGTCTGAAGAAAACAGTGGCGGATTCGGTTCGCAAT    6360
         ************************************************************

DAH      AAGGCAAACTTCATTAATCGCCACCTATTTAACCACAAGCGAACGGGCAGCAATGCCAGC    6344
29B      AAGGCAAACTTCATTAATCGCCACCTATTTAACCACAAGCGAACGGGCAGCAATGCCAGC    6420
e19      AAGGCAAACTTCATTAATCGCCACCTATTTAACCACAAGCGAACGGGCAGCAATGCCAGC    6420
74e19    AAGGCAAACTTCATTAATCGCCACCTATTTAACCACAAGCGAACGGGCAGCAATGCCAGC    6420
74       AAGGCAAACTTCATTAATCGCCACCTATTTAACCACAAGCGAACGGGCAGCAATGCCAGC    6420
211      AAGGCAAACTTCATTAATCGCCACCTATTTAACCACAAGCGAACGGGCAGCAATGCCAGC    6420
246      AAGGCAAACTTCATTAATCGCCACCTATTTAACCACAAGCGAACGGGCAGCAATGCCAGC    6420
353      AAGGCAAACTTCATTAATCGCCACCTATTTAACCACAAGCGAACGGGCAGCAATGCCAGC    6420
         ************************************************************

DAH      CACAAGAGCAATGCCTCCAATGCTCCGAGTACCAGCAGTAACACCAACTTGACAAGTCAC    6404
29B      CACAAGAGCAATGCCTCCAATGCTCCGAGTACCAGCAGTAACACCAACTTGACAAGTCAC    6480
e19      CACAAGAGCAATGCCTCCAATGCTCCGAGTACCAGCAGTAACACCAACTTGACAAGTCAC    6480
74e19    CACAAGAGCAATGCCTCCAATGCTCCGAGTACCAGCAGTAACACCAACTTGACAAGTCAC    6480
74       CACAAGAGCAATGCCTCCAATGCTCCGAGTACCAGCAGTAACACCAACTTGACAAGTCAC    6480
211      CACAAGAGCAATGCCTCCAATGCTCCGAGTACCAGCAGTAACACCAACTTGACAAGTCAC    6480
246      CACAAGAGCAATGCCTCCAATGCTCCGAGTACCAGCAGTAACACCAACTTGACAAGTCAC    6480
353      CACAAGAGCAATGCCTCCAATGCTCCGAGTACCAGCAGTAACACCAACTTGACAAGTCAC    6480
         ************************************************************

DAH      CCAGTGGCTATGGGCAATCTTGGAACTATCGAGAGTGGTGGCAGTGGTTCGGCTGGTAGT    6464
29B      CCAGTGGCTATGGGCAATCTTGGAACTATCGAGAGTGGTGGCAGTGGTTCGGCTGGTAGT    6540
e19      CCAGTGGCTATGGGCAATCTTGGAACTATCGAGAGTGGTGGCAGTGGTTCGGCTGGTAGT    6540
74e19    CCAGTGGCTATGGGCAATCTTGGAACTATCGAGAGTGGTGGCAGTGGTTCGGCTGGTAGT    6540
74       CCAGTGGCTATGGGCAATCTTGGAACTATCGAGAGTGGTGGCAGTGGTTCGGCTGGTAGT    6540
211      CCAGTGGCTGTGGGCAATCTTGGAACTATCGAGAGTGGTGGCAGTGGTTCGGCTGGTAGT    6540
246      CCAGTGGCTATGGGCAATCTTGGAACTATCGAGAGTGGTGGCAGTGGTTCGGCTGGTAGT    6540
353      CCAGTGGCTATGGGCAATCTTGGAACTATCGAGAGTGGTGGCAGTGGTTCGGCTGGTAGT    6540
         ********* **************************************************

DAH      TATACTGGAACACCCCGCTTCTATACTCCATCAGCGACGCCTGGAGGAGGCAGCGGTATG    6524
29B      TATACTGGAACACCCCGCTTCTATACTCCATCAGCGACGCCTGGAGGAGGCAGCGGTATG    6600
e19      TATACTGGAACACCCCGCTTCTATACTCCATCAGCGACGCCTGGAGGAGGCAGCGGTATG    6600
74e19    TATACTGGAACACCCCGCTTCTATACTCCATCAGCGACGCCTGGAGGAGGCAGCGGTATG    6600
74       TATACTGGAACACCCCGCTTCTATACTCCATCAGCGACGCCTGGAGGAGGCAGCGGTATG    6600
211      TATACTGGAACACCCCGCTTCTATACTCCATCAGCGACGCCTGGAGGAGGCAGCGGTATG    6600
246      TATACTGGAACACCCCGCTTCTATACTCCATCAGCGACGCCTGGAGGAGGCAGCGGTATG    6600
353      TATACTGGAACACCCCGCTTCTATACTCCATCAGCGACGCCTGGAGGAGGCAGCGGTATG    6600
         ************************************************************

DAH      GCCATTAGCGACAATCCTAACTACAGACTACTAGACGAGTCAATAGCCAGCGAACAGGCC    6584
29B      GCCATTAGCGACAATCCTAACTACAGACTACTAGACGAGTCAATAGCCAGCGAACAGGCC    6660
e19      GCCATTAGCGACAATCCTAACTACAGACTACTAGACGAGTCAATAGCCAGCGAACAGGCC    6660
74e19    GCCATTAGCGACAATCCTAACTACAGACTACTAGACGAGTCAATAGCCAGCGAACAGGCC    6660
74       GCCATTAGCGACAATCCTAACTACAGACTACTAGACGAGTCAATAGCCAGCGAACAGGCC    6660
211      GCCATTAGCGACAATCCTAACTACAGACTACTAGACGAGTCAATAGCCAGCGAACAGGCC    6660
246      GCCATTAGCGACAATCCTAACTACAGACTACTAGACGAGTCAATAGCCAGCGAACAGGCC    6660
353      GCCATTAGCGACAATCCTAACTACAGACTACTAGACGAGTCAATAGCCAGCGAACAGGCC    6660
         ************************************************************
```

```
DAH      ACCATCCTAACGACTAGCAGCCCCAATCCCAACTACGAGATGATGCATCCACCAACCAGT   6644
29B      ACCATCCTAACGACTAGCAGCCCCAATCCCAACTACGAGATGATGCATCCACCAACCAGT   6720
e19      ACCATCCTAACGACTAGCAGCCCCAATCCCAACTACGAGATGATGCATCCACCAACCAGT   6720
74e19    ACCATCCTAACGACTAGCAGCCCCAATCCCAACTACGAGATGATGCATCCACCAACCAGT   6720
74       ACCATCCTAACGACTAGCAGCCCCAATCCCAACTACGAGATGATGCATCCACCAACCAGT   6720
211      ACCATCCTAACGACTAGCAGCCCCAATCCCAACTACGAGATGATGCATCCACCAACCAGT   6720
246      ACCATCCTAACGACTAGCAGCCCCAATCCCAACTACGAGATGATGCATCCACCAACCAGT   6720
353      ACCATCCTAACGACTAGCAGCCCCAATCCCAACTACGAGATGATGCATCCACCAACCAGT   6720
         ************************************************************

DAH      CTGGTCAGTACCAATCCGAACTATATGCCCATGAATGAGACTCCAGTGCAGATGGCGGGT   6704
29B      CTGGTCAGTACCAATCCGAACTATATGCCCATGAATGAGACTCCAGTGCAGATGGCGGGT   6780
e19      CTGGTCAGTACCAATCCGAACTATATGCCCATGAATGAGACTCCAGTGCAGATGGCGGGT   6780
74e19    CTGGTCAGTACCAATCCGAACTATATGCCCATGAATGAGACTCCAGTGCAGATGGCGGGT   6780
74       CTGGTCAGTACCAATCCGAACTATATGCCCATGAATGAGACTCCAGTGCAGATGGCGGGT   6780
211      CTGGTCAGTACCAATCCGAACTATATGCCCATGAATGAGACTCCAGTGCAGATGGCGGGT   6780
246      CTGGTCAGTACCAATCCGAACTATATGCCCATGAATGAGACTCCAGTGCAGATGGCGGGT   6780
353      CTGGTCAGTACCAATCCGAACTATATGCCCATGAATGAGACTCCAGTGCAGATGGCGGGT   6780
         ************************************************************

DAH      GTGACCATTAGCCATAATCCGAATTACCAGCCCATGCAGGCGCCGTTGAATGCACGCCAA   6764
29B      GTGACCATTAGCCATAATCCGAATTACCAGCCCATGCAGGCGCCGTTGAATGCACGCCAA   6840
e19      GTGACCATTAGCCATAATCCGAATTACCAGCCCATGCAGGCGCCGTTGAATGCACGCCAA   6840
74e19    GTGACCATTAGCCATAATCCGAATTACCAGCCCATGCAGGCGCCGTTGAATGCACGCCAA   6840
74       GTGACCATTAGCCATAATCCGAATTACCAGCCCATGCAGGCGCCGTTGAATGCACGCCAA   6840
211      GTGACCATTAGCCATAATCCGAATTACCAGCCCATGCAGGCGCCGTTGAATGCACGCCAA   6840
246      GTGACCATTAGCCATAATCCGAATTACCAGCCCATGCAGGCGCCGTTGAATGCACGCCAA   6840
353      GTGACCATTAGCCATAATCCGAATTACCAGCCCATGCAGGCGCCGTTGAATGCACGCCAA   6840
         ************************************************************

DAH      AGCCAAAGTAGCTCCGACGAGGACAACGAGCAGGAGGAGGACGATGAGGATGAGGACGAC   6824
29B      AGCCAAAGTAGCTCCGACGAGGACAACGAGCAGGAGGAGGACGATGAGGATGAGGACGAC   6900
e19      AGCCAAAGTAGCTCCGACGAGGACAACGAGCAGGAGGAGGACGATGAGGATGAGGACGAC   6900
74e19    AGCCAAAGTAGCTCCGACGAGGACAACGAGCAGGAGGAGGACGATGAGGATGAGGACGAC   6900
74       AGCCAAAGTAGCTCCGACGAGGACAACGAGCAGGAGGAGGACGATGAGGATGAGGACGAC   6900
211      AGCCAAAGTAGCTCCGACGAGGACAACGAGCAGGAGGAGGACGATGAGGATGAGGACGAC   6900
246      AGCCAAAGTAGCTCCGACGAGGACAACGAGCAGGAGGAGGACGATGAGGATGAGGACGAC   6900
353      AGCCAAAGTAGCTCCGACGAGGACAACGAGCAGGAGGAGGACGATGAGGATGAGGACGAC   6900
         ************************************************************

DAH      GACGTGGACGATGAGCATGTGGAGCACATCAAGATGGAGCGCATGCCATTGAGTCGGCCC   6884
29B      GACGTGGACGATGAGCATGTGGAGCACATCAAGATGGAGCGCATGCCATTGAGTCGGCCC   6960
e19      GACGTGGACGATGAGCATGTGGAGCACATCAAGATGGAGCGCATGCCATTGAGTCGGCCC   6960
74e19    GACGTGGACGATGAGCATGTGGAGCACATCAAGATGGAGCGCATGCCATTGAGTCGGCCC   6960
74       GACGTGGACGATGAGCATGTGGAGCACATCAAGATGGAGCGCATGCCATTGAGTCGGCCC   6960
211      GACGTGGACGATGAGCATGTGGAGCACATCAAGATGGAGCGCATGCCATTGAGTCGGCCC   6960
246      GACGTGGACGATGAGCATGTGGAGCACATCAAGATGGAGCGCATGCCATTGAGTCGGCCC   6960
353      GACGTGGACGATGAGCATGTGGAGCACATCAAGATGGAGCGCATGCCATTGAGTCGGCCC   6960
         ************************************************************

DAH      AGGCAAAGAGCGTTGCCCAGCAAGACGCAGCCGCCTCGCAGTCGCAGCGTCAGCCAAACG   6944
29B      AGGCAAAGAGCGTTGCCCAGCAAGACGCAGCCGCCTCGCAGTCGCAGCGTCAGCCAAACG   7020
e19      AGGCAAAGAGCGTTGCCCAGCAAGACGCAGCCGCCTCGCAGTCGCAGCGTCAGCCAAACG   7020
74e19    AGGCAAAGAGCGTTGCCCAGCAAGACGCAGCCGCCTCGCAGTCGCAGCGTCAGCCAAACG   7020
74       AGGCAAAGAGCGTTGCCCAGCAAGACGCAGCCGCCTCGCAGTCGCAGCGTCAGCCAAACG   7020
211      AGGCAAAGAGCGTTGCCCAGCAAGACGCAGCCGCCTCGCAGTCGCAGCGTCAGCCAAACG   7020
246      AGGCAAAGAGCGTTGCCCAGCAAGACGCAGCCGCCTCGCAGTCGCAGCGTCAGCCAAACG   7020
353      AGGCAAAGAGCGTTGCCCAGCAAGACGCAGCCGCCTCGCAGTCGCAGCGTCAGCCAAACG   7020
         ************************************************************
```

```
DAH      AGAAAATCTCCTACGAATCCCAACTCCGGAATCGGAGCGACAGGAGCCGGAAACCGATCC   7004
29B      AGAAAATCTCCTACGAATCCCAACTCCGGAATCGGAGCGACAGGAGCCGGAAACCGATCC   7080
e19      AGAAAATCTCCTACGAATCCCAACTCCGGAATCGGAGCGACAGGAGCCGGAAACCGATCC   7080
74e19    AGAAAATCTCCTACGAATCCCAACTCCGGAATCGGAGCGACAGGAGCCGGAAACCGATCC   7080
74       AGAAAATCTCCTACGAATCCCAACTCCGGAATCGGAGCGACAGGAGCCGGAAACCGATCC   7080
211      AGAAAATCTCCTACGAATCCCAACTCCGGAATCGGAGCGACAGGAGCCGGAAACCGATCC   7080
246      AGAAAATCTCCTACGAATCCCAACTCCGGAATCGGAGCGACAGGAGCCGGAAACCGATCC   7080
353      AGAAAATCTCCTACGAATCCCAACTCCGGAATCGGAGCGACAGGAGCCGGAAACCGATCC   7080
         ************************************************************

DAH      AACTTGCTTAAAGAGAACTGGCTGCGACCGGCGAGTACGCCAAGGCCTCCACCACCCAAT   7064
29B      AACTTGCTTAAAGAGAACTGGCTGCGACCGGCGAGTACGCCAAGGCCTCCACCACCCAAT   7140
e19      AACTTGCTTAAAGAGAACTGGCTGCGACCGGCGAGTACGCCAAGGCCTCCACCACCCAAT   7140
74e19    AACTTGCTTAAAGAGAACTGGCTGCGACCGGCGAGTACGCCAAGGCCTCCACCACCCAAT   7140
74       AACTTGCTTAAAGAGAACTGGCTGCGACCGGCGAGTACGCCAAGGCCTCCACCACCCAAT   7140
211      AACTTGCTTAAAGAGAACTGGCTGCGACCGGCGAGTACGCCAAGGCCTCCACCACCCAAT   7140
246      AACTTGCTTAAAGAGAACTGGCTGCGACCGGCGAGTACGCCAAGGCCTCCACCACCCAAT   7140
353      AACTTGCTTAAAGAGAACTGGCTGCGACCGGCGAGTACGCCAAGGCCTCCACCACCCAAT   7140
         ************************************************************

DAH      GGATTCATCGGAAGGGAGGCGtaatcgttacgaactgtagttctgtagaaaaacatgtaa   7124
29B      GGATTCATCGGAAGGGAGGCGtaatcgttacgaactgtagttctgtagaaaaacatgtaa   7200
e19      GGATTCATCGGAAGGGAGGCGtaatcgttacgaactgtagttctgtagaaaaacatgtaa   7200
74e19    GGATTCATCGGAAGGGAGGCGtaatcgttacgaactgtagttctgtagaaaaacatgtaa   7200
74       GGATTCATCGGAAGGGAGGCGtaatcgttacgaactgtagttctgtagaaaaacatgtaa   7200
211      GGATTCATCGGAAGGGAGGCGtaatcgttacgaactgtagttctgtagaaaaacatgtaa   7200
246      GGATTCATCGGAAGGGAGGCGtaatcgttacgaactgtagttctgtagaaaaacatgtaa   7200
353      GGATTCATCGGAAGGGAGGCGtaatcgttacgaactgtagttctgtagaaaaacatgtaa   7200
         ************************************************************

DAH      atagatgaggagaagcagcatggatatagatagagtagcaagtctttgccgcaagccaag   7184
29B      atagatgaggagaagcagcatggatatagatagagtagcaagtctttgccgcaagccaag   7260
e19      atagatgaggagaagcagcatggatatagatagagtagcaagtctttgccgcaagccaag   7260
74e19    atagatgaggagaagcagcatggatatagatagagtagcaagtctttgccgcaagccaag   7260
74       atagatgaggagaagcagcatggatatagatagagtagcaagtctttgccgcaagccaag   7260
211      atagatgaggagaagcagcatggatatagatagagtagcaagtctttgccgcaagccaag   7260
246      atagatgaggagaagcagcatggatatagatagagtagcaagtctttgccgcaagccaag   7260
353      atagatgaggagaagcagcatggatatagatagagtagcaagtctttgccgcaagccaag   7260
         ************************************************************

DAH      cttcttggaatttgactcttctctatgcctgcaaattagggtggaggagatgtaggtcca   7244
29B      cttcttggaatttgactcttctctatgcctgcaaattagggtggaggagatgtaggtcca   7320
e19      cttcttggaatttgactcttctctatgcctgcaaattagggtggaggagatgtaggtcca   7320
74e19    cttcttggaatttgactcttctctatgcctgcaaattagggtggaggagatgtaggtcca   7320
74       cttcttggaatttgactcttctctatgcctgcaaattagggtggaggagatgtaggtcca   7320
211      cttcttggaatttgactcttctctatgcctgcaaattagggtggaggagatgtaggtcca   7320
246      cttcttggaatttgactcttctctatgcctgcaaattagggtggaggagatgtaggtcca   7320
353      cttcttggaatttgactcttctctatgcctgcaaattagggtggaggagatgtaggtcca   7320
         ************************************************************

DAH      aagatatttgtatattttcaatatgactggctagcaatccattgtaacatttctagcgat   7304
29B      aagatatttgtatattttcaatatgactggctagcaatccattgtaacatttctagcgat   7380
e19      aagatatttgtatattttcaatatgactggctagcaatccattgtaacatttctagcgat   7380
74e19    aagatatttgtatattttcaatatgactggctagcaatccattgtaacatttctagcgat   7380
74       aagatatttgtatattttcaatatgactggctagcaatccattgtaacatttctagcgat   7380
211      aagatatttgtatattttcaatatgactggctagcaatccattgtaacatttctagcgat   7380
246      aagatatttgtatattttcaatatgactggctagcaatccattgtaacatttctagcgat   7380
353      aagatatttgtatattttcaatatgactggctagcaatccattgtaacatttctagcgat   7380
         ************************************************************
```

```
DAH      gaagcttcaaacaagaagttgattctttagttaaaattttgcattaatagaaaattaagc   7364
29B      gaagcttcaaacaagaagttgattctttagttaaaattttgcattaatagaaaattaagc   7440
e19      gaagcttcaaacaagaagttgattctttagttaaaattttgcattaatagaaaattaagc   7440
74e19    gaagcttcaaacaagaagttgattctttagttaaaattttgcattaatagaaaattaagc   7440
74       gaagcttcaaacaagaagttgattctttagttaaaattttgcattaatagaaaattaagc   7440
211      gaagcttcaaacaagaagttgattctttagttaaaattttgcattaatagaaaattaagc   7440
246      gaagcttcaaacaagaagttgattctttagttaaaattttgcattaatagaaaattaagc   7440
353      gaagcttcaaacaagaagttgattctttagttaaaattttgcattaatagaaaattaagc   7440
         ************************************************************

DAH      acattgtgtcagtcttcgtaaaataacacacctttccgtagatggatgacaccttttctt   7424
29B      acattgtgtcagtcttcgtaaaataacacacctttccgtagatggatgacaccttttctt   7500
e19      acattgtgtcagtcttcgtaaaataacacacctttccgtagatggatgacaccttttctt   7500
74e19    acattgtgtcagtcttcgtaaaataacacacctttccgtagatggatgacaccttttctt   7500
74       acattgtgtcagtcttcgtaaaataacacacctttccgtagatggatgacaccttttctt   7500
211      acattgtgtcagtcttcgtaaaataacacacctttccgtagatggatgacaccttttctt   7500
246      acattgtgtcagtcttcgtaaaataacacacctttccgtagatggatgacaccttttctt   7500
353      acattgtgtcagtcttcgtaaaataacacacctttccgtagatggatgacaccttttctt   7500
         ************************************************************

DAH      cttggtgtcgaaccaaaattgtaaattttagattcatagagcctaattttcaattctcaa   7484
29B      cttggtgtcgaaccaaaattgtaaattttagattcatagagcctaattttcaattctcaa   7560
e19      cttggtgtcgaaccaaaattgtaaattttagattcatagagcctaattttcaattctcaa   7560
74e19    cttggtgtcgaaccaaaattgtaaattttagattcatagagcctaattttcaattctcaa   7560
74       cttggtgtcgaaccaaaattgtaaattttagattcatagagcctaattttcaattctcaa   7560
211      cttggtgtcgaaccaaaattgtaaattttagattcatagagcctaattttcaattctcaa   7560
246      cttggtgtcgaaccaaaattgtaaattttagattcatagagcctaattttcaattctcaa   7560
353      cttggtgtcgaaccaaaattgtaaattttagattcatagagcctaattttcaattctcaa   7560
         ************************************************************

DAH      ttgttcgacgattaggcaaacgactatatggagtcgaagatatggaaacaaaggcttaat   7544
29B      ttgttcgacgattaggcaaacgactatatggagtcgaagatatggaaacaaaggcttaat   7620
e19      ttgttcgacgattaggcaaacgactatatggagtcgaagatatggaaacaaaggcttaat   7620
74e19    ttgttcgacgattaggcaaacgactatatggagtcgaagatatggaaacaaaggcttaat   7620
74       ttgttcgacgattaggcaaacgactatatggagtcgaagatatggaaacaaaggcttaat   7620
211      ttgttcgacgattaggcaaacgactatatggagtcgaagatatggaaacaaaggcttaat   7620
246      ttgttcgacgattaggcaaacgactatatggagtcgaagatatggaaacaaaggcttaat   7620
353      ttgttcgacgattaggcaaacgactatatggagtcgaagatatggaaacaaaggcttaat   7620
         ************************************************************

DAH      ttgctaagaacgagccaaaaagattgctaaccgccttcacgattctttgttcaagccatc   7604
29B      ttgctaagaacgagccaaaaagattgctaaccgccttcacgattctttgttcaagccatc   7680
e19      ttgctaagaacgagccaaaaagattgctaaccgccttcacgattctttgttcaagccatc   7680
74e19    ttgctaagaacgagccaaaaagattgctaaccgccttcacgattctttgttcaagccatc   7680
74       ttgctaagaacgagccaaaaagattgctaaccgccttcacgattctttgttcaagccatc   7680
211      ttgctaagaacgagccaaaaagattgctaaccgccttcacgattctttgttcaagccatc   7680
246      ttgctaagaacgagccaaaaagattgctaaccgccttcacgattctttgttcaagccatc   7680
353      ttgctaagaacgagccaaaaagattgctaaccgccttcacgattctttgttcaagccatc   7680
         ************************************************************

DAH      gaaccaccctaaccgcgcatgtataataccaatagtttcgattaagcctatttatgttta   7664
29B      gaaccaccctaaccgcgcatgtataataccaatagtttcgattaagcctatttatgttta   7740
e19      gaaccaccctaaccgcgcatgtataataccaatagtttcgattaagcctatttatgttta   7740
74e19    gaaccaccctaaccgcgcatgtataataccaatagtttcgattaagcctatttatgttta   7740
74       gaaccaccctaaccgcgcatgtataataccaatagtttcgattaagcctatttatgttta   7740
211      gaaccaccctaaccgcgcatgtataataccaatagtttcgattaagcctatttatgttta   7740
246      gaaccaccctaaccgcgcatgtataataccaatagtttcgattaagcctatttatgttta   7740
353      gaaccaccctaaccgcgcatgtataataccaatagtttcgattaagcctatttatgttta   7740
         ************************************************************
```

```
        71
DAH         cgtatt      7670
29B         cgtatt      7746
e19         cgtatt      7746
74e19       cgtatt      7746
74          cgtatt      7746
211         cgtatt      7746
246         cgtatt      7746
353         cgtatt      7746
            ******
```

**Fig S4. Master PROT alignment HR alleles with Wildtypes *w*DAH and 29B(HR)**

```
DAH        MFNMPRGVTKSKSKRGKIKMENDMAAAATTTACTLGHICVLCRQEMLLDTCCCRQAVEAV    60
29B        MFNMPRGVTKSKSKRGKIKMENDMAAAATTTACTLGHICVLCRQEMLLDTCCCRQAVEAV    60
e19        MFNMPRGVTKSKSKRGKIKMENDMAAAATTTACTLGHICVLCRQEMLLDTCCCRQAVEAV    60
74e19      MFNMPRGVTKSKSKRGKIKMENDMAAAATTTACTLGHICVLCRQEMLLDTCCCRQAVEAV    60
74         MFNMPRGVTKSKSKRGKIKMENDMAAAATTTACTLGHICVLCRQEMLLDTCCCRQAVEAV    60
211        MFNMPRGVTKSKSKRGKIKMENDMAAAATTTACTLGHICVLCRQEMLLDTCCCRQAVEAV    60
246        MFNMPRGVTKSKSKRGKIKMENDMAAAATTTACTLGHICVLCRQEMLLDTCCCRQAVEAV    60
353        MFNMPRGVTKSKSKRGKIKMENDMAAAATTTACTLGHICVLCRQEMLLDTCCCRQAVEAV    60
           ************************************************************

DAH        DSPASSEEAYSSSNSSSCQASSEISAEEVWFLSHDDIVLCRRPKFDEVETTGKKRDVKCS    120
29B        DSPASSEEAYSSSNSSSCQASSEISAEEVWFLSHDDIVLCRRPKFDEVETTGKKRDVKCS    120
e19        DSPASSEEAYSSSNSSSCQASSEISAEEVWFLSHDDIVLCRRPKFDEVETTGKKRDVKCS    120
74e19      DSPASSEEAYSSSNSSSCQASSEISAEEVWFLSHDDIVLCRRPKFDEVETTGKKRDVKCS    120
74         DSPASSEEAYSSSNSSSCQASSEISAEEVWFLSHDDIVLCRRPKFDEVETTGKKRDVKCS    120
211        DSPASSEEAYSSSNSSSCQASSEISAEEVWFLSHDDIVLCRRPKFDEVETTGKKRDVKCS    120
246        DSPASSEEAYSSSNSSSCQASSEISAEEVWFLSHDDIVLCRRPKFDEVETTGKKRDVKCS    120
353        DSPASSEEAYSSSNSSSCQASSEISAEEVWFLSHDDIVLCRRPKFDEVETTGKKRDVKCS    120
           ************************************************************

DAH        GHQCSNECDDGSTKNNRQQRENFNIFSNCHNILRTLQSLLLLMFNCGIFNKRRRRQHQQQ    180
29B        GHQCSNECDDGSTKNNRQQRENFNIFSNCHNILRTLQSLLLLMFNCGIFNKRRRRQHQQQ    180
e19        GHQCSNECDDGSTKNNRQQRENFNIFSNCHNILRTLQSLLLLMFNCGIFNKRRRRQHQQQ    180
74e19      GHQCSNECDDGSTKNNRQQRENFNIFSNCHNILRTLQSLLLLMFNCGIFNKRRRRQHQQQ    180
74         GHQCSNECDDGSTKNNRQQRENFNIFSNCHNILRTLQSLLLLMFNCGIFNKRRRRQHQQQ    180
211        GHQCSNECDDGSTKNNRQQRENFNIFSNCHNILRTLQSLLLLMFNCGIFNKRRRRQHQQQ    180
246        GHQCSNECDDGSTKNNRQQRENFNIFSNCHNILRTLQSLLLLMFNCGIFNKRRRRQHQQQ    180
353        GHQCSNECDDGSTKNNRQQRENFNIFSNCHNILRTLQSLLLLMFNCGIFNKRRRRQHQQQ    180
           ************************************************************

DAH        HHHHYQHHQHHHQQHLQRQQANVSYTKFLLLLQTLAAATTRLSLSPKNYKQQQQLQHNQQ    240
29B        HHHHYQHHQHHHQQHLQRQQANVSYTKFLLLLQTLAAATTRLSLSPKNYKQQQQLQHNQQ    240
e19        HHHHYQHHQHHHQQHLQRQQANVSYTKFLLLLQTLAAATTRLSLSPKNYKQQQQLQHNQQ    240
74e19      HHHHYQHHQHHHQQHLQRQQANVSYTKFLLLLQTLAAATTRLSLSPKNYKQQQQLQHNQQ    240
74         HHHHYQHHQHHHQQHLQRQQANVSYTKFLLLLQTLAAATTRLSLSPKNYKQQQQLQHNQQ    240
211        HHHHYQHHQHHHQQHLQRQQANVSYTKFLLLLQTLAAATTRLSLSPKNYKQQQQLQHNQQ    240
246        HHHHYQHHQHHHQQHLQRQQANVSYTKFLLLLQTLAAATTRLSLSPKNYKQQQQLQHNQQ    240
353        HHHHYQHHQHHHQQHLQRQQANVSYTKFLLLLQTLAAATTRLSLSPKNYKQQQQLQHNQQ    240
           ************************************************************

DAH        LPRATPQQKQQEKDRHKCFHYKHNYSYSPGISLLLFILLANTLAIQAVVLPAHQQHLLHN    300
29B        LPRATPQQKQQEKDRHKCFHYKHNYSYSPGISLLLFILLANTLAIQAVVLPAHQQHLLHN    300
e19        LPRATPQQKQQEKDRHKCFHYKHNYSYSPGISLLLFILLANTLAIQAVVLPAHQQHLLHN    300
74e19      LPRATPQQKQQEKDRHKCFHYKHNYSYSPGISLLLFILLANTLAIQAVVLPAHQQHLLHN    300
74         LPRATPQQKQQEKDRHKCFHYKHNYSYSPGISLLLFILLANTLAIQAVVLPAHQQHLLHN    300
211        LPRATPQQKQQEKDRHKCFHYKHNYSYSPGISLLLFILLANTLAIQAVVLPAHQQHLLHN    300
246        LPRATPQQKQQEKDRHKCFHYKHNYSYSPGISLLLFILLANTLAIQAVVLPAHQQHLLHN    300
353        LPRATPQQKQQEKDRHKCFHYKHNYSYSPGISLLLFILLANTLAIQAVVLPAHQQHLLHN    300
           ************************************************************
```

```
DAH       DIADGLDKTALSVSGTQSRWTRSESNPTMRLSQNVKPCKSMDIRNMVSHFNQLENCTVIE     360
29B       DIADGLDKTALSVSGTQSRWTRSESNPTMRLSQNVKPCKSMDIRNMVSHFNQLENCTVIE     360
e19       DIADGLDKTALSVSGTQSRWTRSESNPTMRLSQNVKPCKSMDIRNMVSHFNQLENCTVIE     360
74e19     DIADGLDKTALSVSGTQSRWTRSESNPTMRLSQNVKPCKSMDIRNMVSHFNQLENCTVIE     360
74        DIADGLDKTALSVSGTQSRWTRSESNPTMRLSQNVKPCKSMDIRNMVSHFNQLENCTVIE     360
211       DIADGLDKTALSVSGTQSRWTRSESNPTMRLSQNVKPCKSMDIRNMVSHFNQLENCTVIE     360
246       DIADGLDKTALSVSGTQSRWTRSESNPTMRLSQNVKPCKSMDIRNMVSHFNQLENCTVIE     360
353       DIADGLDKTALSVSGTQSRWTRSESNPTMRLSQNVKPCKSMDIRNMVSHFNQLENCTVIE     360
          ************************************************************

DAH       GFLLIDLINDASPLNRSFPKLTEVTDYIIIYRVTGLHSLSKIFPNLSVIRGNKLFDGYAL     420
29B       GFLLIDLINDASPLNRSFPKLTEVTDYIIIYRVTGLHSLSKIFPNLSVIRGNKLFDGYAL     420
e19       GFLLIDLINDASPLNRSFPKLTEVTDYIIIYRVTGLHSLSKIFPNLSVIRGNKLFDGYAL     420
74e19     GFLLIDLINDASPLNRSFPKLTEVTDYIIIYRVTGLHSLSKIFPNLSVIRGNKLFDGYAL     420
74        GFLLIDLINDASPLNRSFPKLTEVTDYIIIYRVTGLHSLSKIFPNLSVIRGNKLFDGYAL     420
211       GFLLIDLINDASPLNRSFPKLTEVTDYIIIYRVTGLHSLSKIFPNLSVIRGNKLFDGYAL     420
246       GFLLIDLINDASPLNRSFPKLTEVTDYIIIYRVTGLHSLSKIFPNLSVIRGNKLFDGYAL     420
353       GFLLIDLINDASPLNRSFPKLTEVTDYIIIYRVTGLHSLSKIFPNLSVIRGNKLFDGYAL     420
          ************************************************************

DAH       VVYSNFDLMDLGLHKLRSITRGGVRIEKNHKLCYDRTIDWLEILAENETQLVVLTENGKE     480
29B       VVYSNFDLMDLGLHKLRSITRGGVRIEKNHKLCYDRTIDWLEILAENETQLVVLTENGKE     480
e19       VVYSNFDLMDLGLHKLRSITRGGVRIEKNHKLCYDRTIDWLEILAENETQLVVLTENGKE     480
74e19     VVYSNFDLMDLGLHKLRSITRGGVRIEKNHKLCYDRTIDWLEILAENETQLVVLTENGKE     480
74        VVYSNFDLMDLGLHKLRSITRGGVRIEKNHKLCYDRTIDWLEILAENETQLVVLTENGKE     480
211       VVYSNFDLMDLGLHKLRSITRGGVRIEKNHKLCYDRTIDWLEILAENETQLVVLTENGKE     480
246       VVYSNFDLMDLGLHKLRSITRGGVRIEKNHKLCYDRTIDWLEILAENETQLVVLTENGKE     480
353       VVYSNFDLMDLGLHKLRSITRGGVRIEKNHKLCYDRTIDWLEILAENETQLVVLTENGKE     480
          ************************************************************

DAH       KECRLSKCPGEIRIEEGHDTTAIEGELNASCQLHNNRRLCWNSKLCQTKCPEKCRNNCID     540
29B       KECRLSKCPGEIRIEEGHDTTAIEGELNASCQLHNNRRLCWNSKLCQTKCPEKCRNNCID     540
e19       KECRLSKCPGEIRIEEGHDTTAIEGELNASCQLHNNRRLCWNSKLCQTKCPEKCRNNCID     540
74e19     KECRLSKCPGEIRIEEGHDTTAIEGELNASCQLHNNRRLCWNSKLCQTKCPEKCRNNCID     540
74        KECRLSKCPGEIRIEEGHDTTAIEGELNASCQLHNNRRLCWNSKLCQTKCPEKCRNNCID     540
211       KECRLSKCPGEIRIEEGHDTTAIEGELNASCQLHNNRRLCWNSKLCQTKCPEKCRNNCID     540
246       KECRLSKCPGEIRIEEGHDTTAIEGELNASCQLHNNRRLCWNSKLCQTKCPEKCRNNCID     540
353       KECRLSKCPGEIRIEEGHDTTAIEGELNASCQLHNNRRLCWNSKLCQTKCPEKCRNNCID     540
          ************************************************************

DAH       EHTCCSQDCLGGCVIDKNGNESCISCRNVSFNNICMDSCPKGYYQFDSRCVTANECITLT     600
29B       EHTCCSQDCLGGCVIDKNGNESCISCRNVSFNNICMDSCPKGYYQFDSRCVTANECITLT     600
e19       EHTCCSQDCLGGCVIDKNGNESCISCRNVSFNNICMDSCPKGYYQFDSRCVTANECITLT     600
74e19     EHTCCSQDCLGGCVIDKNGNESCISCRNVSFNNICMDSCPKGYYQFDSRCVTANECITLT     600
74        EHTCCSQDCLGGCVIDKNGNESCISCRNVSFNNICMDSCPKGYYQFDSRCVTANECITLT     600
211       EHTCCSQDCLGGCVIDKNGNESCISCRNVSFNNICMDSCPKGYYQFDSRCVTANECITLT     600
246       EHTCCSQDCLGGCVIDKNGNESCISCRNVSFNNICMDSCPKGYYQFDSRCVTANECITLT     600
353       EHTCCSQDCLGGCVIDKNGNESCISCRNVSFNNICMDSCPKGYYQFDSRCVTANECITLT     600
          ************************************************************

DAH       KFETNSVYSGIPYNGQCITHCPTGYQKSENKRMCEPCPGGKCDKECSSGLIDSLERAREF     660
29B       KFETNSVYSGIPYNGQCITHCPTGYQKSENKRMCEPCPGGKCDKECSSGLIDSLERAREF     660
e19       KFETNSVYSGIPYNGQCITHCPTGYQKSENKRMCEPCPGGKCDKECSSGLIDSLERAREF     660
74e19     KFETNSVYSGIPYNGQCITHCPTGYQKSENKRMCEPCPGGKCDKECSSGLIDSLERAREF     660
74        KFETNSVYSGIPYNGQCITHCPTGYQKSENKRMCEPCPGGKCDKECSSGLIDSLERAREF     660
211       KFETNSVYSGIPYNGQCITHCPTGYQKSENKRMCEPCPGGKCDKECSSGLIDSLERAREF     660
246       KFETNSVYSGIPYNGQCITHCPTGYQKSENKRMCEPCPGGKCDKECSSGLIDSLERAREF     660
353       KFETNSVYSGIPYNGQCITHCPTGYQKSENKRMCEPCPGGKCDKECSSGLIDSLERAREF     660
          ************************************************************
```

74

```
DAH      HGCTIITGTEPLTISIKRESGAHVMDELKYGLAAVHKIQSSLMVHLTYGLKSLKFFQSLT    720
29B      HGCTIITGTEPLTISIKRESGAHVMDELKYGLAAVHKIQSSLMVHLTYGLKSLKFFQSLT    720
e19      HGCTIITGTEPLTISIKRESGAHVMDELKYGLAAVHKIQSSLMVHLTYGLKSLKFFQSLT    720
74e19    HGCTIITGTEPLTISIKRESGAHVMDELKYGLAAVHKIQSSLMVHLTYGLKSLKFFQSLT    720
74       HGCTIITGTEPLTISIKRESGAHVMDELKYGLAAVHKIQSSLMVHLTYGLKSLKFFQSLT    720
211      HGCTIITGTEPLTISIKRESGAHVMDELKYGLAAVHKIQSSLMVHLTYGLKSLKFFQSLT    720
246      HGCTIITGTEPLTISIKRESGAHVMDELKYGLAAVHKIQSSLMVHLTYGLKSLKFFQSLT    720
353      HGCTIITGTEPLTISIKRESGAHVMDELKYGLAAVHKIQSSLMVHLTYGLKSLKFFQSLT    720
         ************************************************************

DAH      EISGDPPMDADKYALYVLDNRDLDELWGPNQTVFIRKGGVFFHFNPKLCVSTINQLLPML    780
29B      EISGDPPMDADKYALYVLDNRDLDELWGPNQTVFIRKGGVFFHFNPKLCVSTINQLLPML    780
e19      EISGDPPMDADKYALYVLDNRDLDELWGPNQTVFIRKGGVFFHFNPKLCVSTINQLLPML    780
74e19    EISGDPPMDADKYALYVLDNRDLDELWGPNQTVFIRKGGVFFHFNPKLCVSTINQLLPML    780
74       EISGDPPMDADKYALYVLDNRDLDELWGPNQTVFIRKGGVFFHFNPKLCVSTINQLLPML    780
211      EISGDPPMDADKYALYVLDNRDLDELWGPNQTVFIRKGGVFFHFNPKLCVSTINQLLPML    780
246      EISGDPPMDADKYALYVLDNRDLDELWGPNQTVFIRKGGVFFHFNPKLCVSTINQLLPML    780
353      EISGDPPMDADKYALYVLDNRDLDELWGPNQTVFIRKGGVFFHFNPKLCVSTINQLLPML    780
         ************************************************************

DAH      ASKPKFFEKSDVGADSNGNRGSCGTAVLNVTLQSVGANSAMLNVTTKVEIGEPQKPSNAT    840
29B      ASKPKFFEKSDVGADSNGNRGSCGTAVLNVTLQSVGANSAMLNVTTKVEIGEPQKPSNAT    840
e19      ASKPKFFEKSDVGADSNGNRGSCGTAVLNDTLQSVGANSAMLNVTTKVEIGEPQKPSNAT    840
74e19    ASKPKFFEKSDVGADSNGNRGSCGTAVLNDTLQSVGANSAMLNVTTKVEIGEPQKPSNAT    840
74       ASKPKFFEKSDVGADSNGNRGSCGTAVLNVTLQSVGANSAMLNVTTKVEIGEPQKPSNAT    840
211      ASKPKFFEKSDVGADSNGNRGSCGTAVLNVTLQSVGANSAMLNVTTKVEIGEPQKPSNAT    840
246      ASKPKFFEKSDVGADSNGNRGSCGTAVLNVTLQSVGANSAMLNVTTKVEIGEPQKPSNAT    840
353      ASKPKFFEKSDVGADSNGNRGSCGTAVLNVTLQSVGANSAMLNVTTKVEIGEPQKPSNAT    840
         ***************************** ******************************

DAH      IVFKDPRAFIGFVFYHMIDPYGNSTKSSDDPCDDRWKVSSPEKSGVMVLSNLIPYTNYSY    900
29B      IVFKDPRAFIGFVFYHMIDPYGNSTKSSDDPCDDRWKVSSPEKSGVMVLSNLIPYTNYSY    900
e19      IVFKDPRAFIGFVFYHMIDPYGNSTKSSDDPCDDRWKVSSPEKSGVMVLSNLIPYTNYSY    900
74e19    IVFKDPRAFIGFVFYHMIDPYGNSTKSSDDPCDDRWKVSSPEKSGVMVLSNLIPYTNYSY    900
74       IVFKDPRAFIGFVFYHMIDPYGNSTKSSDDPCDDRWKVSSPEKSGVMVLSNLIPYTNYSY    900
211      IVFKDPRAFIGFVFYHMIDPYGNSTKSSDDPCDDRWKVSSPEKSGVMVLSNLIPYTNYSY    900
246      IVFKDPRAFIGFVFYHMIDPYGNSTKSSDDPCDDRWKVSSPEKSGVMVLSNLIPYTNYSY    900
353      IVFKDPRAFIGFVFYHMIDPYGNSTKSSDDPCDDRWKVSSPEKSGVMVLSNLIPYTNYSY    900
         ************************************************************

DAH      YVRTMAISSELTNAESDVKNFRTNPGRPSKVTEVVATAISDSKINVTWSYLDKPYGVLTR    960
29B      YVRTMAISSELTNAESDVKNFRTNPGRPSKVTEVVATAISDSKINVTWSYLDKPYGVLTR    960
e19      YVRTMAISSELTNAESDVKNFRTNPGRPSKVTEVVATAISDSKINVTWSYLDKPYGVLTR    960
74e19    YVRTMAISSELTNAESDVKNFRTNPGRPSKVTEVVATAISDSKINVTWSYLDKPYGVLTR    960
74       YVRTMAISSELTNAESDVKNFRTNPGRPSKVTEVVATAISDSKINVTWSYLDKPYGVLTR    960
211      YVRTMAISSELTNAESDVKNFRTNPGRPSKVTEVVATAISDSKINVTWSYLDKPYGVLTR    960
246      YVRTMAISSELTNAESDVKNFRTNPGRPSKVTEVVATAISDSKINVTWSYLDKPYGVLTR    960
353      YVRTMAISSELTNAESDVKNFRTNPGRPSKVTEVVATAISDSKINVTWSYLDKPYGVLTR    960
         ************************************************************

DAH      YFIKAKLINRPTRNNNRDYCTEPLVKAMENDLPATTPTKKISDPLAGDCKCVEGSKKTSS    1020
29B      YFIKAKLINRPTRNNNRDYCTEPLVKAMENDLPATTPTKKISDPLAGDCKCVEGSKKTSS    1020
e19      YFIKAKLINRPTRNNNRDYCTEPLVKAMENDLPATTPTKKISDPLAGDCKCVEGSKKTSS    1020
74e19    YFIKAKLINRPTRNNNRDYCTEPLVKAMENDLPATTPTKKISDPLAGDCKCVEGSKKTSS    1020
74       YFIKAKLINRPTRNNNRDYCTEPLVKAMENDLPATTPTKKISDPLAGDCKCVEGSKKTSS    1020
211      YFIKAKLINRPTRNNNRDYCTEPLVKAMENDLPATTPTKKISDPLAGDCKCVEGSKKTSS    1020
246      YFIKAKLINRPTRNNNRDYCTEPLVKAMENDLPATTPTKKISDPLAGDCKCVEGSKKTSS    1020
353      YFIKAKLINRPTRNNNRDYCTEPLVKAMENDLPATTPTKKISDPLAGDCKCVEGSKKTSS    1020
         ************************************************************
```

```
DAH     QEYDDRKVQAGMEFENALQNFIFVPNIRKSKNGSSDKSDGAEGAALDSNAIPNGGATNPS   1080
29B     QEYDDRKVQAGMEFENALQNFIFVPNIRKSKNGSSDKSDGAEGAALDSNAIPNGGATNPS   1080
e19     QEYDDRKVQAGMEFENALQNFIFVPNIRKSKNGSSDKSDGAEGAALDSNAIPNGGATNPS   1080
74e19   QEYDDRKVQAGMEFENALQNFIFVPNIRKSKNGSSDKSDGAEGAALDSNAIPNGGATNPS   1080
74      QEYDDRKVQAGMEFENALQNFIFVPNIRKSKNGSSDKSDGAEGAALDSNAIPNGGATNPS   1080
211     QEYDDRKVQAGMEFENALQNFIFVPNIRKSKNGSSDKSDGAEGAALDSNAIPNGGATNPS   1080
246     QEYDDRKVQAGMEFENALQNFIFVPNIRKSKNGSSDKSDGAEGAALDSNAIPNGGATNPS   1080
353     QEYDDRKVQAGMEFENALQNFIFVPNIRKSKNGSSDKSDGAEGAALDSNAIPNGGATNPS   1080
        ************************************************************

DAH     RRRRDVALEPELDDVEGSVLLRHVRSITDDTDAFFEKDDENTYKDEEDLSSNKQFYEVFA   1140
29B     RRRRDVALEPELDDVEGSVLLRHVRSITDDTDAFFEKDDENTYKDEEDLSSNKQFYEVFA   1140
e19     RRRRDVALEPELDDVEGSVLLRHVRSITDDTDAFFEKDDENTYKDEEDLSSNKQFYEVFA   1140
74e19   RRRRDVALEPELDDVEGSVLLRHVRSITDDTDAFFEKDDENTYKDEEDLSSNKQFYEVFA   1140
74      RRRRDVALEPELDDVEGSVLLRHVRSITDDTDAFFEKDDENTYKDEEDLSSNKQFYEVFA   1140
211     RRRRDVALEPELDDVEGSVLLRHVRSITDDTDAFFEKDDENTYKDEEDLSSNKQFYEVFA   1140
246     RRRRDVALEPELDDVEGSVLLRHVRSITDDTDAFFEKDDENTYKDEEDLSSNKQFYEVFA   1140
353     RRRRDVALEPELDDVEGSVLLRHVRSITDDTDAFFEKDDENTYKDEEDLSSNKQFYEVFA   1140
        ************************************************************

DAH     KELPPNQTHFVFEKLRHFTRYAIFVVACREEIPSEKLRDTSFKKSLCSDYDTVFQTTKRK   1200
29B     KELPPNQTHFVFEKLRHFTRYAIFVVACREEIPSEKLRDTSFKKSLCSDYDTVFQTTKRK   1200
e19     KELPPNQTHFVFEKLRHFTRYAIFVVACREEIPSEKLRDTSFKKSLCSDYDTVFQTTKRK   1200
74e19   KELPPNQTHFVFEKLRHFTRYAIFVVACREEIPSEKLRDTSFKKSLCSDYDTVFQTTKRK   1200
74      KELPPNQTHFVFEKLRHFTRYAIFVVACREEIPSEKLRDTSFKKSLCSDYDTVFQTTKRK   1200
211     KELPPNQTHFVFEKLRHFTRYAIFVVACREEIPSEKLRDTSFKKSLCSDYDTVFQTTKRK   1200
246     KELPPNQTHFVFEKLRHFTRYAIFVVACREEIPSEKLRDTSFKKSLCSDYDTVFQTTKRK   1200
353     KELPPNQTHFVFEKLRHFTRYAIFVVACREEIPSEKLRDTSFKKSLCSDYDTVFQTTKRK   1200
        ************************************************************

DAH     KFADIVMDLKVDLEHANNTESPVRVRWTPPVDPNGEIVTYEVAYKLQKPDQVEEKKCIPA   1260
29B     KFADIVMDLKVDLEHANNTESPVRVRWTPPVDPNGEIVTYEVAYKLQKPDQVEEKKCIPA   1260
e19     KFADIVMDLKVDLEHANNTESPVRVRWTPPVDPNGEIVTYEVAYKLQKPDQVEEKKCIPA   1260
74e19   KFADIVMDLKVDLEHANNTESPVRVRWTPPVDPNGEIVTYEVAYKLQKPDQVEEKKCIPA   1260
74      KFADIVMDLKVDLEHANNTESPVRVRWTPPVDPNGEIVTYEVAYKLQKPDQVEEKKCIPA   1260
211     KFADIVMDLKVDLEHANNTESPVRVRWTPPVDPNGEIVTYEVAYKLQKPDQVEEKKCIPA   1260
246     KFADIVMDLKVDLEHANNTESPVRVRWTPPVDPNGEIVTYEVAYKLQKPDQVEEKKCIPA   1260
353     KFADIVMDLKVDLEHANNTESPVRVRWTPPVDPNGEIVTYEVAYKLQKPDQVEEKKCIPA   1260
        ************************************************************

DAH     ADFNQTAGYLIKLNEGLYSFRVRANSIAGYGDFTEVEHIKVEPPPSYAKVFFWLLGIGLA   1320
29B     ADFNQTAGYLIKLNEGLYSFRVRANSIAGYGDFTEVEHIKVEPPPSYAKVFFWLLGIGLA   1320
e19     ADFNQTAGYLIKLNEGLYSFRVRANSIAGYGDFTEVEHIKVEPPPSYAKVFFWLLGIGLA   1320
74e19   ADFNQTAGYLIKLNEGLYSFRVRANSIAGYGDFTEVEHIKVEPPPSYAKVFFWLLGIGLA   1320
74      ADFNQTAGYLIKLNEGLYSFRVRANSIAGYGDFTEVEHIKVEPPPSYAKVFFWLLGIGLA   1320
211     ADFNQTAGYLIKLNEGLYSFRVRANSIAGYGDFTEVEHIKVEPPPSYAKVFFWLLGIGLA   1320
246     ADFNQTAGYLIKLNEGLYSFRVRANSIAGYGDFTEVEHIKVEPPPSYAKVFFWLLGIGLA   1320
353     ADFNQTAGYLIKLNEGLYSFRVRANSIAGYGDFTEVEHIKVEPPPSYAKVFFWLLGIGLA   1320
        ************************************************************

DAH     FLIVSLFGYVCYLHKRKVPSNDLHMNTEVNPFYASMQYIPDDWEVLRENIIQLAPLGQGS   1380
29B     FLIVSLFGYVCYLHKRKVPSNDLHMNTEVNPFYASMQYIPDDWEVLRENIIQLAPLGQGS   1380
e19     FLIVSLFGYVCYLHKRKVPSNDLHMNTEVNPFYASMQYIPDDWEVLRENIIQLAPLGQGS   1380
74e19   FLIVSLFGYVCYLHKRKVPSNDLHMNTEVNPFYASMQYIPDDWEVLRENIIQLAPLGQGS   1380
74      FLIVSLFGYVCYLHKRKVPSNDLHMNTEVNPFYASMQYIPDDWEVLRENIIQLAPLGQGS   1380
211     FLIVSLFGYVCYLHKRKVPSNDLHMNTEVNPFYASMQYIPDDWEVLRENIIQLAPLGQGS   1380
246     FLIVSLFGYVCYLHKRKVPSNDLHMNTEVNPFYASMQYIPDDWEVLRENIIQLAPLGQGS   1380
353     FLIVSLFGYVCYLHKRKVPSNDLHMNTEVNPFYASMQYIPDDWEVLRENIIQLAPLGQGS   1380
        ************************************************************
```

```
DAH       FGMVYEGILKSFPPNGVDRECAIKTVNENATDRERTNFLSEASVMKEFDTYHVVRLLGVC   1440
29B       FGMVYEGILKSFPPNGVDRECAIKTVNENATDRERTNFLSEASVMKEFDTYHVVRLLGVC   1440
e19       FGMVYEGILKSFPPNGVDRECAIKTVNENATDRERTNFLSEASVMKEFDTYHVVRLLGVC   1440
74e19     FGMVYEGILKSFPPNGVDRECAIKTVNENATDRERTNFLSEASVMKEFDTYHVVRLLGVC   1440
74        FGMVYEGILKSFPPNGVDRECAIKTVNENATDRERTNFLSEASVMKEFDTYHVVRLLGVC   1440
211       FGMVYEGILKSFPPNGVDRECAIKTVNENATDRERTNFLSEASVMKEFDTYHVVRLLGVC   1440
246       FGMMYEGILKSFPPNGVDRECAIKTVNENATDRERTNFLSEASVMKEFDTYHVVRLLGVC   1440
353       FGMVYEGILKSFPPNGVDRECAIKTVNENATDRERTNFLSEASVMKEFDTYHVVRLLGVC   1440
          ***:********************************************************

DAH       SRGQPALVVMELMKKGDLKSYLRAHRPEERDEAMMTYLNRIGVTGNVQPPTYGRIYQMAI   1500
29B       SRGQPALVVMELMKKGDLKSYLRAHRPEERDEAMMTYLNRIGVTGNVQPPTYGRIYQMAI   1500
e19       SRGQPALVVMELMKKGDLKSYLRAHRPEERDEAMMTYLNRIGVTGNVQPPTYGRIYQMAI   1500
74e19     SRGQPALVVMELMKKGDLKSYLRAHRPEERDEAMMTYLNRIGVTGNVQPPTYGRIYQMAI   1500
74        SRGQPALVVMELMKKGDLKSYLRAHRPEERDEAMMTYLNRIGVTGNVQPPTYGRIYQMAI   1500
211       SRGQPALVVMELMKKGDLKSYLRAHRPEERDEAMMTYLNRIGVTGNVQPPTYGRIYQMAI   1500
246       SRGQPALVVMELMKKGDLKSYLRAHRPEERDEAMMTYLNRIGVTGNVQPPTYGRIYQMAI   1500
353       SRGQPALVVMELMKKGDLKSYLRAHCPEERDEAMMTYLNRIGVTGNVQPPTYGRIYQMAI   1500
          ************************* **********************************

DAH       EIADGMAYLAAKKFVHRDLAARNCMVADDLTVKIGDFGMTRDIYETDYYRKGTKGLLPVR   1560
29B       EIADGMAYLAAKKFVHRDLAARNCMVADDLTVKIGDFGMTRDIYETDYYRKGTKGLLPVR   1560
e19       EIADGMAYLAAKKFVHRDLAARNCMVADDLTVKIGDFGMTRDIYETDYYRKGTKGLLPVR   1560
74e19     EIADGMAYLAAKKFVHRDLAARNCMVADDLTVKIGDFGMTRDFYETDYYRKGTKGLLPVR   1560
74        EIADGMAYLAAKKFVHRDLAARNCMVADDLTVKIGDFGMTRDFYETDYYRKGTKGLLPVR   1560
211       EIADGMAYLAAKKFVHRDLAARNCMVADDLTVKIGDFGMTRDIYETDYYRKGTKGLLPVR   1560
246       EIADGMAYLAAKKFVHRDLAARNCMVADDLTVKIGDFGMTRDIYETDYYRKGTKGLLPVR   1560
353       EIADGMAYLAAKKFVHRDLAARNCMVADDLTVKIGDFGMTRDIYETDYYRKGTKGLLPVR   1560
          ******************************************:*****************

DAH       WMPPESLRDGVYSSASDVFSFGVVLWEMATLAAQPYQGLSNEQVLRYVIDGGVMERPENC   1620
29B       WMPPESLRDGVYSSASDVFSFGVVLWEMATLAAQPYQGLSNEQVLRYVIDGGVMERPENC   1620
e19       WMPPESLRDGVYSSASDVFSFGVVLWEMATLAAQPYQGLSNEQVLRYVIDGGVMERPENC   1620
74e19     WMPPESLRDGVYSSASDVFSFGVVLWEMATLAAQPYQGLSNEQVLRYVIDGGVMERPENC   1620
74        WMPPESLRDGVYSSASDVFSFGVVLWEMATLAAQPYQGLSNEQVLRYVIDGGVMERPENC   1620
211       WMPPESLRDGVYSSASDVFSFGVVLWEMATLAAQPYQRLSNEQVLRYVIDGGVMERPENC   1620
246       WMPPESLRDGVYSSASDVFSFGVVLWEMATLAAQPYQGLSNEQVLRYVIDGGVMERPENC   1620
353       WMPPESLRDGVYSSASDVFSFGVVLWEMATLAAQPYQGLSNEQVLRYVIDGGVMERPENC   1620
          ************************************* **********************

DAH       PDFLHKLMQRCWHHRSSARPSFLDIIAYLEPQCPNSQFKEVSFYHSEAGLQHREKERKER   1680
29B       PDFLHKLMQRCWHHRSSARPSFLDIIAYLEPQCPNSQFKEVSFYHSEAGLQHREKERKER   1680
e19       PDFLHKLMQRCWHHRSSARPSFLDIIAYLEPQCPNSQFKEVSFYHSEAGLQHREKERKER   1680
74e19     PDFLHKLMQRCWHHRSSARPSFLDIIAYLEPQCPNSQFKEVSFYHSEAGLQHREKERKER   1680
74        PDFLHKLMQRCWHHRSSARPSFLDIIAYLEPQCPNSQFKEVSFYHSEAGLQHREKERKER   1680
211       PDFLHKLMQRCWHHRSSARPSFLDIIAYLEPQCPNSQFKEVSFYHSEAGLQHREKERKER   1680
246       PDFLHKLMQRCWHHRSSARPSFLDIIAYLEPQCPNSQFKEVSFYHSEAGLQHREKERKER   1680
353       PDFLHKLMQRCWHHRSSARPSFLDIIAYLEPQCPNSQFKEVSFYHSEAGLQHREKERKER   1680
          ************************************************************

DAH       NQLDAFAAVPLDQDLQDREQQEDATTPLRMGDYQQNSSLDQPPESPIAMVDDQGSHLPFS   1740
29B       NQLDAFAAVPLDQDLQDREQQEDATTPLRMGDYQQNSSLDQPPESPIAMVDDQGSHLPFS   1740
e19       NQLDAFAAVPLDQDLQDREQQEDATTPLRMGDYQQNSSLDQPPESPIAMVDDQGSHLPFS   1740
74e19     NQLDAFAAVPLDQDLQDREQQEDATTPLRMGDYQQNSSLDQPPESPIAMVDDQGSHLPFS   1740
74        NQLDAFAAVPLDQDLQDREQQEDATTPLRMGDYQQNSSLDQPPESPIAMVDDQGSHLPFS   1740
211       NQLDAFAAVPLDQDLQDREQQEDATTPLRMGDYQQNSSLDQPPESPIAMVDDQGSHLPFS   1740
246       NQLDAFAAVPLDQDLQDREQQEDATTPLRMGDYQQNSSLDQPPESPIAMVDDQGSHLPFS   1740
353       NQLDAFAAVPLDQDLQDREQQEDATTPLRMGDYQQNSSLDQPPESPIAMVDDQGSHLPFS   1740
          ************************************************************
```

```
DAH      LPSGFIASSTPDGQTVMATAFQNIPAAQGDISATYVVPDADALDGDRGYEIYDPSPKCAE   1800
29B      LPSGFIASSTPDGQTVMATAFQNIPAAQGDISATYVVPDADALDGDRGYEIYDPSPKCAE   1800
e19      LPSGFIASSTPDGQTVMATAFQNIPAAQGDISATYVVPDADALDGDRGYEIYDPSPKCAE   1800
74e19    LPSGFIASSTPDGQTVMATAFQNIPAAQGDISATYVVPDADALDGDRGYEIYDPSPKCAE   1800
74       LPSGFIASSTPDGQTVMATAFQNIPAAQGDISATYVVPDADALDGDRGYEIYDPSPKCAE   1800
211      LPSGFIASSTPDGQTVMATAFQNIPAAQGDISATYVVPDADALDGDRGYEIYDPSPKCAE   1800
246      LPSGFIASSTPDGQTVMATAFQNIPAAQGDISATYVVPDADALDGDRGYEIYDPSPKCAE   1800
353      LPSGFIASSTPDGQTVMATAFQNIPAAQGDISATYVVPDADALDGDRGYEIYDPSPKCAE   1800
         ************************************************************

DAH      LPTSRSGSTGGGKLSGEQHLLPRKGRQPTIMSSSMPDDVIGGSSLQPSTASAASSNASSH   1860
29B      LPTSRSGSTGGGKLSGEQHLLPRKGRQPTIMSSSMPDDVIGGSSLQPSTASAASSNASSH   1860
e19      LPTSRSGSTGGGKLSGEQHLLPRKGRQPTIMSSSMPDDVIGGSSLQPSTASAASSNASSH   1860
74e19    LPTSRSGSTGGGKLSGEQHLLPRKGRQPTIMSSSMPDDVIGGSSLQPSTASAASSNASSH   1860
74       LPTSRSGSTGGGKLSGEQHLLPRKGRQPTIMSSSMPDDVIGGSSLQPSTASAASSNASSH   1860
211      LPTSRSGSTGGGKLSGEQHLLPRKGRQPTIMSSSMPDDVIGGSSVQPSTASAASSNASSH   1860
246      LPTSRSGSTGGGKLSGEQHLLPRKGRQPTIMSSSMPDDVIGGSSLQPSTASAASSNASSH   1860
353      LPTSRSGSTGGGKLSGEQHLLPRKGRQPTIMSSSMPDDVIGGSSLQPSTASAASSNASSH   1860
         ********************************************:***************

DAH      TGRPSLKKTVADSVRNKANFINRHLFNHKRTGSNASHKSNASNAPSTSSNTNLTSHPVAM   1920
29B      TGRPSLKKTVADSVRNKANFINRHLFNHKRTGSNASHKSNASNAPSTSSNTNLTSHPVAM   1920
e19      TGRPSLKKTVADSVRNKANFINRHLFNHKRTGSNASHKSNASNAPSTSSNTNLTSHPVAM   1920
74e19    TGRPSLKKTVADSVRNKANFINRHLFNHKRTGSNASHKSNASNAPSTSSNTNLTSHPVAM   1920
74       TGRPSLKKTVADSVRNKANFINRHLFNHKRTGSNASHKSNASNAPSTSSNTNLTSHPVAM   1920
211      TGRPSLKKTVADSVRNKANFINRHLFNHKRTGSNASHKSNASNAPSTSSNTNLTSHPVAV   1920
246      TGRPSLKKTVADSVRNKANFINRHLFNHKRTGSNASHKSNASNAPSTSSNTNLTSHPVAM   1920
353      TGRPSLKKTVADSVRNKANFINRHLFNHKRTGSNASHKSNASNAPSTSSNTNLTSHPVAM   1920
         ***********************************************************:

DAH      GNLGTIESGGSGSAGSYTGTPRFYTPSATPGGGSGMAISDNPNYRLLDESIASEQATILT   1980
29B      GNLGTIESGGSGSAGSYTGTPRFYTPSATPGGGSGMAISDNPNYRLLDESIASEQATILT   1980
e19      GNLGTIESGGSGSAGSYTGTPRFYTPSATPGGGSGMAISDNPNYRLLDESIASEQATILT   1980
74e19    GNLGTIESGGSGSAGSYTGTPRFYTPSATPGGGSGMAISDNPNYRLLDESIASEQATILT   1980
74       GNLGTIESGGSGSAGSYTGTPRFYTPSATPGGGSGMAISDNPNYRLLDESIASEQATILT   1980
211      GNLGTIESGGSGSAGSYTGTPRFYTPSATPGGGSGMAISDNPNYRLLDESIASEQATILT   1980
246      GNLGTIESGGSGSAGSYTGTPRFYTPSATPGGGSGMAISDNPNYRLLDESIASEQATILT   1980
353      GNLGTIESGGSGSAGSYTGTPRFYTPSATPGGGSGMAISDNPNYRLLDESIASEQATILT   1980
         ************************************************************

DAH      TSSPNPNYEMMHPPTSLVSTNPNYMPMNETPVQMAGVTISHNPNYQPMQAPLNARQSQSS   2040
29B      TSSPNPNYEMMHPPTSLVSTNPNYMPMNETPVQMAGVTISHNPNYQPMQAPLNARQSQSS   2040
e19      TSSPNPNYEMMHPPTSLVSTNPNYMPMNETPVQMAGVTISHNPNYQPMQAPLNARQSQSS   2040
74e19    TSSPNPNYEMMHPPTSLVSTNPNYMPMNETPVQMAGVTISHNPNYQPMQAPLNARQSQSS   2040
74       TSSPNPNYEMMHPPTSLVSTNPNYMPMNETPVQMAGVTISHNPNYQPMQAPLNARQSQSS   2040
211      TSSPNPNYEMMHPPTSLVSTNPNYMPMNETPVQMAGVTISHNPNYQPMQAPLNARQSQSS   2040
246      TSSPNPNYEMMHPPTSLVSTNPNYMPMNETPVQMAGVTISHNPNYQPMQAPLNARQSQSS   2040
353      TSSPNPNYEMMHPPTSLVSTNPNYMPMNETPVQMAGVTISHNPNYQPMQAPLNARQSQSS   2040
         ************************************************************

DAH      SDEDNEQEEDDEDEDDDVDDEHVEHIKMERMPLSRPRQRALPSKTQPPRSRSVSQTRKSP   2100
29B      SDEDNEQEEDDEDEDDDVDDEHVEHIKMERMPLSRPRQRALPSKTQPPRSRSVSQTRKSP   2100
e19      SDEDNEQEEDDEDEDDDVDDEHVEHIKMERMPLSRPRQRALPSKTQPPRSRSVSQTRKSP   2100
74e19    SDEDNEQEEDDEDEDDDVDDEHVEHIKMERMPLSRPRQRALPSKTQPPRSRSVSQTRKSP   2100
74       SDEDNEQEEDDEDEDDDVDDEHVEHIKMERMPLSRPRQRALPSKTQPPRSRSVSQTRKSP   2100
211      SDEDNEQEEDDEDEDDDVDDEHVEHIKMERMPLSRPRQRALPSKTQPPRSRSVSQTRKSP   2100
246      SDEDNEQEEDDEDEDDDVDDEHVEHIKMERMPLSRPRQRALPSKTQPPRSRSVSQTRKSP   2100
353      SDEDNEQEEDDEDEDDDVDDEHVEHIKMERMPLSRPRQRALPSKTQPPRSRSVSQTRKSP   2100
         ************************************************************
```

```
DAH       TNPNSGIGATGAGNRSNLLKENWLRPASTPRPPPPNGFIGREA     2143
29B       TNPNSGIGATGAGNRSNLLKENWLRPASTPRPPPPNGFIGREA     2143
e19       TNPNSGIGATGAGNRSNLLKENWLRPASTPRPPPPNGFIGREA     2143
74e19     TNPNSGIGATGAGNRSNLLKENWLRPASTPRPPPPNGFIGREA     2143
74        TNPNSGIGATGAGNRSNLLKENWLRPASTPRPPPPNGFIGREA     2143
211       TNPNSGIGATGAGNRSNLLKENWLRPASTPRPPPPNGFIGREA     2143
246       TNPNSGIGATGAGNRSNLLKENWLRPASTPRPPPPNGFIGREA     2143
353       TNPNSGIGATGAGNRSNLLKENWLRPASTPRPPPPNGFIGREA     2143
          *******************************************
```

**Table S1. Survival statistics of alleles in Quantitative Complementation Test.** Male and female cohorts. Wildtype test alleles are *InR*$^{+}$ on extracted third chromosomes (NC series) sampled from the Raleigh Farmer's Market by T.F. Mackay. Mutant *InR* test alleles are archival EMS-treated third chromosomes with lesions mapped over a deficiency, reported in Fernandez (59) provided by M. Frasch. N(0): initial cohort size, *InR*$^{allele}$/ *InR*$^{E19}$ (*InR*$^{allele}$ /E19) and *InR*$^{allele}$ / TM3, *InR*$^{+}$ *Sb* (*InR*$^{allele}$ /TM3) combined. * : mutant *InR* alleles were tested in two independent blocks; data shows combined sample size and average (after block adjustment) survival parameter estimates.

Survival parameters are adjusted across blocks. In each of five periods used to construct life tables (blocks), the relative hazard coefficient $\beta$ was estimated for a universal wildtype chromosome (NC442) as NC442/E19 relative to NC442/TM3 (Table S2). To adjust for period effects among blocks, the $\beta_{NC442}$ expressed in the block was subtracted from each $\beta$ estimated for wildtype and *InR* mutants tested within the corresponding block. Proportional Hazard is the ratio of estimated mortality among compared cohorts after adjusting for their common time dependent underlying hazard $h_0(t)$, estimated as $\exp(|\beta|)$; unity indicates no overall mortality difference. **Bold** values indicate test allele decreases mortality (*InR*$^{allele}$ /TM3: *InR*$^{allele}$ /E19). Normal values indicate test allele elevates mortality (*InR*$^{allele}$ /E19: *InR*$^{allele}$ /TM3). Estimates are used to visualize the distribution of allelic effects (fold-change) among naturally segregating polymorphisms of *InR* (Figure 2, main text). Life Expectancy is observed difference between the Kaplan-Meier estimated median life span of the *InR*$^{allele}$ /E19 cohort and the *InR*$^{allele}$ /TM3 cohort, adjusted by subtracting the difference in life expectancy between NC442/E19 relative to NC442/TM3 of the block.

Block adjusted $\beta$ for each *InR* EMS allele is evaluated by *z*-test relative to the distribution of $\beta$ among NC alleles where mean value is -0.15 (both sexes) and SD is 0.44 and 0.48 for males and females respectively. Adjusting for multiple comparisons with Bonferroni correction ($p < 0.0055$), the alleles *InR*$^{74}$ and *InR*$^{211}$ in both sexes decrease mortality more than expected; *InR*$^{353}$ reduces mortality about 2.5-fold in females, but not significantly by our criteria; *InR*$^{262}$ modestly decreases mortality in both sexes, but no more than many wildtype *InR* alleles; *InR*$^{327}$ is strongly adult deleterious in both sexes.

| **Test Allele. Males** | | **Survival Parameters (block adjusted)** | | | | |
|---|---|---|---|---|---|---|
| | | Change Life Expectancy allele/E19-allele/TM3 (days) | | Proportional Hazard (allele/E19:allele/TM3) | | |
| **Wiltype** | N(0) | | $\beta$ | **(allele/TM3:allele/E19)** | | |
| NC434 | 424 | 13.8 | -0.52 | **1.68** | | |
| NC469 | 371 | 3.6 | -0.32 | **1.38** | | |
| NC361 | 429 | 0.1 | 0.04 | 1.04 | | |
| NC424 | 335 | -1.9 | 0.10 | 1.11 | | |
| NC443 | 424 | -3.7 | 0.14 | 1.15 | | |
| NC464 | 399 | 1.3 | -0.25 | **1.29** | | |
| NC391 | 414 | 5.2 | -0.71 | **2.03** | | |
| NC453 | 476 | 9.2 | -0.95 | **2.58** | | |
| NC291 | 613 | -2.1 | 0.32 | 1.37 | | |
| NC245 | 560 | 0.2 | -0.35 | **1.42** | | |
| NC297 | 650 | -9.9 | 0.47 | 1.60 | | |
| NC65 | 449 | -5.7 | 0.13 | 1.14 | | |
| NC241 | 482 | 5.1 | -0.74 | **2.10** | | |
| NC351 | 467 | 3.9 | -0.09 | 1.09 | | |
| NC325 | 575 | -8.4 | 0.50 | 1.65 | | |
| NC264 | 503 | -0.2 | -0.18 | **1.19** | | |
| NC96 | 640 | -2.1 | 0.30 | 1.35 | | |
| NC99 | 488 | 5.0 | -0.62 | **1.86** | | |
| | mean | 0.74 | -0.15 | | | |
| | SD | 5.94 | 0.44 | | ***z*-test** | |
| **Mutant** | | | | | *z* | Prob. |
| InR74 | 606* | 18.8 | -1.48 | **4.39** | -3.148 | 0.0016 |
| InR76 | 429 | -4.0 | 0.75 | 2.13 | 2.105 | 0.0353 |
| InR211 | 665* | 13.3 | -0.96 | **2.62** | -1.937 | 0.0528 |
| InR246 | 381 | 0.4 | -0.06 | **1.06** | 0.202 | 0.8402 |
| InR262 | 417 | 7.3 | -0.43 | **1.54** | -0.683 | 0.4947 |
| InR273 | 507 | -7.7 | 0.66 | 1.93 | 1.879 | 0.0603 |
| InR327 | 451 | -22.1 | 1.64 | 5.13 | 4.179 | < 0.0001 |
| InR351 | 575 | -0.9 | 0.03 | 1.03 | 0.390 | 0.6967 |
| InR353 | 882* | 5.7 | -0.41 | **1.50** | -0.624 | 0.5326 |

| **Test Allele. Females** | | **Survival Parameters (block adjusted)** | | | | |
|---|---|---|---|---|---|---|
| | | Life Expectancy allele/E19-allele/TM3 (days) | | Proportional Hazard (allele/E19:allele/TM3) | | |
| **Wiltype** | N(0) | | $\beta$ | **(allele/TM3:allele/E19)** | | |
| NC434 | 396 | 10.4 | -0.75 | **2.11** | | |
| NC469 | 377 | 8.5 | -0.32 | **1.38** | | |
| NC361 | 491 | -11.3 | 0.93 | 2.54 | | |
| NC424 | 457 | -0.6 | -0.24 | **1.27** | | |
| NC443 | 409 | -8.9 | 0.74 | 2.10 | | |
| NC464 | 451 | -11.0 | -0.28 | **1.32** | | |
| NC391 | 497 | 4.6 | -0.75 | **2.11** | | |
| NC453 | 557 | 6.1 | -0.67 | **1.95** | | |
| NC291 | 475 | -1.8 | 0.28 | 1.32 | | |
| NC245 | 517 | 3.4 | -0.21 | **1.24** | | |
| NC297 | 512 | -1.4 | 0.00 | 1.00 | | |
| NC65 | 645 | 1.5 | -0.30 | **1.34** | | |
| NC241 | 531 | 7.0 | -0.63 | **1.88** | | |
| NC351 | 550 | -0.6 | 0.44 | 1.55 | | |
| NC325 | 607 | 2.5 | -0.22 | **1.24** | | |
| NC264 | 533 | 2.4 | -0.16 | **1.18** | | |
| NC96 | 523 | 3.3 | -0.18 | **1.20** | | |
| NC99 | 464 | 3.5 | -0.35 | **1.42** | | |
| | mean | 1.0 | -0.15 | | | |
| | SD | 6.2 | 0.48 | | ***z*-test** | |
| **Mutant** | | | | | *z* | Prob. |
| InR74 | 720* | 25.1 | -1.80 | **6.03** | -3.462 | 0.0006 |
| InR76 | 551 | 3.9 | -0.49 | **1.63** | -0.719 | 0.3788 |
| InR211 | 954* | 12.2 | -1.65 | **5.21** | -3.158 | 0.0020 |
| InR246 | 438 | -7.2 | 0.29 | 1.34 | 0.920 | 0.3520 |
| InR262 | 403 | 8.2 | -0.49 | **1.63** | -0.710 | 0.5418 |
| InR273 | 558 | -1.1 | 0.13 | 1.14 | 0.590 | 0.4966 |
| InR327 | 598 | -24.3 | 1.51 | 4.52 | 3.478 | < 0.0001 |
| InR351 | 574 | -0.6 | 0.00 | 1.00 | 0.313 | 0.7040 |
| InR353 | 1075* | 9.0 | -0.90 | **2.45** | -1.573 | 0.1156 |

**Table S2.** Block survival statistics of the universal wildtype test chromosome (*InR*⁺) NC442 in Quantitative Complementation Test as NC442/E19 relative to NC442/TM3.

| | **Block** | **N(0)** | $\beta$ | **s.e.** | **Life expectancy** (difference, days) NC442/E19 – NC442/TM3 |
|---|---|---|---|---|---|
| Males | | | | | |
| NC442 | 1 | 500 | -0.258 | 0.091 | 4.9 |
| NC442 | 2 | 395 | -1.18 | 0.114 | 14 |
| NC442 | 3 | 433 | -0.504 | 0.099 | 5 |
| NC442 | 4 | 665 | -0.999 | 0.084 | 17.9 |
| NC442 | 5 | 452 | -0.825 | 0.101 | 8.1 |
| Females | | | | | |
| NC442 | 1 | 463 | -0.095 | 0.094 | 1.92 |
| NC442 | 2 | 457 | -1.279 | 0.102 | 16.5 |
| NC442 | 3 | 571 | -0.33 | 0.085 | 4.4 |
| NC442 | 4 | 424 | -0.973 | 0.106 | 14.49 |
| NC442 | 5 | 445 | -0.596 | 0.097 | 8.2 |

**Table S3. Summary life table statistics for *InR* hemizygotes and InR wildtype homozygotes.** Data of Figure 5A (Trial 1) and 5B (Trial 2) in main text, females. Wildtype *InR* included three independent accessions: $InR^{+(HR)29B}$ (29+), $InR^{+(HR)13A}$ (13+) and *w*Dah. Null *InR* alleles were derived from the accessions used to produce wildtype $InR^{+(HR)29B}$ (29+) and $InR^{+(HR)13A}$ (13+). Trials were conducted in independent periods, with contemporary cohorts of indicated genotypes.

| Genotype class | Genotype | N(0) | Median survival (d) | Median 95% c.i. | Mean survival (d) | Mean s.e. |
|---|---|---|---|---|---|---|
| **Trial 1** | | | | | | |
| **Wildtype homozygote** | | | | | | |
| | 29+/29+ | 351 | 44 | 42-44 | 41.7 | 0.52 |
| | wDah/29+ | 352 | 40 | 38-42 | 39.6 | 0.55 |
| | 29+/wDah | 361 | 44 | 44-46 | 42.2 | 0.57 |
| | wDah/wDah | 348 | 48 | 46-48 | 45.9 | 0.62 |
| **Hemizygote** | | | | | | |
| | 29+/29null | 366 | 48 | 48-50 | 46.9 | 0.52 |
| | wDah/29null | 348 | 44 | 42-46 | 44.1 | 0.59 |
| **Trial 2** | | | | | | |
| **Wildtype homozygote** | | | | | | |
| | 13+/13+ | 443 | 34 | 36-36 | 33.7 | 0.51 |
| | 13+/29+ | 451 | 36 | 36-38 | 35.1 | 0.44 |
| | 29+/13+ | 342 | 38 | 38-40 | 37.99 | 0.42 |
| | 29+/29+ | 475 | 46 | 44-46 | 43.4 | 0.45 |
| | wDah/13+ | 492 | 44 | 44-44 | 43.6 | 0.32 |
| | wDah/29+ | 476 | 40 | 40-40 | 39.7 | 0.38 |
| | wDah/wDah | 461 | 42 | 42-42 | 41.2 | 0.37 |
| **Hemizygote** | | | | | | |
| | 13+/13null | 446 | 42 | 40-42 | 39.8 | 0.54 |
| | 29+/29null | 484 | 50 | 48-50 | 48.8 | 0.41 |
| | wDah/13null | 465 | 44 | 44-46 | 44 | 0.38 |
| | wDah/29null | 467 | 42 | 40-44 | 42.3 | 0.47 |

Relative mRNA abundance of *InR* from whole body lysates of 20 females per biological replicate. Three biological replicates each genotype, each with three technical replicates. InR mRNA CT normalized to Rp49 of each sample, 2(-ddCT) estimation method. Means with s.e. and difference confidence interval, t= 3.12 , p = 0.036.

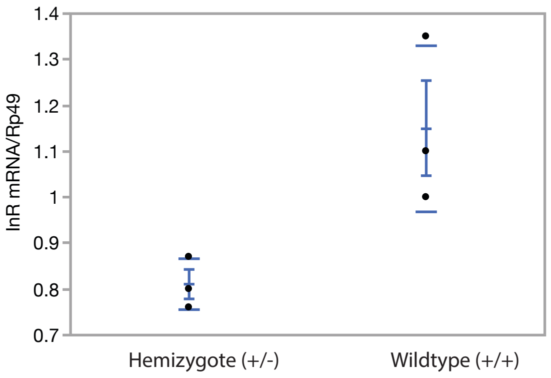

**Table S4. Summary survival statistics of homologous recombination genotypes presented in main text figure 6**. Subfigures with controls (+/+, +/-) in blue or green were contemporary cohorts. Proportional hazard analysis within each subfigure with relative mortality risk compared to wildtype or hemizygote, log-likelihood test. Accessions are independently generated stocks of indicated HR alleles.

| FIG. | HR GENOTYPE & ACCESSION | $N_0$ | MEDIAN SURVIVAL (95% C.I.) | RISK RATIO TO WILDTYPE | $P < \chi^2$ | RISK RATIO TO HEMIZYGOTE | $P < \chi^2$ |
|---|---|---|---|---|---|---|---|
| **6A** | **+/+** | **328** | **44 (42, 44)** | | | | |
| | +/*InR*(E19)-2C | 337 | 40 (40, 42) | 0.828 | 0.0154 | | |
| | +/*InR*(E19) -14B | 331 | 42 (40, 44) | 0.895 | 0.155 | | |
| | +/*InR*(E19)-22A | 351 | 48 (46, 52) | 1.495 | <0.0001 | | |
| **6B** | **+/+** | **486** | **40 (38, 40)** | | | | |
| | +/*InR*(74) | 473 | 44 (44, 44) | 0.523 | <0.0001 | | |
| | +/*InR*(211) | 487 | 38 (38, 38) | 1.06 | 0.34 | | |
| | +/*InR*(74, E19) | 468 | 40 (40, 40) | 0.810 | 0.0012 | | |
| **6C** | **+/+** | **334** | **48 (46, 48)** | | | 1.54 | <0.0001 |
| | **+/- (hemizygote)** | **346** | **50 (50, 52)** | 0.648 | <0.0001 | | |
| | +/*InR*(74) | 291 | 50 (48, 50) | 0.660 | <0.0001 | 1.02 | 0.816 |
| | +/*InR*(211) | 347 | 44 (42, 44) | 1.60 | <0.0001 | 2.47 | <0.0001 |
| | +/*InR*(246) | 332 | 50 (48, 50) | 0.743 | <0.0001 | 1.15 | 0.076 |
| | +/*InR*(353) | 347 | 60 (58, 62) | 0.215 | <0.0001 | 0.331 | <0.0001 |
| | +/*InR*(74, E19) | 343 | 48 (48, 50) | 0.881 | 0.0996 | 1.36 | <0.0001 |
| **6D** | **+/+** | **348** | **46 (44, 48)** | | | | |
| | +/*InR*(353)-8.1 | 359 | 62 (62, 64) | 0.247 | <0.0001 | | |
| | +/*InR*(353)-15.4 | 357 | 56 (54, 58) | 0.261 | <0.0001 | | |
| | +/*InR*(353)-20.1 | 362 | 58 (56,58) | 0.275 | <0.0001 | | |
| **6E** | **+/+** | **486** | **40 (38, 40)** | | | | |
| | *InR*(E19)/*InR*(74) | 446 | 54 (52, 56) | 0.179 | <0.0001 | | |
| | *InR*(E19)/*InR*(211) | 471 | 52 (50, 52) | 0.238 | <0.0001 | | |
| | *InR*(211)/*InR*(74, E19) | 456 | 48 (46, 48) | 0.350 | <0.0001 | | |
| | *InR*(74)/*InR*(211) | 350 | 46 (46, 48) | 0.334 | <0.0001 | | |
| **6F** | **+/+** | **334** | **48 (46, 48)** | | | | |
| | **+/- (hemizygote)** | **346** | **50 (50, 52)** | | | | |
| | *InR*(E19)/*InR*(74) | 324 | 62 (62, 64) | 0.138 | <0.0001 | 0.215 | <0.0001 |
| | *InR*(E19)/*InR*(211) | 351 | 58 (56, 58) | 0.259 | <0.0001 | 0.405 | <0.0001 |
| | *InR*(74)/*InR*(211) | 339 | 58 (58, 60) | 0.234 | <0.0001 | 0.366 | <0.0001 |
| **6G** | **+/+** | **334** | **48 (46, 48)** | | | | |
| | **+/- (hemizygote)** | **346** | **50 (50, 52)** | | | | |
| | *InR*(E19)/*InR*(353) | 330 | 62 (58, 66) | 0.154 | <0.0001 | 0.216 | <0.0001 |
| | *InR*(E19)/*InR*(246) | 320 | 46 (44, 48) | 0.75 | 0.0004 | 1.05 | 0.529 |
| | *InR*(74)/*InR*(353) | 348 | 70 (68, 70) | 0.142 | <0.0001 | 0.199 | <0.0001 |
| **6H** | **+/+** | **359** | **46 (44, 48)** | | | | |
| | *InR*(E19)/*InR*(246) | 347 | 48 (46, 50) | 0.679 | <0.0001 | | |
| | *InR*(E19)/*InR*(353) | 360 | 61 (60,64) | 0.213 | <0.0001 | | |
| | *InR*(74)/*InR*(353) | 354 | 67 (66,68) | 0.189 | <0.0001 | | |
